# An On-Demand Nanodisc Platform for Reconstitution of Functional Membrane Proteins into Model and Living Membranes

**DOI:** 10.64898/2026.08.17.745191

**Authors:** Long-Kai Chen, Yun-Shan Wang, Wei-Hsuan Chang, Che-Kai Lin, Pei-Tzu Huang, Ming-Chang Yu, Wen-Xin Liu, Tzu-Ting Huang, Chien-Yu Ko, Ru-Hsuan Bai, Sheng-Kai Wang, Yun-Wei Chiang, Chun-Wei Lin

## Abstract

Membrane proteins are central to transport, signaling, and pharmacological regulation, yet their direct functional reconstitution into defined membrane environments remains technically challenging. Detergent-based workflows have enabled major advances in membrane protein research, but some applications require complementary strategies that better preserve native-like lipid environments. Cell-based expression approaches, meanwhile, require long incubation times and suffer from cell-type-dependent variability. Here, we establish nanodiscs as modular carriers for the rapid delivery of both lipids and full-length membrane proteins into model and cellular membranes. Using supported lipid bilayers, we first show that membrane scaffold protein (MSP) nanodiscs mediate efficient lipid transfer within minutes, with fluorescence recovery after photobleaching confirming lateral mobility of the delivered lipids. We then extend this strategy to the bacterial calcium channel BsYetJ, achieving concentration-dependent protein incorporation and single-molecule diffusion within supported lipid bilayers. Importantly, BsYetJ-loaded nanodiscs enable direct reconstitution of functional channels into intact mammalian plasma membranes across multiple cell lines. Calcium imaging demonstrates robust BsYetJ-mediated calcium influx, confirming that the delivered channel retains ion-conductive activity after transfer into heterologous cellular membranes. Crucially, this nanodisc-mediated delivery bypasses the variable trafficking pathways inherent to different host systems, allowing for the direct reconstitution of membrane proteins into target membranes while preserving their functional activity. Furthermore, unlike MSP nanodiscs, styrene-maleic acid (SMA) nanodiscs can directly capture membrane proteins from native cell membranes. This capability makes them particularly well-suited for studying complex and challenging membrane proteins. Therefore, we further generalize this platform using SMA nanodiscs . We demonstrate that, similar to MSP nanodiscs, SMA nanodiscs can efficiently deliver lipid cargo to supported bilayers and mammalian cells. By directly capturing full-length dopamine D2 receptor from cellular membranes and transferring it into naive target cells, we achieve functional GPCR reconstitution, as validated by specific binding of a custom fluorescent agonist. Together, these results demonstrate that nanodiscs can serve not only as stabilizing membrane mimetics but also as active delivery vehicles for on-demand membrane protein reconstitution. This approach provides a rapid and broadly applicable platform for interrogating ion channels, GPCRs, and other challenging pharmacological targets in user-defined membrane environments.

## Introduction

Membrane proteins play critical roles in a vast array of biological processes, functioning as cellular carriers for transporting substances (1–3), acting as enzymes to catalyze reactions (4–6), and serving as receptors for signal transduction (7–9). Consequently, structural or functional abnormalities in these proteins are frequently the underlying cause of various diseases. Because they constitute the majority of current pharmacological targets (10), studying membrane proteins is of paramount importance to the development of novel therapeutics.

Despite their importance, *in vitro* reconstitution and characterization of membrane proteins present significant challenges. Membrane proteins are highly hydrophobic and naturally tend to aggregate and precipitate in aqueous solutions. The most common traditional method for purification relies on the use of detergents (11–13). Due to their amphipathic nature, detergents dissolve the native cell membrane and surround the proteins to form soluble micelles, enabling downstream research. While detergents are often effective at initially extracting and stabilizing the overall structure of membrane proteins, they inevitably remove the proteins from their native lipid bilayer environment (14, 15). Although detergent micelles successfully maintain protein solubility, a true lipid bilayer is still fundamentally required to accurately investigate physiological processes, such as protein-protein interactions, lipid-protein dynamics, and complex functional assemblies. Therefore, maintaining membrane proteins within a stabilizing lipid bilayer environment remains absolutely crucial for studying them effectively.

Nanodiscs have emerged as a highly successful alternative to overcome detergent-based limitations by preserving proteins in a functional, native-like lipid bilayer (16–18). Scaffolding strategies have evolved to predominantly utilize membrane scaffold proteins (MSPs) and styrene-maleic acid lipid particles (SMALPs) (19, 20). MSP nanodiscs encapsulate phospholipids within an amphipathic MSP belt—engineered from human apolipoprotein A-I (Apo-A1) (21)—whose length directly dictates the disk’s physical diameter (22). Because Apo-A1 and its High-Density Lipoprotein (HDL) precursors naturally mediate physiological reverse cholesterol transport (23), MSP nanodiscs possess remarkably high biocompatibility, facilitating their widespread application as drug delivery carriers and imaging platforms (24–27). Complementing this approach, SMALPs utilize a synthetic SMA copolymer that directly intercalates into biological membranes, solubilizing and capturing membrane proteins along with their endogenous lipids to form stable, detergent-free particles (28–30). Together, these versatile scaffolding strategies efficiently incorporate challenging targets—including cytochrome P450s (31, 32), bacteriorhodopsin (11, 33, 34), and GPCRs (29, 35)—while ensuring aqueous solubility, making them fully compatible with diverse high-resolution structural techniques such as cryo-EM (36, 37), NMR (31, 38), MS (37), SAXS (39), and ESR (40–42). Beyond structural characterization, nanodiscs also provide a useful platform for single-molecule fluorescence measurements of membrane proteins. For example, native nanodiscs combined with TIRF-based single-molecule photobleaching analysis have enabled quantitative determination of membrane-protein oligomeric states while preserving their proximal native membrane environment (43).

The use of nanodiscs as membrane transport tools has attracted increasing interest in recent years. Several studies have shown that nanodiscs can deliver lipids to diverse membrane systems, including SUVs, GUVs, and SLBs, as well as to the plasma membranes of living cells (44–46). Nanodiscs have also been used to transfer membrane proteins into cellular membranes while retaining measurable biological activity, highlighting their potential as delivery vehicles for membrane protein reconstitution (47–50). However, the systematic use of nanodisc-mediated delivery across different membrane environments remains relatively limited, and the molecular behavior of membrane proteins following delivery is not yet fully understood. Building on our previous efforts in this direction, we sought to establish a broader framework for nanodisc-mediated membrane transfer (40, 41, 51, 52). In the present study, we investigate nanodiscs as active carriers for the rapid reconstitution of lipids and full-length membrane proteins into both model and living membranes, with particular emphasis on their incorporation, mobility, localization, and functional competence after delivery.

We present a systematic investigation establishing the nanodisc as a broadly applicable carrier for the direct transport of both lipids and membrane proteins across disparate membrane systems. Focusing on complex structural targets, we successfully achieved the non-native functional reconstitution of two full-length, multi-pass transmembrane proteins: the channel protein BsYetJ and the GPCR Dopamine Receptor D2 (D2R) (53, 54). We first validated this transport mechanism using the supported lipid bilayers (SLBs) as a model membrane system, confirming both lipid integration and the structural reconstitution of BsYetJ. We then extended this delivery system to living biological models. To rigorously evaluate the platform’s robustness, we successfully utilized nanodiscs to reconstitute bacterially expressed BsYetJ into the intact plasma membranes of a diverse panel of mammalian cell lines, deliberately selected to range from easily manipulated to inherently difficult-to-transfect models. Notably, across these diverse cellular environments, we observed a distinct, BsYetJ-mediated increase in intracellular calcium ion flux. Crucially, this direct delivery strategy bypasses the variable trafficking pathways inherent to different host systems. This approach allows for the direct reconstitution of membrane proteins into target membranes to seamlessly execute their functionality. Building upon these findings, we synthesized and deployed styrene-maleic acid (SMA) polymer nanodiscs. Because SMA nanodiscs can directly capture membrane proteins from native cell membranes, they are well-suited for studying complex and challenging membrane targets. Leveraging this capability, we successfully captured and functionally reconstituted full-length D2R into naive mammalian cell lines. Using a custom-synthesized fluorescent agonist, we rigorously validated the functional integrity of the delivered D2R, confirming that the receptor firmly maintains its native ligand-binding conformation post-transfer. Collectively, these results demonstrate the successful functional integration of a fully active channel protein into the plasma membrane of living cells via a nanodisc-mediated approach, thereby verifying its capability to mediate transmembrane transport while simultaneously providing a versatile platform for the in vitro interrogation of complex GPCRs.

## Results and Discussion

### Nanodisc-Mediated Lipid Integration into Model Membrane Systems

As a foundational component of this research, we have successfully established a robust nanodisc-mediated delivery platform utilizing nanodiscs as vehicles to incorporate lipid molecules into SLBs. Given that a central objective of this project is the *in vitro* reconstitution of both the dopamine D2 receptor (D2R) signaling mechanisms and the functional calcium conductance of the bacterial channel BsYetJ, we focused on utilizing nanodiscs for the rapid, real-time modulation of lipid composition within pre-formed membrane systems. As illustrated in the schematic in Figure 1A, nanodisc-mediated lipid incorporation into the SLB was efficiently achieved at room temperature within approximately one minute. We further characterized the concentration-dependence of this process. As shown in Figure 1B, increasing the nanodisc concentration resulted in a higher density of delivered lipids, evidenced by the enhanced fluorescence intensity of the incorporated probes within the SLB. The rapid kinetics of this delivery were confirmed in Figures 1C-D, where substantial lipid accumulation was observed in less than 60 seconds.

**Figure 1.**
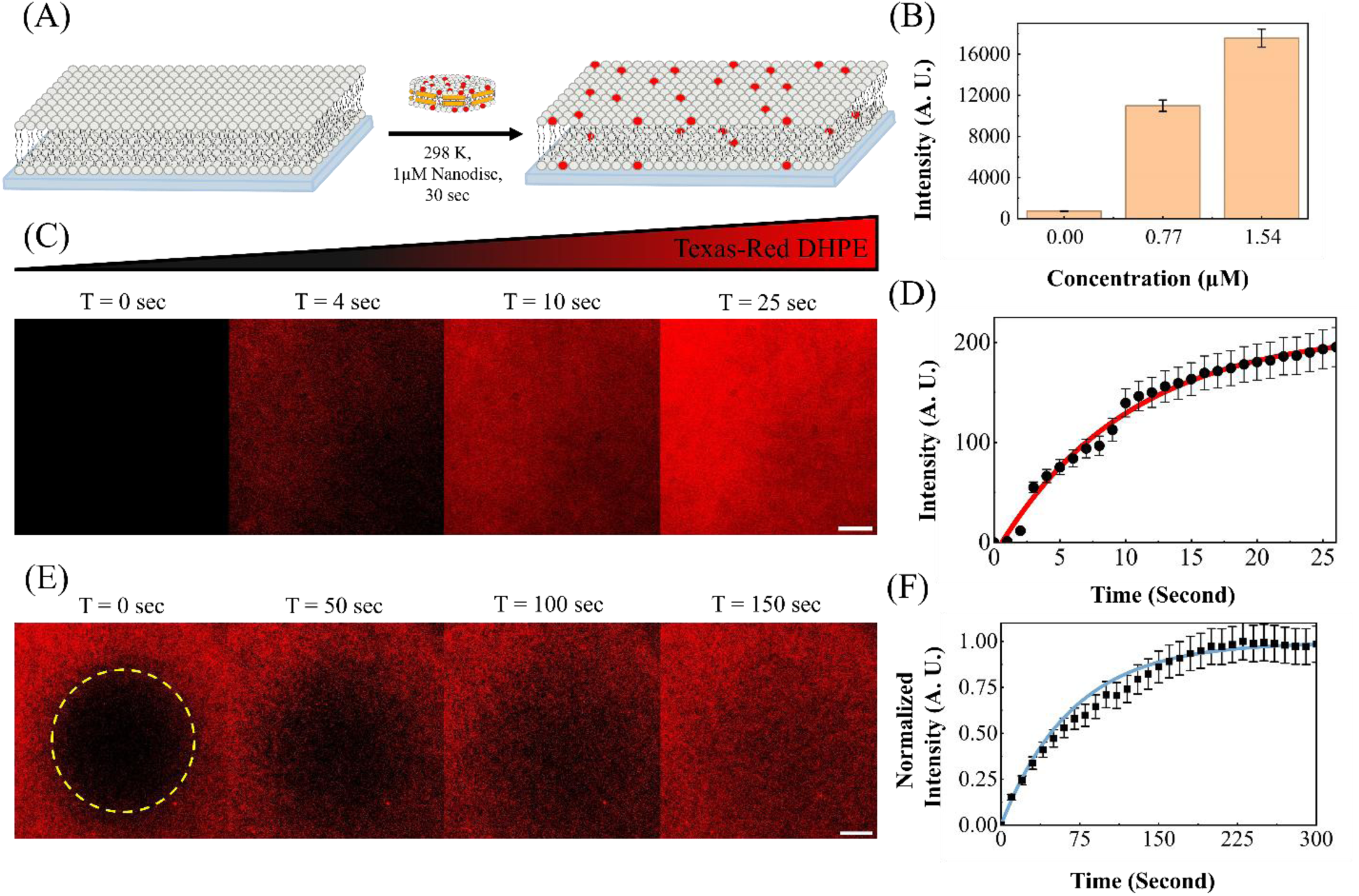
Nanodisc-mediated lipid integration for the real-time modulation of supported lipid bilayer (SLB) composition. (A) Schematic illustration of the delivery process: the introduction of Membrane Scaffold Protein (MSP) nanodiscs loaded with Texas Red-DHPE enables the direct incorporation of fluorescent lipids into an initially non-fluorescent SLB, resulting in observable fluorescence from the membrane. (B) Concentration-dependent lipid integration, demonstrating that the fluorescence intensity of the SLB increases correspondingly with the concentration of the applied nanodisc solution (up to 1.54 mM). (C) Total Internal Reflection Fluorescence (TIRF) images showing the rapid increase in SLB fluorescence intensity within 25 seconds upon the addition of 1 µM Texas Red-DHPE-loaded nanodiscs. (D) Time-course quantification of the fluorescence intensity derived from the TIRF images in (C). (E) FRAP experiment demonstrating that the fluorescence within the photobleached region (yellow dashed circle) recovers within 3 minutes, confirming the two-dimensional fluidity of the integrated lipids. (F) The corresponding fluorescence recovery curve plotted as a function of time for the experiment shown in (E).

To verify that the delivered lipids were functionally integrated into the SLB framework rather than merely adsorbed, we performed Fluorescence Recovery After Photobleaching (FRAP) experiments (Figures 1C–F). While the nanodisc-mediated delivery of lipid cargo is completed within approximately one minute, the observed fluorescence recovery over a five-minute period demonstrates that the delivered lipids undergo two-dimensional Brownian motion, thereby preserving the lateral mobility and fluidity characteristic of the SLB system (55). This robust recovery behavior further confirms that the integrated lipids have become an intrinsic component of the bilayer, rather than transiently associated with the surface.

To further determine whether the delivered lipids were incorporated into both leaflets of the SLB, we performed Cu(II)-mediated fluorescence quenching experiments (Figure S1). After nanodisc-mediated delivery of Texas Red-labeled lipids, Cu²⁺ ions were introduced into the bulk solution as a membrane-impermeable quenching agent that selectively accesses fluorophores exposed on the upper leaflet of the SLB. The addition of Cu²⁺ caused a rapid decrease in fluorescence intensity by approximately 50% within 10 s, indicating selective quenching of the solvent-exposed leaflet. Because the remaining fluorescence arises from lipids protected in the lower leaflet, this result demonstrates that nanodisc-delivered lipids are distributed across both leaflets of the bilayer rather than being restricted to the outer surface. Together with the FRAP analysis, these quenching experiments confirm that nanodisc-mediated delivery results in bona fide transbilayer lipid integration while preserving bilayer fluidity.

As an additional mechanistic validation, we examined the post-delivery localization and mobility of the nanodisc scaffold during lipid transfer using single-molecule TIRF imaging of Alexa Fluor 647-labeled MSPs (Figure S2). The MSP was engineered with a C-terminal cysteine residue to enable fluorophore conjugation through maleimide chemistry. Upon addition of labeled MSP nanodiscs to the SLB, the number of detected MSP molecules rapidly increased over time, indicating that MSPs accumulate on or near the bilayer during the lipid delivery process. Quantitative analysis further showed that this accumulation was concentration dependent, with higher nanodisc concentrations producing a larger number of MSP molecules within the imaging field. However, single-molecule tracking revealed that most MSP molecules displayed limited displacement after delivery. This relatively immobile behavior suggests that, after lipid release, MSPs traverse the SLB and become sterically trapped on the underlying glass substrate rather than remaining as freely diffusing components of the membrane. These observations support a delivery mechanism in which lipids are incorporated into the fluid bilayer, whereas the MSP scaffold is largely excluded from lateral membrane diffusion after the delivery process. To confirm the versatility of this delivery mechanism beyond solid-supported architectures, we further demonstrated that MSP nanodiscs efficiently integrate lipid cargoes into the free-standing, three-dimensional bilayers of giant unilamellar vesicles (GUVs) (Figure S3).

This methodology provides an alternative route to lipid transfer that avoids several biophysical constraints associated with membrane fusion, including hydration repulsion, the high activation energy required for stalk intermediate formation, and the complex mechanics of merging distinct lipid monolayers (56–58). By reducing the influence of these barriers, the nanodisc-mediated approach offers an efficient means of transferring lipids between distinct membrane platforms and establishes a foundation for the subsequent delivery of functional membrane proteins.

### Nanodisc-Mediated Delivery of Functional Membrane Proteins into Supported Lipid Bilayers

Building upon our initial findings, we further extended the nanodisc-mediated delivery platform to incorporate the membrane protein BsYetJ into SLBs. BsYetJ is a bacterial multi-pass transmembrane channel protein responsible for mediating calcium ion flux, making it an ideal complex target to validate our functional reconstitution platform. The schematic illustration of the nanodisc-mediated delivery process is presented in Figure 2A. By maintaining experimental conditions consistent with those optimized in Figure 1, we demonstrate the versatility and robustness of this methodology. To characterize the integrated proteins at the molecular level, BsYetJ was fluorescently labeled with Alexa Fluor 647 and visualized using high-sensitivity EMCCD imaging, which allowed for the direct detection of individual protein molecules. As shown in Figures 2B and 2C, the surface density of BsYetJ molecules integrated into the SLB was dependent on the nanodisc concentration, indicating a controlled and predictable delivery mechanism.

**Figure 2.**
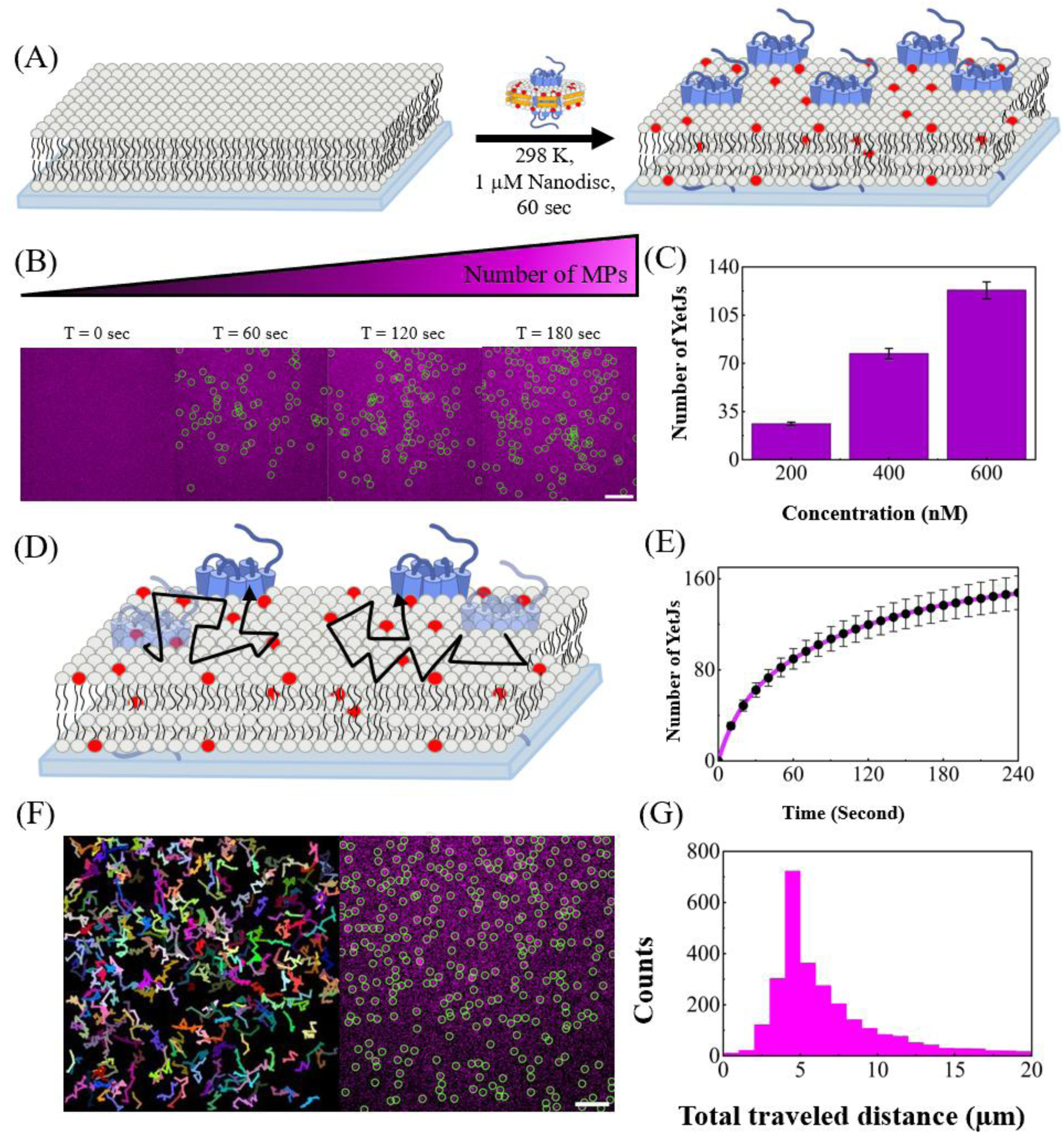
MSP nanodisc-mediated delivery and single-molecule dynamics of BsYetJ within supported lipid bilayers. (A) Schematic illustration of the nanodisc-mediated delivery of BsYetJ into the SLB. (B) Single-molecule fluorescence images of Alexa Fluor 647-labeled BsYetJ during the delivery process following the addition of 1 µM nanodiscs. The surface density of Alexa Fluor 647-labeled BsYetJ (highlighted by green circles) in the snapshots at T = 0, 60, 120, and 180 s increases rapidly upon nanodisc addition, reaching a plateau within 3 minutes (see also part E) Scale bar: 10 µm. (C) Concentration-dependent delivery of BsYetJ, demonstrating that the surface density of individual proteins integrated into the SLB at T = 240 s increases proportionally with the concentration of the applied nanodisc solution. The actual number of BsYetJ molecules on the SLB is corrected for the degree of labeling (25%) from the sample preparation. MPs: membrane proteins. Scale bar: 10 µm. (D) Schematic representation depicting the two-dimensional Brownian motion of the delivered BsYetJ, alongside co-delivered Texas Red-DHPE lipids (red dots), within the fluid SLB environment. (E) Time-course quantification of the number of delivered BsYetJ molecules, derived from the imaging data shown in (C). (F) Representative single-particle tracking trajectories (left) corresponding to the single-molecule image (right), illustrating the lateral mobility of individual BsYetJ proteins. Scale bar: 10 µm. (G) Histogram of the total traveled distances derived from the single-molecule trajectories shown in (F).

To improve membrane protein delivery efficiency, we also evaluated an indirect loading strategy based on His-tag-mediated nanodisc recruitment (Figure S4). In this approach, the polyhistidine tag on the membrane scaffold protein, originally introduced for purification, was repurposed to transiently tether nanodiscs to the SLB surface. This physical recruitment increases the local proximity between nanodiscs and the bilayer, thereby enhancing the probability of successful BsYetJ transfer. Following delivery, imidazole-mediated washing was used to remove non-integrated, His-tagged nanodiscs, allowing the remaining fluorescence signal to be attributed to BsYetJ molecules embedded within the SLB.

The insertion of membrane proteins and the concomitant delivery of lipids occurred rapidly, reaching completion within minutes (Figures 2E). Following delivery, the BsYetJ molecules exhibited two-dimensional Brownian motion, characteristic of membrane proteins in a fluid lipid environment (Figure 2D). We further performed single-particle tracking to map the trajectories of individual BsYetJ molecules (Figure 2F) and analyzed the resulting displacement distributions to quantify their lateral mobility (Figure 2G). These results establish a critical technical foundation for the *in vitro* reconstitution of complex membrane proteins, such as the D2 receptor (D2R), representing a significant step toward the overarching goals of this study.

We next examined the insertion topology of nanodisc-delivered BsYetJ using a TCEP-mediated fluorescence quenching assay (Figure S5). As a validation of the assay, mScarlet was first anchored to the upper leaflet of the SLB through maleimide chemistry. Upon addition of TCEP, the mScarlet fluorescence was nearly completely quenched within 10 s, confirming that TCEP can efficiently quench solvent-exposed mScarlet while remaining membrane-impermeable under these conditions.

We then applied this approach to C-terminally mScarlet-tagged BsYetJ delivered into SLBs by MSP nanodiscs. After TCEP addition, approximately half of the BsYetJ-mScarlet fluorescence signal was quenched, whereas the remaining population was protected from quenching. This near 50% reduction indicates that nanodisc-mediated delivery produces both insertion orientations of BsYetJ-mScarlet at roughly comparable populations within the SLB. Therefore, in addition to confirming successful membrane incorporation and lateral mobility, these topology experiments demonstrate that the delivered BsYetJ spans the bilayer and adopts mixed transmembrane orientations after reconstitution.

### Nanodisc-Mediated Reconstitution of Functional Membrane Proteins into Mammalian Cell Membranes

By leveraging the nanodisc-mediated membrane protein delivery technology developed in this study, we successfully reconstituted the bacterial calcium channel protein, BsYetJ, into the plasma membranes of several adherent mammalian cell lines. Traditionally, the investigation of membrane proteins relies on live-cell experiments where the target protein is introduced via transient transfection. However, this conventional approach presents several critical limitations. First, even rapid transient transfection protocols necessitate a prolonged incubation period—typically exceeding 8 hours—to accumulate a sufficient density of membrane proteins for reliable analysis. Furthermore, during this extended expression phase, the host cell is continuously exposed to the newly synthesized and accumulating proteins. This prolonged exposure can perturb cellular homeostasis and trigger unintended stress responses, thereby precluding the establishment of a truly independent experimental baseline. (59) Additionally, the efficiency of chemical transfection is highly variable and heavily dependent on the inherent properties of the specific cell line, which introduces significant experimental discrepancy. (60) By bypassing these bottlenecks, our methodology provides a rapid and uniform insertion of purified proteins. As will be detailed in the subsequent functional assays, the verification of ion conductance in these channel proteins establishes the technical feasibility of the *in vitro* reconstitution of full-length, multi-pass transmembrane proteins within biological systems.

As a foundational proof of concept, Figure S6 demonstrates the efficient nanodisc-mediated delivery of cargo lipids alone into target membranes. Following this successful lipid-only validation, we next investigated the simultaneous delivery of both the functional BsYetJ protein and its associated lipid molecules into the plasma membranes of various adherent mammalian cells (Figure 3A). Upon delivery, the membrane proteins are inserted into the target cell membrane, which simultaneously exhibits fluorescence from the co-incorporated fluorescently labeled lipids. Figure 3B-D displays the results for three distinct mammalian adherent cell lines—HEK, COS7, and OVCAR3—listed in descending order of their inherent susceptibility to chemical DNA transfection. (60, 61) Despite these variations, our nanodisc-mediated platform achieved efficient protein and lipid delivery across all three cell lines. Notably, epifluorescence tracking of dual-labeled nanodiscs revealed that while the cargo lipids efficiently integrate into the plasma membrane, the MSP scaffold dissociates and localizes predominantly within the cytosol, corroborating the delivery mechanism previously observed in our SLB models (Figure S7).

**Figure 3.**
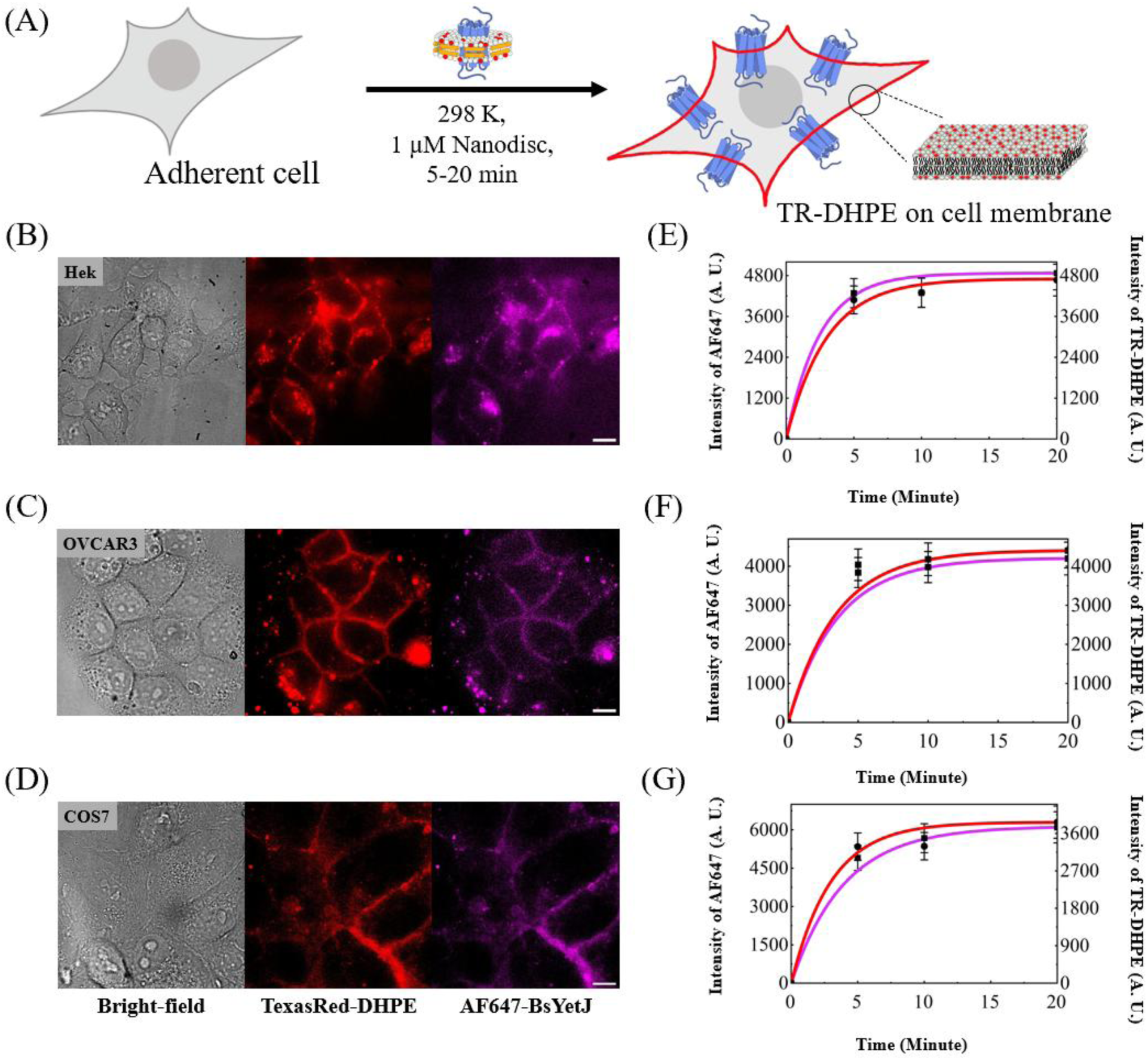
Rapid and versatile nanodisc-mediated delivery of functional membrane proteins into the plasma membrane of diverse mammalian cell lines. (A) Schematic illustration depicting the simultaneous incorporation of nanodisc-associated lipids (Texas Red-DHPE) and membrane proteins (Alexa Fluor 647-labeled BsYetJ) into the plasma membrane of naïve cells, resulting in observable dual fluorescence post-delivery. (B-D) Representative bright-field and epifluorescence images of the treated cells. The fluorescence signals clearly localize to the cellular plasma membrane, demonstrating successful integration. Quantitative Pearson correlation analysis shown in Figure S9 confirms strong spatial colocalization between the delivered lipids and BsYetJ (*r* = 0.711 for Hek, 0.721 for OVCAR3 and 0.676 for COS7). Scale bar: 10 µm. (E-G) The corresponding time-course quantification reveals the rapid membrane accumulation of both membrane proteins and lipids, with the fluorescence intensities of both Texas Red-DHPE (red) and Alexa Fluor 647-labeled BsYetJ (magenta) reaching a plateau within 10 minutes.

We also evaluated the delivery efficiency across the broader cell population and examined whether nanodisc treatment affected cell viability (Figure S8). Low-magnification epifluorescence imaging showed that nearly all cells exhibited membrane-associated Texas Red-DHPE fluorescence following treatment with 1 μM MSP nanodiscs, corresponding to an approximate 100% lipid delivery efficiency under the tested conditions. Importantly, Trypan Blue exclusion assays performed after nanodisc treatment showed negligible intracellular staining, while the cells remained adherent to the coverslip. These observations indicate that MSP nanodisc-mediated delivery maintained near-complete cell viability under the experimental conditions used. Together, these results support the high delivery efficiency of the nanodisc platform across individual cells while showing minimal acute effects on cell viability.

Distinct fluorescence from both the membrane proteins and lipids was observed within approximately five minutes of adding the BsYetJ-loaded nanodiscs to the culture medium. For each cell line, the images sequentially present bright-field transmission, fluorescent lipid (Texas Red-DHPE) signals, and Alexa Fluor 647-labeled BsYetJ fluorescence. The corresponding plots in Figures 3E-G track the lipid fluorescence intensity over time at the cellular membrane, confirming that the entire delivery process is completed within approximately five minutes. Consequently, this methodology provides a highly effective tool for the instantaneous introduction of membrane proteins into the plasma membrane, ensuring protein consistency while allowing any subsequent protein-induced cellular changes to be monitored in real time.

### Functional Characterization of Reconstituted Bacterial Ion Channels in Mammalian Plasma Membranes

To verify that BsYetJ channel proteins delivered via nanodiscs retain their native functionality, we assessed their ability to facilitate calcium ion conductance across the membrane interface. Building upon the successful delivery demonstrated above, we performed the challenging task of reconstituting these bacterial channels into the complex plasma membranes of various adherent mammalian cell lines. We employed the intracellular calcium sensor Fluo-8AM to detect changes in cytosolic concentrations. Functional reconstitution was confirmed by a robust increase in Fluo-8AM fluorescence, triggered by the influx of calcium through active BsYetJ channels.

Figure 4A provides a schematic representation of this bacterial-to-mammalian reconstitution and functional validation process. As shown in Figures 4B-D, fluorescence imaging experiments were conducted across multiple mammalian cell lines. The panels display bright-field transmission images alongside time-lapse Fluo-8AM fluorescence frames acquired at 0, 30, and 60 minutes. In all tested cell lines, the introduction of extracellular calcium (2 mM) established a gradient that resulted in a significant increase in fluorescence, indicating successful calcium influx mediated by BsYetJ. The corresponding intensity-versus-time plots in Figure 4 E-G represent statistical data averaged from a large number of cells, confirming that BsYetJ maintains its native conformation and ion-conductive properties upon nanodisc-mediated reconstitution. Notably, this nanodisc-mediated approach ensures consistent reconstitution efficiency across different cell lines, effectively overcoming the cell-type-dependent variability typically associated with traditional chemical transient transfection. To generate the robust statistical data presented in Figures 4E–G, Figure S10 presents the high-throughput version of the imaging assay, utilizing large field-of-view epifluorescence microscopy to simultaneously capture a greater population of cells.

**Figure 4.**
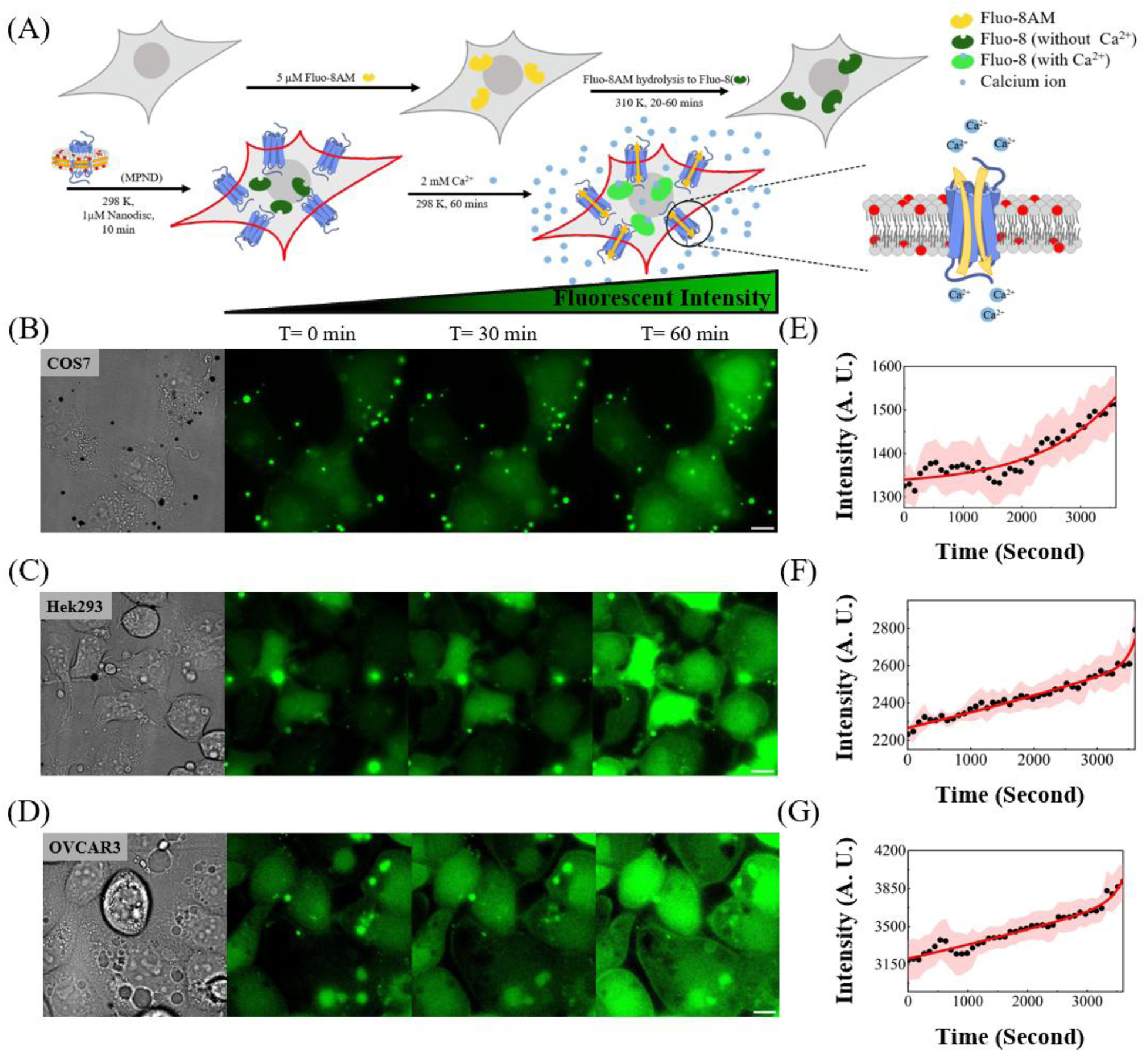
Functional validation of nanodisc-mediated BsYetJ calcium conductance in diverse mammalian cell lines. (A) Schematic illustration of the intracellular calcium imaging assay. The calcium indicator Fluo-8 AM is introduced to the cells, followed by the nanodisc-mediated delivery and integration of both full-length BsYetJ and co-incorporated fluorescent lipids into the cellular plasma membrane. Upon the addition of calcium ions to the extracellular medium, influx through the reconstituted channels is captured via chelation by the hydrolyzed indicator Fluo-8, resulting in a robust increase in fluorescence intensity. (B–D) Representative bright-field transmission images (left) and corresponding time-lapse epifluorescence frames of Fluo-8 at t = 0, 30, and 60 minutes (right) tracking calcium influx across different mammalian cell lines. (E–G) Time-course fluorescence intensity traces, demonstrating a time-dependent increase in cytosolic calcium levels mediated by the delivered channels. To generate the robust statistical data presented in these panels, large field-of-view epifluorescence microscopy was utilized as a high-throughput imaging assay to average data from multiple cells across various imaging fields (detailed in Figure S10), confirming that BsYetJ preserves its native conformation and ion-conductive properties upon nanodisc-mediated reconstitution.

Figures S11 and S12 detail the critical control experiments for the calcium influx assays presented in Figure 4. As shown in Figure S11, in the absence of nanodiscs, we observed only a progressive decrease in Fluo-8AM fluorescence caused by photobleaching during excitation. Furthermore, the high-throughput experimental data presented in Figure S12 provide robust statistical quantification for the fluorescence traces detailed in Figures S11E–G. Similarly, Figures S13 and S14 represent a second set of controls utilizing nanodiscs that were not pre-loaded with BsYetJ (empty nanodiscs). Despite the presence of these nanodiscs, the absence of the channel protein prevented calcium influx, yielding fluorescence decay profiles similar to those observed in the untreated controls. Figure S14 presents the large field-of-view epifluorescence images utilized to perform the statistical analysis of fluorescence intensity over time for these empty-nanodisc-treated cells, with the resulting traces across different cell lines detailed in Figures S13E–G. This workflow confirms that the observed fluorescence trends are attributable to photobleaching rather than ion channel activity. By bypassing complex cellular trafficking pathways, this versatile platform enables the efficient, cross-species reconstitution of bacterial proteins into mammalian membranes. Furthermore, direct validation of calcium conductance confirms that these full-length, multi-pass channels preserve their native functionality, demonstrating the overall robustness of the methodology for *in vitro* studies.

Intracellular calcium sensing with Fluo-8AM typically requires esterase-mediated hydrolysis to generate the active, carboxyl-containing form, Fluo-8. To facilitate direct comparison with results from other research groups who utilize detergent-mediated membrane permeabilization for the direct delivery of Fluo-8, we performed a parallel series of validation experiments, as summarized in Figure S15-17. Figure S16 and Figure S17 serve as negative control groups in the absence of BsYetJ. The latter specifically accounts for the potential effects of BsYetJ-free nanodiscs (empty nanodisc). In both control scenarios, the fluorescence intensity of the internalized Fluo-8 exhibited a progressive decline over time, consistent with excitation-induced photobleaching in Figures S16C and S17C. In contrast, Figure S15A illustrates the successful reconstitution of BsYetJ into HEK293 cell membranes using BsYetJ-loaded nanodiscs. The resulting calcium ion conductance through the integrated channels led to a robust and time-dependent increase in Fluo-8 fluorescence (Figure S15C). The corresponding fluorescence intensity traces, averaged from multiple cells across various imaging fields, are presented to the right of the images, further confirming the functional integrity and ion-conductive capacity of the reconstituted BsYetJ

### Establishing SMA Nanodiscs as Efficient Delivery Vehicles for Model and Cellular Membranes

Following the successful validation of nanodisc-mediated membrane protein reconstitution using BsYetJ, we extended our platform to incorporate polymer-type nanodiscs. This transition to styrene-maleic acid (SMA) polymer nanodiscs represents a critical prerequisite for the direct capture of the dopamine D2 receptor (D2R) from cellular membranes for subsequent *in vitro* reconstitution. While our initial characterization of membrane protein reconstitution and calcium ion conductance primarily utilized MSP-type nanodiscs, the study of D2R—a complex protein implicated in neurological and motor disorders—demands more direct reconstitution tools.

We successfully prepared SMA-Type nanodiscs by reacting SMA polymers with small unilamellar vesicles (SUVs), followed by purification via size-exclusion chromatography. We subsequently evaluated the efficacy of SMA nanodiscs as delivery vehicles. Figure 5A provides a schematic of SMA nanodiscs, loaded with Texas Red-DHPE, incorporating their lipid cargo into SLBs. TIRF imaging reveals a rapid and uniform enhancement in fluorescence intensity (Figure 5B), demonstrating that significant lipid integration is achieved within one minute at room temperature. FRAP experiments (Figure 5C) further confirmed the successful integration and lateral mobility of the delivered lipids. The rapid fluorescence recovery within the photobleached region (indicated by the dashed circle) demonstrates efficient two-dimensional Brownian motion and exchange with the surrounding bilayer. To confirm the versatility of this delivery mechanism beyond planar, solid-supported architectures, we further demonstrated that SMA nanodiscs efficiently integrate lipid cargoes into the free-standing, three-dimensional bilayers of giant unilamellar vesicles (GUVs) (Figure S18).

**Figure 5.**
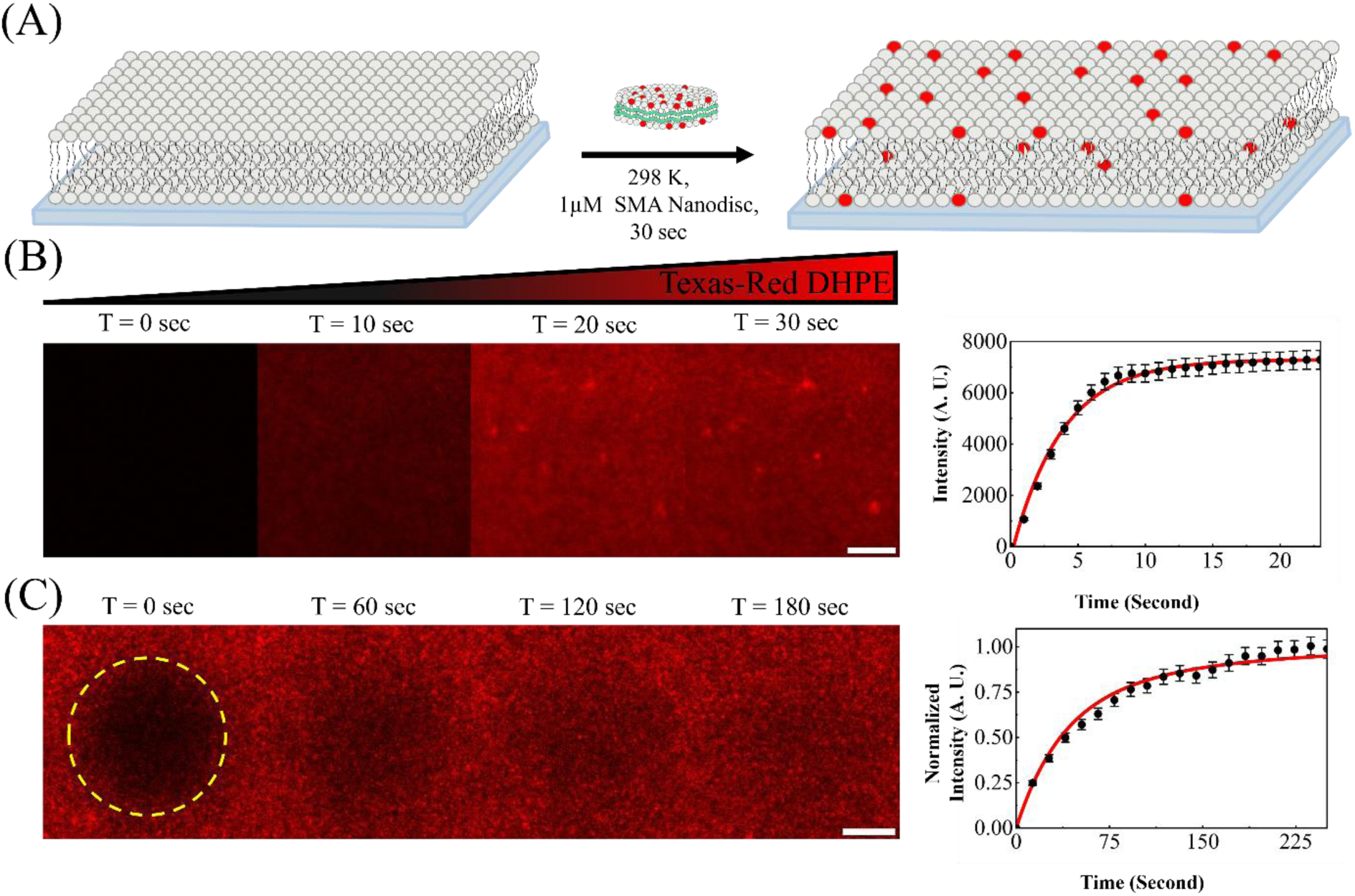
Rapid Membrane Integration and Subsequent Lateral Mobility of Lipids Delivered via SMA-Type Nanodiscs. (A) Schematic illustration demonstrating that SMA nanodiscs efficiently integrate a mixed lipid cargo into SLBs, analogous to the MSP platform, thereby doping the membranes with fluorescent lipids. (B) TIRF microscopy images of SLBs (left) capturing the rapid and homogeneous enhancement of fluorescence intensity immediately following the addition of 1 µM SMA nanodiscs. The corresponding time-course profile of fluorescence intensity (right) reveals that the delivery process reaches completion within one minute. (C) FRAP analysis validating the successful membrane integration of the lipid cargo, performed after removing residual unbound SMA nanodiscs via a TBS buffer wash. The robust fluorescence recovery within the photobleached area confirms that the delivered lipids possess lateral mobility and undergo efficient two-dimensional Brownian motion within the bilayer environment. The quantitative recovery trace of fluorescence intensity over time is presented on the right.

Furthermore, we validated the delivery capabilities of SMA nanodiscs in complex mammalian cell environments. As shown in the schematic in Figure 6A, SMA nanodiscs effectively transport lipid cargo to the plasma membranes of adherent cells, resulting in a distinct, membrane-localized fluorescent signal. Figure 6B demonstrates successful delivery across multiple mammalian cell lines. Notably, this nanodisc-mediated approach maintains high efficiency even in cell lines that are traditionally recalcitrant to chemical transient transfection, such as the OVCAR3 line. The fluorescence intensities over time during this delivery process are characterized in Figure 6C, showing that incorporation reaches completion within ten minutes. Finally, Figure 6D illustrates the concentration dependence of the delivery, where increased SMA nanodisc concentrations correlate with higher levels of lipid incorporation into the target cell membranes. Low-magnification imaging further showed that nearly all cells exhibited membrane-associated Texas Red-DHPE fluorescence after SMA nanodisc treatment, corresponding to an approximate 100% delivery efficiency under the tested conditions (Figure S8). Trypan Blue exclusion assays showed negligible intracellular staining and preserved cell adhesion, indicating near-complete cell viability following delivery.

**Figure 6.**
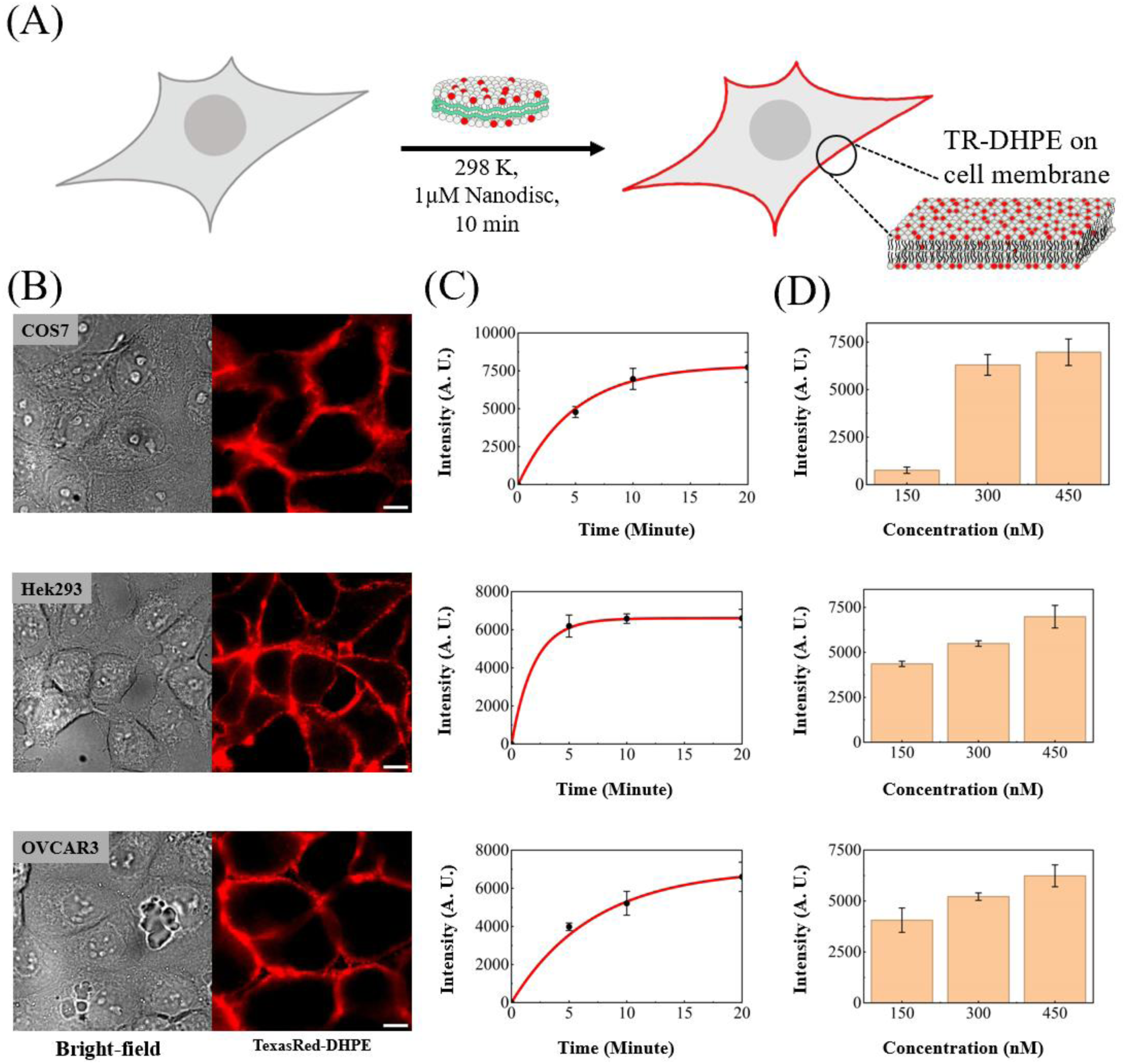
Efficient and Rapid SMA Nanodisc-Mediated Lipid Delivery into Diverse Mammalian Plasma Membranes. (A) Schematic illustration demonstrating the use of SMA nanodiscs to deliver lipid cargo into the plasma membranes of mammalian cells, resulting in robust membrane labeling at the cell periphery. (B) Bright-field (left) and corresponding epifluorescence images (right) of various mammalian cell lines treated with Texas Red-DHPE-loaded SMA nanodiscs. The images confirm that the fluorescent signal is predominantly localized to the cellular plasma membranes. (C) Time-course profile of fluorescence intensities at the cell periphery during the delivery process. Cells were incubated in imaging buffer containing 1 µM SMA nanodiscs, washed with imaging buffer after 10 min, and immediately imaged via epifluorescence microscopy. (D) Concentration dependence of lipid delivery assessed at SMA nanodisc concentrations of 150, 300, and 450 nM. Nanodiscs loaded with 1% Texas Red-DHPE and 99% DOPC were added to the cells in imaging buffer. Following an incubation period of 10 min, peripheral fluorescence intensities were quantified from epifluorescence images and plotted.

### SMA Nanodisc-Mediated Delivery and Functional Reconstitution of D2R

Building upon the successful validation of our polymer-based delivery platform, we report the preparation and purification of D2R-loaded SMA nanodiscs directly harvested from cellular membranes for the delivery and reconstitution of the receptor into naive target cells. To facilitate precise fluorescence tracking, the D2R construct used in this study contains an mNeonGreen fluorescent protein genetically fused to its C-terminus via a flexible linker (GSGSGSENLYFQGGSGSGS). This result demonstrates the transfer of full-length D2R between distinct membrane systems and highlights the utility of SMA nanodiscs for GPCR reconstitution. By using SMA nanodiscs as delivery vehicles, we achieved the functional reconstitution of D2R into a variety of naive cell lines lacking endogenous expression of the receptor.

As illustrated in the schematic in Figure 7A, D2R-loaded SMA nanodiscs were purified and subsequently applied to intact plasma membranes, enabling the direct integration of the receptor into naive target cells. The functional integrity of the reconstituted D2R was rigorously validated using a custom-synthesized fluorescent agonist, Alexa Fluor 647-labeled (S)-(−)-2-(N-phenethyl-N-propyl)amino-5-hydroxytetralin (PPHT). This probe was prepared by conjugating the Alexa Fluor 647 dye to a PPHT derivative via a linker using click chemistry. The bioactivity and specificity of the synthesized agonist were first confirmed in live-cell control experiments (Figure S19), where robust binding was observed exclusively in cells expressing D2R.

**Figure 7.**
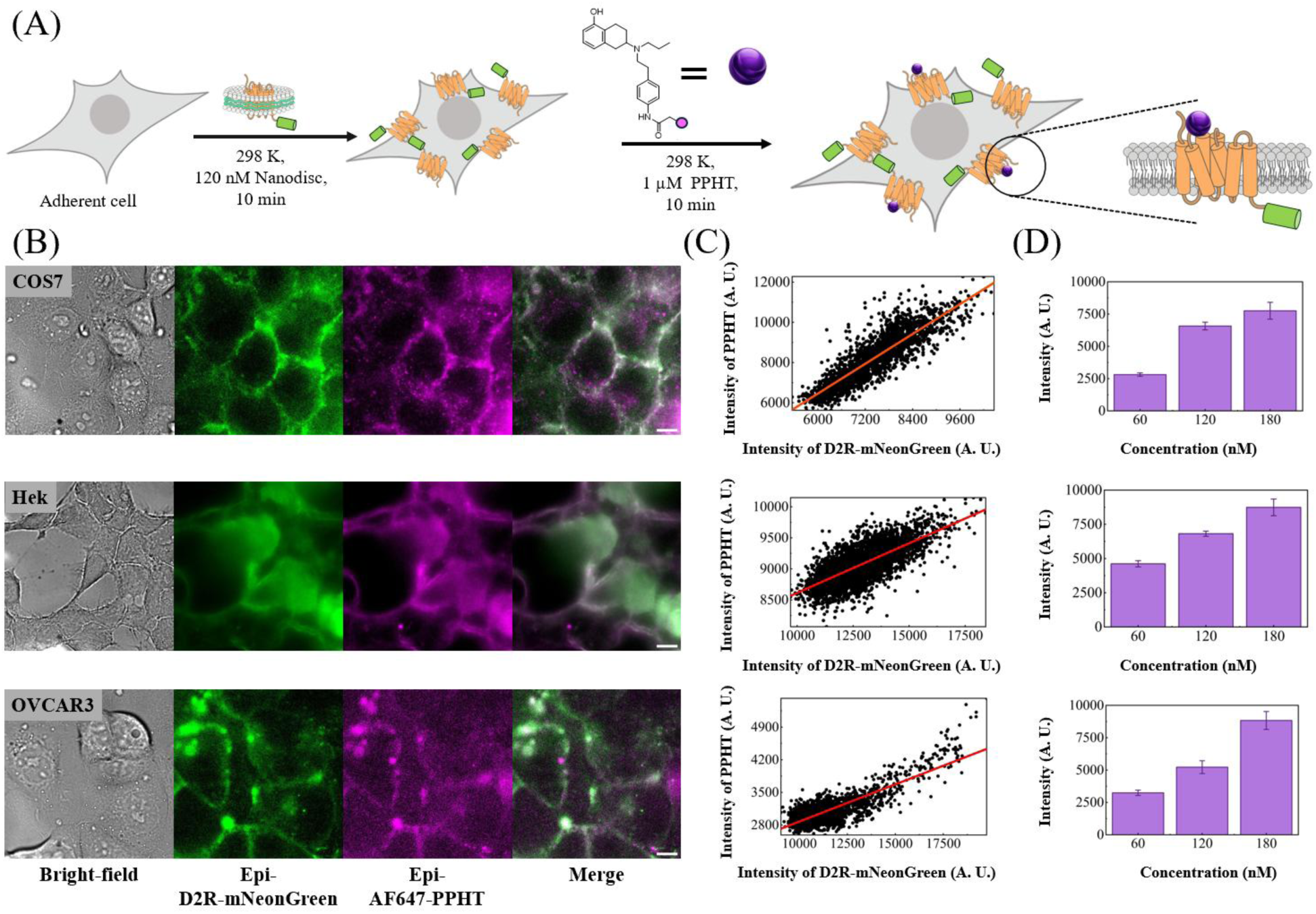
SMA Nanodisc-Mediated Delivery and Functional Reconstitution of D2R in Naive Target Cells. (A) Schematic illustration detailing the delivery of mNeonGreen-fused D2R to the cellular plasma membrane via SMA nanodiscs, followed by the introduction of a custom fluorescent agonist (Alexa Fluor 647-labeled PPHT) to evaluate receptor functionality. Unbound agonist was removed via an imaging buffer wash prior to fluorescence visualization. (B) Bright-field and corresponding epifluorescence images of various mammalian cell lines. Merged composite images reveal robust spatial colocalization between the D2R (mNeonGreen) and agonist (Alexa Fluor 647) signals, visually confirming specific receptor-ligand binding post-delivery. (C) 2D cytofluorograms illustrating the spatial fluorescence intensity correlation between the mNeonGreen and Alexa Fluor 647 channels across the imaging field (using 4× 4 pixel binning). Pearson colocalization analysis yields strong positive correlation coefficients (*r*) ranging from 0.72 to 0.87 across all tested cell lines, mathematically corroborating the highly specific binding interaction. (D) Concentration-dependent binding analysis of the fluorescent agonist to the reconstituted D2R. Cells were uniformly treated with a standardized concentration of D2R-loaded SMA nanodiscs, followed by a 10-minute incubation with varying concentrations of the ligand. Quantification of the resulting epifluorescence intensities confirms that increasing the ligand concentration drives a corresponding enhancement in receptor binding occupancy.

Figure 7B presents epifluorescence images of D2R reconstitution across multiple cell lines. The successful integration of the receptor was confirmed by the strong membrane-localized fluorescence of an mNeonGreen tag fused to the C-terminus of D2R. Upon the addition of our synthesized agonist, we observed high spatial colocalization between the agonist and the reconstituted D2R, indicating that the receptor maintains its native ligand-binding conformation and functionality post-delivery. Pearson colocalization analysis and spatial fluorescence intensity profiling (Figure 7C) further corroborate the highly specific binding between D2R and the ligand. Furthermore, Figure 7D illustrates the concentration dependence of this interaction; increasing the agonist concentration yielded a corresponding enhancement in ligand fluorescence, reflecting higher binding occupancy at the reconstituted D2R sites. Critcally, control experiments in which the fluorescent ligand was introduced to untreated naive cells (lacking D2R-loaded SMA nanodiscs) exhibited only basal background fluorescence (Figure S20). This indicates that non-specific binding is negligible and suggests that endogenous D2R expression in these target membranes is below the detection limit of our assay. These control results provide strong evidence that the robust PPHT fluorescence observed in Figure 7B is primarily attributable to specific binding events between the delivered receptor and the ligand.

Future studies could extend this fluorescence-based approach to quantitatively characterize the delivered receptor population. With appropriate fluorescence calibration, the mNeonGreen-tagged D2R and fluorescent PPHT signals could be used to estimate the numbers of delivered receptors and bound ligands, respectively (55). Such measurements could help determine the fraction of delivered D2R that remains functionally competent after membrane transfer. Future work could also examine whether all membrane-associated receptors adopt an appropriate transmembrane orientation and insertion state for ligand binding, as well as whether a subset of the delivered receptors subsequently undergoes membrane trafficking and redistributes away from the plasma membrane. Quantitative analysis of receptor abundance, ligand occupancy, and membrane localization could therefore provide a more complete understanding of the functional state and cellular behavior of nanodisc-delivered D2R.

Crucially, this delivery method enables the rapid, on-demand introduction of D2R into living cell membranes and provides a direct means of monitoring the immediate cellular consequences of membrane protein incorporation. Unlike MSP nanodiscs, which require the in vitro assembly of purified scaffold proteins, lipids, and detergent-stabilized membrane proteins, SMA nanodiscs can directly capture membrane proteins from the plasma membranes of expressing cells and subsequently transfer them into distinct target cell membranes. This cross-membrane transfer strategy circumvents the time and experimental resources required for genetic knock-in approaches and avoids the delay associated with de novo protein expression following transient transfection. As a result, cellular responses can be monitored immediately after receptor introduction, providing a complementary approach for dissecting early membrane-protein-dependent events.

Such capabilities are particularly relevant to GPCRs, which coordinate complex interactions between small-molecule ligands and larger downstream signaling proteins at the membrane interface. The ability to introduce defined receptor populations into user-selected membrane environments therefore offers a bottom-up strategy for examining receptor function and signaling with temporal control that complements conventional live-cell expression approaches. By separating receptor delivery from endogenous biosynthesis, this platform may also facilitate the observation of transient or early-stage molecular events that are difficult to resolve during prolonged expression. To further demonstrate the versatility of this approach across distinct membrane protein classes, we showed that SMA nanodiscs can also deliver functional BsYetJ ion channels, which mediate real-time intracellular calcium influx in living cells (Figure S21). Together, these findings establish SMA nanodisc-mediated membrane-to-membrane transfer as a broadly applicable platform for investigating GPCRs, ion channels, and other complex membrane proteins in diverse, user-defined cellular environments.

## Conclusion

In summary, we have established the nanodisc as a robust and broadly applicable carrier for the targeted delivery of lipids and membrane proteins across disparate membrane systems. By utilizing this versatile platform, we successfully achieved the functional reconstitution of two complex, multi-pass transmembrane proteins: the bacterial calcium channel BsYetJ and the G protein-coupled receptor D2R. Importantly, this platform was successfully applied to intact mammalian plasma membranes across multiple cell lines. Nanodisc-delivered BsYetJ generated robust calcium influx signals, confirming that the reconstituted channel retained its functional conformation after transfer into a heterologous cellular membrane. By directly capturing full-length D2R from cellular membranes and transferring it into naive target cells, we demonstrated functional GPCR reconstitution, as validated by specific binding of a custom fluorescent agonist. Collectively, these results demonstrate that nanodisc-mediated delivery provides a rapid, modular, and broadly applicable strategy for introducing complex membrane proteins into user-defined membrane environments. This targeted delivery platform provides a critical technical foundation for the real-time, *in vitro*, and *in vivo* interrogation of challenging pharmacological targets, such as GPCRs and ion channels, in their native-like environments.

## Materials and Experimental Methods

### Materials

1,2-Dioleoyl-*sn*-glycero-3-phosphocholine (DOPC), 1,2-dioleoyl-*sn*-glycero-3-phosphate (sodium salt) (DOPA), and 1,2-dioleoyl-*sn*-glycero-3-[(N-(5-amino-1-carboxypentyl) iminodiacetic acid) succinyl] (Ni^2+^-NTA-DOGS, nickel salt) were purchased from Avanti Polar Lipids. Texas Red 1,2-dihexadecanoyl-*sn*-glycero-3-phosphoethanolamine (TR-DHPE, triethylammonium salt) and Alexa Fluor 647 C2 maleimide were obtained from Thermo Fisher Scientific. Sulfuric acid (H2SO4) and hydrogen peroxide (H2O2) were sourced from Honeywell Fluka, and Tris-buffered saline (TBS) was purchased from Protech Technology. Fetal bovine serum (FBS), penicillin-streptomycin solution (100×), Dulbecco’s modified Eagle medium (DMEM), Roswell Park Memorial Institute medium (RPMI), and sodium pyruvate were supplied by Corning Life Sciences. GlutaMAX and insulin were purchased from Thermo Fisher Scientific. Unless otherwise noted, all reagents were used as received.

### Preparation of Supported Lipid Bilayers

Supported lipid bilayers (SLBs) were prepared via vesicle rupture on cleaned glass substrates. Small unilamellar vesicles (SUVs) were formulated from a 98:2 molar ratio of DOPC and DOPA in chloroform. The lipid mixture was concentrated using a rotary evaporator at 40 °C for 3 min, followed by drying under a stream of nitrogen for 15 min to ensure complete solvent removal. The dried lipid film was resuspended in deionized water to yield a final lipid concentration of 2 mg mL⁻¹. The suspension was then vortexed and pipetted thoroughly, and SUVs were generated via probe-tip sonication. Glass coverslips (bottom thickness 170 ± 5 µm; Ibidi) were cleaned with piranha solution (H₂SO₄/H₂O₂, 3:1 v/v) for 8 min and assembled with flow chambers (Sticky-Slide VI 0.4; Ibidi). The SUV suspension was mixed 1:1 (v/v) with TBS buffer, injected into the chamber, and incubated for 20 min to form SLBs.

### Preparation of MSP Nanodiscs

Membrane scaffold protein MSP1D1, herein referred to as MSP, was expressed in *E. coli* and purified as previously described (40). The MSP-encoding pET-28a plasmid was transformed into *E. coli* BL21(DE3) cells, and transformants were selected on kanamycin-containing agar plates (30 µg mL⁻¹) at 37 °C. A single colony was used to inoculate 10 mL of Terrific Broth (TB) supplemented with kanamycin (30 µg mL⁻¹). After the culture reached an OD₆₀₀ of 0.6–0.8, it was expanded into 0.6 L of TB containing kanamycin. Protein expression was induced with 1 mM IPTG when the OD₆₀₀ reached approximately 2.0–2.5, and the culture was incubated for an additional 4 h at 28 °C. Cells were collected by centrifugation and stored at −80 °C until purification.

For MSP purification, frozen cell pellets were resuspended in 30 mL of lysis buffer containing 20 mM sodium phosphate, 0.1 M NaCl, 1% Triton X-100, and 10 mM MgSO₄ at pH 7.4. DNase I (10 µg mL⁻¹) and phenylmethylsulfonyl fluoride (PMSF; 300 µL of a 0.1 M solution in ethanol) were added immediately before lysis. Cells were disrupted by sonication, and the lysate was clarified by centrifugation at 12800 × g for 50 min. The supernatant was applied to a HisTrap HP column (5 mL; GE Healthcare) pre-equilibrated with 20 mM sodium phosphate, 0.1 M NaCl, and 1% Triton X-100 at pH 7.4. The column was washed sequentially with 25 mL of 40 mM Tris-HCl, 0.3 M NaCl, and 1% Triton X-100 at pH 8.0, followed by 25 mL of 40 mM Tris-HCl, 0.3 M NaCl, and 50 mM sodium cholate at pH 8.0, and then 25 mL of 40 mM Tris-HCl, 0.3 M NaCl, and 40 mM imidazole at pH 8.0. MSP was eluted with 40 mM Tris-HCl, 0.3 M NaCl, and 0.4 M imidazole at pH 8.0. The eluted protein was buffer-exchanged into MSP buffer consisting of 20 mM Tris-HCl and 0.1 M NaCl at pH 7.4, concentrated to approximately 10 mg mL⁻¹ using a 10 kDa MWCO centrifugal concentrator, and quantified by absorbance at 280 nm using an extinction coefficient of 21430 M⁻¹ cm⁻¹.

### Preparation of nanodisc samples

MSP nanodiscs were assembled using defined lipid/MSP and sodium cholate/lipid molar ratios of 65:1 and 2:1, respectively. For fluorescent nanodisc preparation, DOPC was mixed with 4 mol% fluorophore-conjugated lipid, either FITC-DHPE or Texas Red-DHPE, in chloroform. At this labeling density, more than 99.5% of nanodiscs are expected to contain at least one fluorescent lipid, based on a binomial distribution. The lipid mixture was dried under a gentle stream of nitrogen for 30 min to form a thin lipid film. The dried lipids were solubilized with sodium cholate and sonicated until the solution became clear. Purified MSP was then added to the detergent-solubilized lipid mixture, and the sample was incubated on ice for 30 min.

Nanodisc self-assembly was initiated by detergent removal using SM-2 Bio-Beads (1 g mL⁻¹; Bio-Rad). The mixture was incubated with Bio-Beads at 4 °C for 5 h, after which the beads were removed by centrifugation. The assembled nanodiscs were further purified by size-exclusion chromatography using a Superdex 200 10/300 GL column. Fractions containing monodisperse nanodiscs were collected, buffer-exchanged into nanodisc buffer containing 50 mM HEPES and 0.1 M NaCl at pH 7.0, and concentrated using Amicon Ultra-50K centrifugal filter units. The final nanodisc concentration was determined from the absorbance at 280 nm.

### Preparation of D2R-Loaded SMA Nanodiscs

The cells were harvested and resuspended in a hypotonic buffer containing protease inhibitors. Following incubation on ice, the cells were disrupted using a Dounce homogenizer. The lysate was initially cleared of cellular debris by low-speed centrifugation (1,000×g, 15 min), and the resulting supernatant was subjected to ultracentrifugation (45,000×g, 1 h) to pellet the crude membrane fraction. The obtained membrane pellet was resuspended in SMA buffer (pH 8.0, 5 mM imidazole) and homogenized. To form styrene-maleic acid lipid particles (SMALPs), the membrane solution was mixed with an equal volume of Styrene-Maleic Acid (SMA) copolymer buffer to achieve a final concentration of 1.5% SMA. The mixture was incubated at room temperature for 2 hours with continuous rotation to facilitate D2R extraction. Non-solubilized material was removed by another round of ultracentrifugation (45,000×g, 1 h). The supernatant containing the solubilized D2R SMALPs was incubated overnight with pre-equilibrated Ni-NTA resin. The mixture was loaded onto a gravity flow column, and the non-binding flow-through was collected. The resin was washed sequentially with SMA buffers containing increasing concentrations of imidazole (20 mM and 40 mM) to remove non-specific proteins. Target D2R SMALPs were subsequently eluted with a high-imidazole buffer (300 mM). The collected fractions were analyzed by SDS-PAGE and further purified using size exclusion chromatography.

### Preparation of Empty SMA Nanodiscs

A chloroform solution containing 10 mg of total lipid (DOPC/TR-DHPE, 99:1 molar ratio) was transferred to a round-bottom glass tube. Glass tubes and pipettes were used exclusively to prevent polymer leaching from plastic materials in the presence of chloroform. The solution was dried under vacuum by rotary evaporation for at least 30 min, followed by drying under a gentle stream of nitrogen to yield a homogeneous thin lipid film. After additional vacuum drying to remove residual solvent, the film was rehydrated with 50 mM Tris buffer (1 mL) by vigorous vortexing, with warming to 50 °C applied when necessary to ensure complete rehydration. The mixture was then bath-sonicated to produce a translucent, milky-white SUV suspension. An SMA solution (1 mL, 5% w/v) was added dropwise to the SUV suspension. Formation of empty SMA nanodiscs (SMANDs) was indicated by clarification of the solution upon SMA-mediated vesicle solubilization and further confirmed by dynamic light scattering (DLS) analysis of the size distribution. The empty SMA nanodiscs were further purified by liquid chromatography using a size-exclusion column. Purified SMANDs were stored at 4 °C and used within two weeks.

### Nanodisc Loading onto Supported Lipid Bilayers and Adherent Cells

For SLB loading, nanodisc solution (100 µL, 1 µM in TBS buffer) was introduced into the microfluidic flow chamber containing the preformed SLB at 25 °C. Delivery was monitored via time-lapse TIRF microscopy and quantified by assessing the temporal change in mean fluorescence intensity or single-molecule fluorescence spot density within the imaging field. For adherent cells, nanodisc solution (200 µL, 1 µM in Tris-based imaging buffer) was added to cells in an ibiTreat No. 1.5 polymer-bottom µ-Slide 8-well plate (Ibidi). Following a 10 min incubation at 25 °C to allow for membrane delivery, wells were washed two to three times with imaging buffer prior to microscopy.

### Cell Culture

HEK293 and COS7 cells (Bioresource Collection and Research Center, #60019 and #60094) were seeded in ibiTreat 8-well plates at a density of 3.6 × 10^5^ cells per well. They were cultured in 198 µL DMEM supplemented with 10% FBS, 1% penicillin/streptomycin, 1 mM sodium pyruvate, and 2 mM GlutaMAX at 37 °C under 5% CO2. NIH:OVCAR3 cells (#60551) were seeded similarly and maintained in 198 µL RPMI medium containing 20% FBS, 1% penicillin/streptomycin, 1 mM sodium pyruvate, 0.01 mg mL^-1^ bovine insulin, and 2 mM GlutaMAX at 37 °C under 5% CO2.

### Fluo-8 AM Loading and Calcium Influx Assay

Following 24 h of culture, cells were washed with sterile Tris buffer (25 mM Tris, 143 mM NaCl, 3 mM KCl, 1 mM MgCl₂, 5.5 mM D-glucose, pH 7.4) and incubated with Fluo-8 AM (5 µM; AAT Bioquest) and EGTA (0.2 mM) at 37 °C for 15 min. After excess dye was removed by two washes with Tris buffer, a transmembrane calcium gradient was established immediately before imaging by replacing two-thirds of the buffer with calcium-containing imaging buffer, yielding a final extracellular Ca²⁺ concentration of 2 mM. The delivery of the membrane protein was initiated by adding Alexa Fluor 647-labeled BsYetJ nanodiscs or empty nanodisc controls.

### Microscopy and Image Analysis

Microscopy and Image Analysis: Total Internal Reflection Fluorescence (TIRF) imaging was conducted using a Nikon Eclipse Ti inverted microscope equipped with a TIRF system and an iXon EMCCD camera (Andor Technology). Imaging was performed with a 100× 1.49 NA oil-immersion objective, a Perfect Focus system, and a U-N4S four-laser unit. Solid-state lasers at 488, 561, and 640 nm were used with measured powers of 5.2, 6.9, and 7.8 mW, respectively. Emission was collected through a 405/488/561/638 nm Quad TIRF filter, and images were acquired using Nikon NIS-Elements software. Wide-field epifluorescence imaging was performed using standard LED illumination, either Nikon Intensilight C-HGFIE or Lumencor Spectra X LED, with a 100× 1.40 NA oil-immersion objective and appropriate filter sets for DAPI, FITC/GFP, Texas Red, and Cy5. Typical exposure times were 50–200 ms. Fluorescence intensity changes were quantified using built-in ImageJ functions. Single-molecule movies were analyzed using the ImageJ plugin TrackMate(62) to identify molecular spots and obtain single-particle trajectories.

### D2R Expression in Expi293F Cells

Expi293F cells (Thermo Fisher Scientific) were maintained in Expi293 Expression Medium at 37 °C under 8% CO₂ with continuous agitation at 125 rpm. Cells were subcultured when the density reached 3 × 10⁶ to 5 × 10⁶ cells mL⁻¹ to maintain logarithmic growth. Transient transfection was performed using the ExpiFectamine 293 Transfection Kit. Cells were diluted to 3 × 10⁶ cells mL⁻¹ in a total volume of 25 mL and transfected with target membrane protein plasmid DNA (25 µg) and ExpiFectamine 293 reagent, both pre-incubated in Opti-MEM I Reduced Serum Medium. ExpiFectamine 293 Transfection Enhancers 1 and 2 were added 18–22 h post-transfection. Cells were harvested 72 h post-transfection by centrifugation at 3000g for 10 min.

### Chemical Synthesis of the Alexa Fluor 647-Labeled PPHT Ligand

The detailed synthetic route for the precursor PPHT ligand is provided in the Supporting Information. The subsequent steps detailing the conjugation of AF647 to PPHT are outlined in Scheme 1 and described below.

**Scheme 1.**
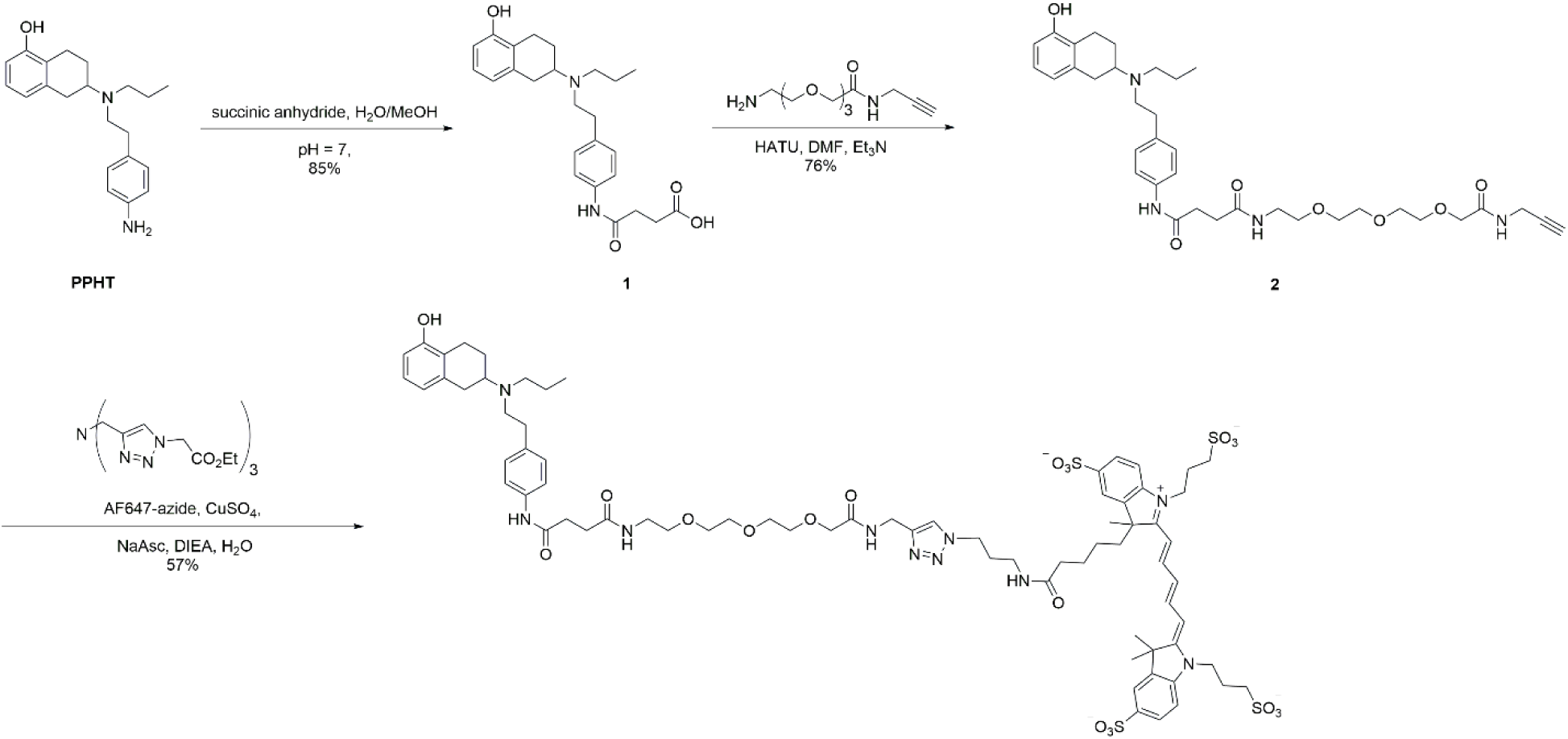
Synthesis of the AF647-labeled PPHT fluorescent agonist.

### 4-((4-(2-((5-hydroxy-1,2,3,4-tetrahydronaphthalen-2-yl)(propyl)amino)ethyl)phenyl)amino)-4-oxobutanoic acid (1)

To a solution of **PPHT** (20.0 mg, 0.062 mmol) in 1.24 mL toluene was added succinic anhydride (7.0 mg, 0.068 mmol) and stirred for 1 h at 50 °C. The reaction mixture was then cooled to room temperature and concentrated by rotary evaporation under reduced pressure to give a white solid. The solid was then added 0.43 mL THF, 0.14 mL H2O and lithium hydroxide monohydrate (0.003 g, 0.081 mmol), the reaction mixture was then stirred for overnight. The crude mixture was purified by HPLC and the desired product was obtained as a white solid **1** (15.0 mg, 65%). ^1^H NMR (400MHz, CD3OD) δ 7.51 (d, J = 7.8 Hz, 2H,ArH), 7.25 (d, J = 7.7 Hz, 2H, ArH), 6.95 (t, J = 7.6 Hz, 1H, ArH), 6.62 (d, J = 8.7 Hz, 1H, ArH), 6.60 (d, J = 9.8 Hz, 1H, ArH), 3.78 (m, 1H, ArC*H2*CHN), 3.54 (m, 1H, ArC*H2*CH2CHN), 3.40 (m, 1H, ArC*H2*CH2CHN), 3.24 (m, 1H, ArC*H2*CHN), 3.14-3.05 (m, 4H, ArC*H2*C*H2*N) , 3.14-3.05 (m, 1H, ArCH2C*H*N), 2.70-2.60 (m, 4H, NHCOC*H2*C*H2*COOH), 2.70-2.60 (m, 1H, NC*H2*CH2CH3), 2.33 (m, 1H,NC*H2*CH2CH3), 1.94-1.82 (m, 1H, ArCH2C*H2*CHN), 1.94-1.82 (m, 2H, NCH2C*H2*CH3), 1.35-1.29 (m, 1H, ArCH2C*H2*CHN), 1.06 (t, J = 7.2 Hz, 3H, NCH2CH2C*H3*). ^13^C NMR (100.6 MHz, CD3OD) δ 176.30, 172.92, 156.17, 139.28, 134.72, 132.92, 130.25, 128.08, 123.08, 121.70, 121.25, 113.54, 62.08, 53.94, 53.42, 32.32, 31.76, 30.82, 29.96, 24.; ESI-MS (m/z): [M+H] ^+^ calcd for C25H33N2O4: 425.2440, found: 425.2446.

### N1-(4-(2-((5-hydroxy-1,2,3,4-tetrahydronaphthalen-2-yl)(propyl)amino)ethyl)phenyl)-N4-(11-oxo-3,6,9-trioxa-12-azapentadec-14-yn-1-yl)succinamide (2)

To a solution of compound **1** (0.50 mg, 1.18 μmol) in Dimethylformamide (60 μL) added HATU (1.79 mg, 4.72 μmol), 2-(2-(2-(2-aminoethoxy)ethoxy)ethoxy)-N-(prop-2-yn-1-yl)acetamide (0.86 mg, 3.54 μmol) and triethylamine (0.82 μL), then stirred at room temperature for 2 h. The crude product was purified by HPLC with a reversed-phase C18 column, and was lyophilized to give white solid **2** (0.58 mg, 0.90 μmol, 76 %). ^1^H NMR (850 MHz, CD3OD) : δ 7.54 (d, *J* = 8.5 Hz, 2H, ArH), 7.28 (d, *J* = 8.5 Hz, 2H, ArH), 6.97 (t, J = 7.7 Hz, 1H, ArH), 6.65 (d, *J* = 7.6 Hz, 1H, ArH), 6.62 (d, *J* = 7.9 Hz, 1H, ArH), 4.02-3.99 (m, 4H, NCH2CH2OCH2CH2OCH2CH2OC*H2*CONH, NHC*H2*CCH), 3.78-3.72 (m, 1H, ArCH2C*H*N),3.70-3.63 (m, 8H, NCH2CH2OC*H2*C*H2*OC*H2*C*H2*OCH2CONH), 3.55 (t, *J* = 5.5 Hz, 2H, NCH2C*H2*OCH2CH2OCH2CH2OCH2CONH), 3.50-3.40 (m, 1H, ArC*H2*CH2CHN), 3.38 (t, *J* = 5.5 Hz, 2H, NC*H2*CH2OCH2CH2OCH2CH2OCH2CONH), 3.30-3.22 (m, 1H, ArC*H2*CHN), 3.17-3.02 (m, 5H, ArC*H2*C*H2*N, ArCH2C*H*N), 2.69-2.61 (m, 3H, ArCH2C*H*N, NC*H2*CH2Ar), 2.62-2.52 (m, 3H, NCH2C*H2*Ar, NHCH2CC*H*), 2.36-2.31 (m, 1H, ArCH2C*H2*CHN), 1.94-1.88 (m, 1H, ArCH2C*H2*CHN), 1.88-1.80 (m, 2H, NCH2C*H2*CH3), 1.31-1.26 (m, 1H, ArCH2C*H2*CHN), 1.06 (t, *J* = 7.3 Hz, 3H, NCH2CH2C*H3*). ^13^C NMR (212.5 MHz, CD3OD) δ 174.77, 173.00, 172.55, 156.17, 139.25, 134.88, 133.08, 130.26, 128.06, 123.13, 121.64, 121.27, 113.49, 80.65, 72.21, 71.98, 71.54, 71.46, 71.28, 71.27, 70.60, 61.90, 53.93, 53.42, 40.47, 32.93, 31.92, 31.81, 30.91, 28.98, 25.01, 23.73, 20.00, 11.35.; ESI-MS (m/z): [M+H] ^+^ calcd for C36H50N4O7: 650.3740, found: 650.3751.

### AF647-labeled PPHT

Tris(triazoly)-amine ligand (4.2 equiv., 40 mM in DMSO), CuSO4 (aq) (4.2 equiv., 40 mM in H2O), and sodium ascorbate (aq) (6.3 equiv., 800 mM) was added to a vial. To a solution of the compound **2** (0.62 mg, 0.96 μmol) and AF647-azide(1.0 mg, 0.96 μmol) in water (1 mM) was added the aforementioned solution and DIEA (4.0 equiv.) to react at room temperature for 3 h. The product was purified by HPLC with a Vydac C8 column, and was lyophilized to give white solid **PPHT-AF647** (0.86 mg, 0.55 μmol, 57 %). ^1^H NMR (850 MHz, d^6^-DMSO) : δ 9.94 (br s, 1H, ArN*H*CO), 9.38 (br s, 1H, ArO*H*), 9.19 (br s, 1H, C-23N*H*C-24), 8.38-8.28 (m, 2H, H-52, H-52’), 7.95-7.91 (m, 2H, H-49, H-49’), 7.80-7.75 (m, 2H, H-47, H-47’), 7.73 (s, 1H, H-34), 7.63-7.59(m, 2H, H-46, H-46’), 7.54-7.49 (m, 2H, H-18), 7.39 (d, 1H, *J* = 7.91 Hz, H-55), 7.34 (d, 1H, *J* = 8.23 Hz, H-51), 7.21 (d, 2H, *J* = 8.08 Hz, H-17), 6.95 (td, 1H, *J* = 7.65, 2.81 Hz, H-7), 6.63 (d, 1H, *J* = 7.91 Hz, H-8), 6.58 (d, 1H, *J* = 7.65, H-6), 6.48 (t, 1H, *J* = 12.24, H-53), 6.41 (br s, 2H, CON*H*), 4.34 (d, 2H, *J* = 5.87, H-30), 4.31-4.22 (m, 4H, H-56, H-56’), 4.22-4.17 (m, 2H, H-35), 3.91 (s, 2H, H-32), 3.74-3.67 (m, 1H, H-10), 3.60-3.51 (m, 4H, H-26, H-27), 3.49-3.44 (m, 4H, H-28, H-29), 3.15-2.94 (m, 6H, H-24, H-25, H-37), 2.94-2.87 (m, 2H, H-11), 2.87-2.80 (m, 1H, H-10), 2.66-2.59 (m, 3H, H-1, H-3), 2.59-2.52 (m, 8H, H-14, H-15, H-21, H-22), 2.30-2.24 (m, 1H, H-39), 2.16-2.09 (m, 1H, H-39), 2.02-1.96 (m, 4H, H-57, H-57’), 1.89-1.84 (m, 2H, H-12), 1.84-1.70 (m, 5H, H-2, H-36, H-42), 1.70-1.60 (m, 9H, C-43C*H3*, C-43’C*H3*, C-43’C*H3*), 1.34-1.28 (q, 1H, *J* = 7.48 Hz, H-40), 1.28-1.21 (m, 1H, H-2), 0.94 (t, 3H, *J* = 6.8 Hz, H-13), 0.77-0.68 (m, 1H, H-41), 0.45-0.38 (m, 1H, H-41). ^13^C NMR (212.5 MHz, d^6^-DMSO) δ 171.93, 171.32, 170.44, 169.29, 154.77, 154.45, 153.60, 145.22, 144.97, 144.59, 143.10, 142.06, 138.17, 134.26, 131.12, 131.08, 129.04, 126.50, 126.10, 126.06, 125.98, 122.96, 121.87, 119.84, 119.64, 119.60, 119.08, 112.14, 110.19, 109.87, 70.28, 69.88, 69.66, 69.54, 59.52, 59.06, 59.52, 53.20, 51.70, 51.68, 51.59, 51.49, 48.95, 48.08, 47.82, 47.07, 38.58, 35.59, 35.20, 33.85, 31.67, 30.29, 29.88, 29.85, 29.78, 29.33, 28.98, 27.02, 26.96, 25.54, 23.65, 23.46, 23.34, 23.06, 22.70, 22.44, 18.33, 18.23, 11.01, 10.96.; ESI-MS (m/z): [M-3H] ^3-^ calcd for C74H97N10O20S43-: 524.5260, found: 524.5260.

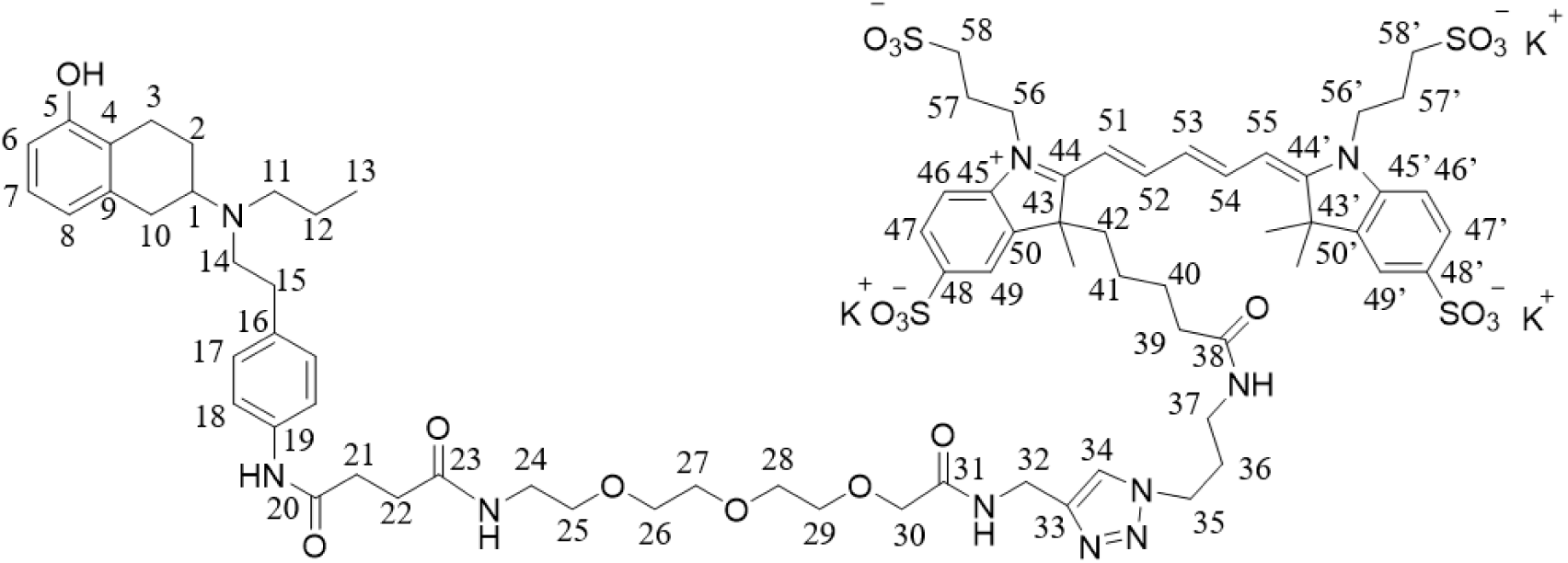

### BsYetJ Expression and Purification

BsYetJ was expressed and purified as described previously (41). A pET-24-derived pNYCOMPS vector encoding C-terminal His-tagged BsYetJ was used. Site-directed mutations were introduced using the QuikChange mutagenesis kit (Invitrogen), confirmed by DNA sequencing, and transformed into *E. coli* BL21(DE3) pLysS cells (Agilent). For expression, 10 mL overnight culture was used to inoculate 0.5 L TB medium containing 50 μg/mL kanamycin and 25 μg/mL chloramphenicol. Cultures were grown at 37 °C, induced at OD600 = 0.6–0.8 with 0.4 mM IPTG, and harvested after 4 h by centrifugation at 7000 × g for 5 min at 4 °C. Pellets were stored at –80 °C.

For purification, pellets were resuspended in lysis buffer containing 50 mM HEPES, 0.3 M NaCl, 20 mM imidazole, 5% glycerol, and 1 mM MgCl2, pH 7.8, and disrupted by sonication. Lysates were clarified by centrifugation and ultracentrifugation to isolate membranes. Membranes were solubilized with 1.5% β-DDM for 2 h, filtered, and loaded onto a 5 mL HisTrap HP column. After washing, BsYetJ was eluted with imidazole-containing buffer, exchanged into storage buffer, and treated overnight with TEV protease at 4 °C. Untagged BsYetJ was isolated by reverse HisTrap purification, and purity was assessed by SDS–PAGE.

### GUV electroformation

Giant unilamellar vesicles (GUVs) were prepared by electroformation using indium tin oxide (ITO)-coated glass slides (3 × 3 cm). Lipid mixtures containing DOPC, 2 mol% DOPA, and 0.2 mol% 18:1 Biotinyl Cap PE were dissolved in chloroform at a total lipid concentration of 1.0 mg/mL. The lipid solution was deposited onto the conductive surface of each ITO-coated slide and evenly spread to form a thin lipid film. The films were dried under a gentle stream of nitrogen (N₂) for 30 min to remove residual solvent.

Two lipid-coated ITO slides were assembled with their conductive surfaces facing inward and separated by a 2-mm-thick rubber spacer containing a 15-mm-diameter hole. The chamber was filled with 400 μL of 250 mM sucrose solution and connected to a custom-made AC field generator. Electroformation was performed using a sinusoidal AC field consisting of sequential swelling (1.0 Vpp, 15 Hz, 60 min), growth (3.0 Vpp, 10 Hz, 120 min), and detachment (3.0 Vpp, 2 Hz, 30 min) phases. The resulting GUV suspension was collected immediately and used for subsequent experiments.

## Author Contributions

L.-K.C., C.-K.L., S.-K.W., Y.-W.C. and C.-W.L. designed research; L.-K.C., C.-K.L. and T.-T. H. performed research; P.-T.H., M.-C.Y. and W.-X.L. contributed critical reagents and important experimental materials; L.-K.C., C.-K.L., T.-T.H., S.-K.W., Y.-W.C. and C.-W.L. analyzed and discussed data; and S.-K.W., Y.-W.C. and C.-W.L. wrote the paper.

## Acknowledgements

This work was supported by grants from the National Science and Technology Council of Taiwan (114-2113-M-007-015 and 114-2113-M-007-023) and the Ministry of Education of Taiwan (MOE-111-YSFMS-0002-003-P1). We gratefully acknowledge Prof. Chun-Cheng Lin and Prof. Shey-Cherng Tzou for their assistance in establishing Expi293F cell culture and the corresponding membrane protein expression workflow in our laboratory. We thank Prof. Ching-Ching Yu for providing access to instruments for DNA cloning and protein purification. We are also grateful to Jing Zheng Enterprise Ltd. for assistance in developing the customized GUV electroformation device used for GUV experiments, as well as the specialized flow chambers and chambered coverslips used in this study.

