## Supporting information for "An On-Demand Nanodisc Platform for Reconstitution of Functional Membrane Proteins into Model and Living Membranes"

**Title:**

### Delivering Active GPCRs and Ion Channels: Nanodisc-Mediated Reconstitution of D2R and BsYetJ Across Model and Living Membrane Systems

Long-Kai Chen, Yun-Shan Wang, Wei-Hsuan Chang, Che-Kai Lin, Pei-Tzu Huang, Ming-Chang Yu, Wen-Xin Liu, Tzu-Ting Huang, Chien-Yu Ko, Ru-Hsuan Bai, Sheng-Kai Wang*, Yun-Wei Chiang* and Chun-Wei Lin*

Department of Chemistry, National Tsing Hua University, Hsinchu, Taiwan 300044

### Supporting Information

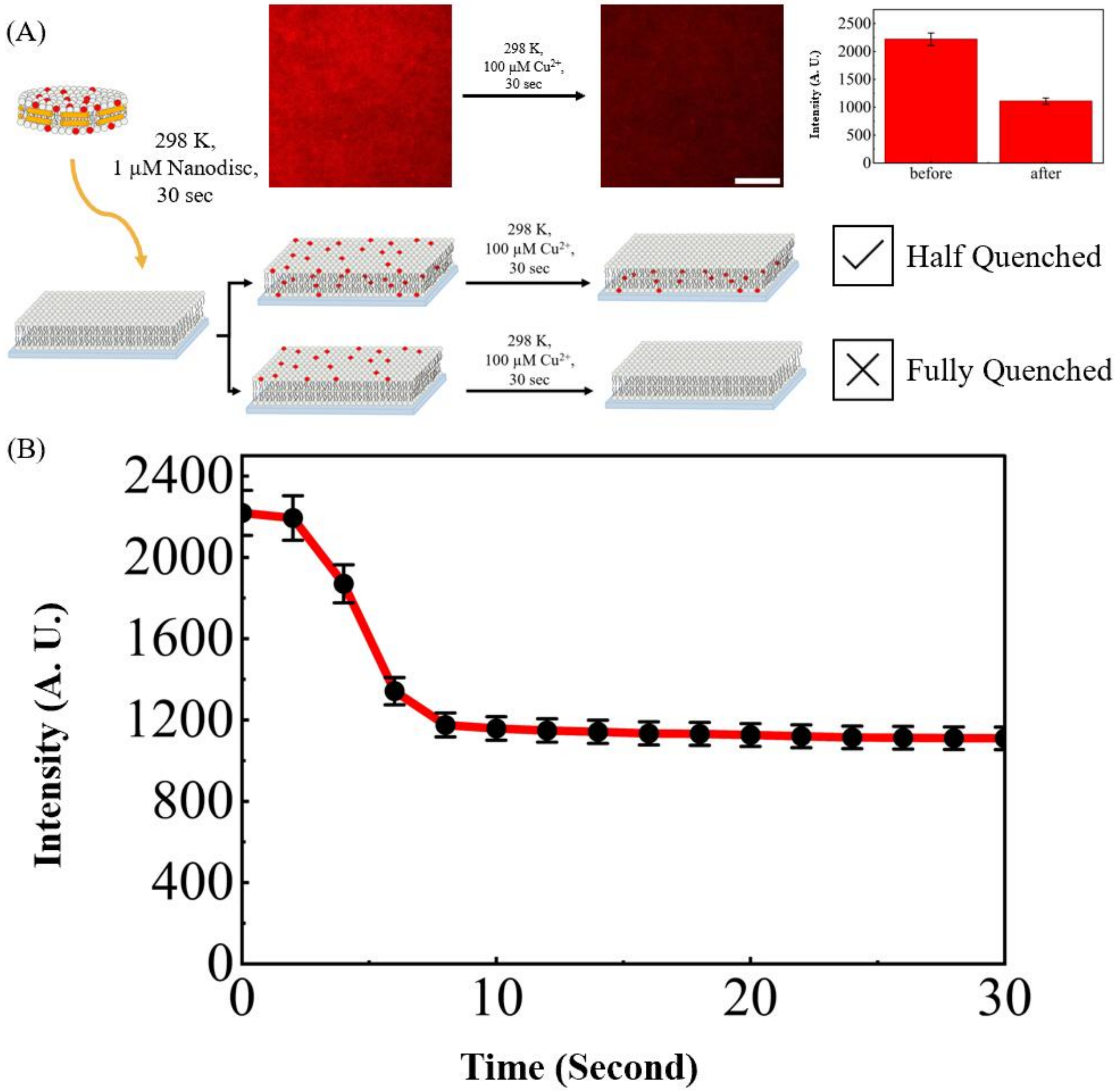


**Figure S1.** Cu(II)-mediated quenching experiments confirming transbilayer lipid integration into supported lipid bilayers. (A) Schematic representation and TIRF images of the fluorescence quenching assay. Following nanodisc-mediated delivery of Texas Red-labeled lipids into the SLB, $Cu^{2+}$ ions (introduced as $CuSO_4$) are added to the bulk solution as a quenching agent. Because the quencher selectively accesses the solvent-exposed upper leaflet of the SLB, the observed ~50% reduction in total fluorescence intensity—as shown in the adjacent plot—demonstrates that the delivered lipids are evenly distributed across both the upper and lower leaflets of the bilayer. (B) Real-time fluorescence intensity traces tracking the quenching process. Upon the addition of 10 mM $Cu^{2+}$ to the bulk solution at t = 0 s, the fluorescence signal rapidly decreases by half within 10 s, confirming the selective quenching of the upper leaflet.

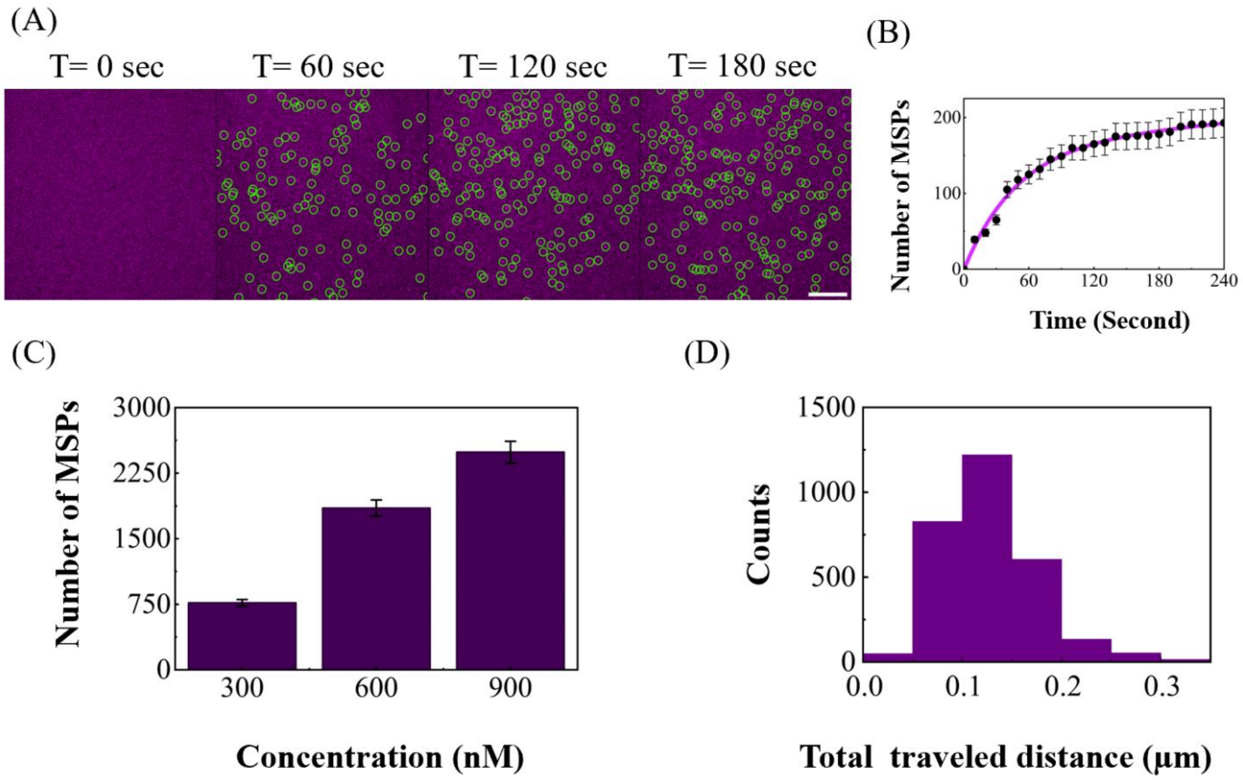


**Figure S2.** Single-molecule tracking and spatiotemporal dynamics of Membrane Scaffold Proteins (MSP) during nanodisc-mediated lipid delivery. (A) Single-molecule TIRF imaging of Alexa Fluor 647-labeled MSP during lipid delivery to the SLB from t = 0 to 180 s. A concentration of [OOO] nM of nanodiscs containing the labeled MSP was added to the SLB. To enable labeling, a cysteine residue was engineered at the C-terminus of the MSP, allowing for fluorophore conjugation via maleimide chemistry. The images demonstrate a rapid increase in the number of MSP molecules on the bilayer over time. Scale bar = 10 μm. (B) Real-time quantification of the number of MSP molecules accumulating on the SLB within the imaging field corresponding to part A. (C) Concentration-dependent accumulation of MSP. The number of MSP molecules in a fixed 80 μm × 80 μm field of view increases as a function of the nanodisc concentration. Measurements were taken following a 10-min incubation of the nanodiscs with the SLB, immediately followed by a buffer wash. (D) Histogram of the total traveled distance for individual MSP molecules. The distribution indicates that most MSP molecules are relatively immobile, suggesting they traverse the SLB and become sterically trapped on the underlying glass substrate.

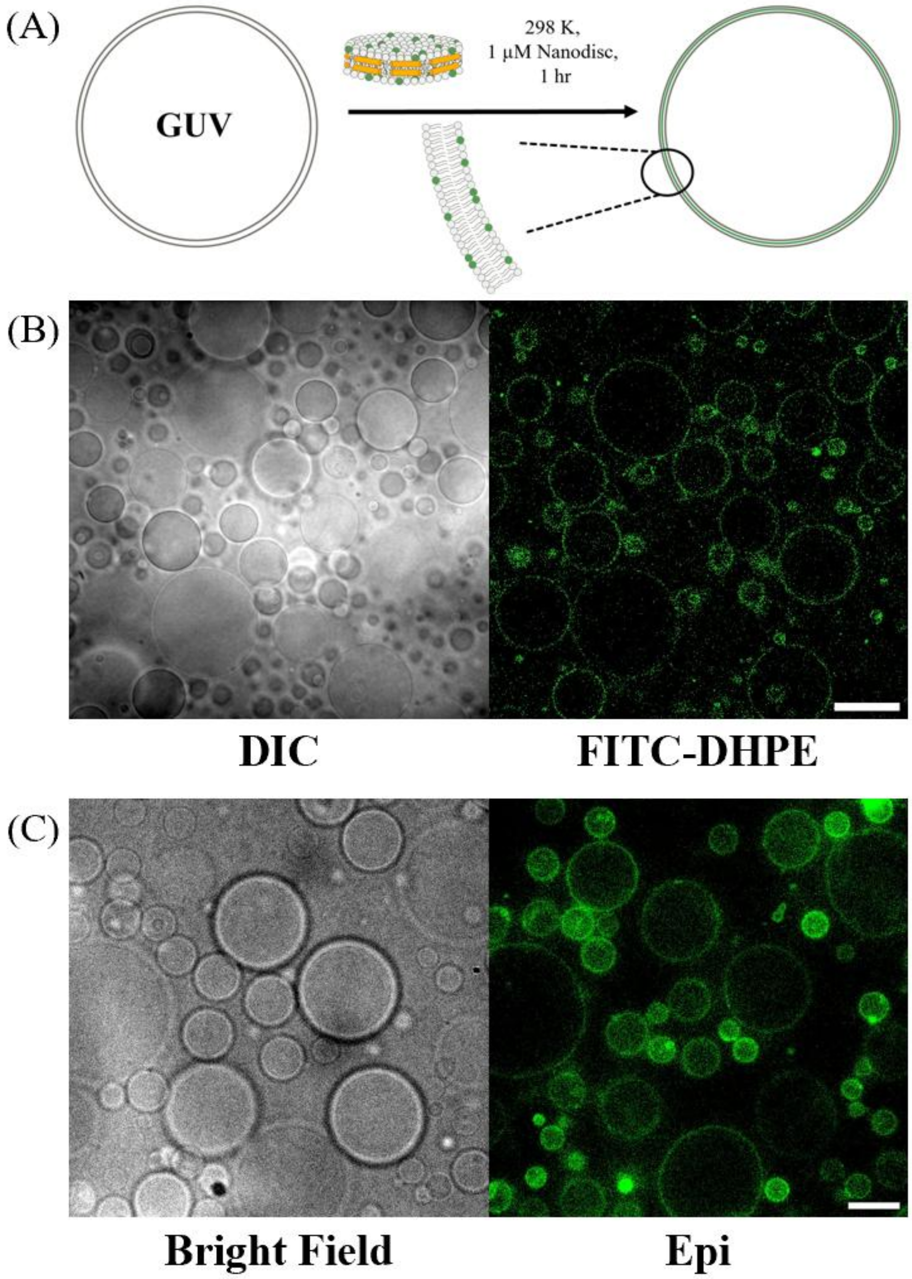


**Figure S3.** MSP nanodisc-mediated delivery of fluorescent lipid cargoes into giant unilamellar vesicle (GUV) model membranes. (A) Schematic illustration of the nanodisc-mediated lipid delivery process. MSP nanodiscs composed of 96 mol% DOPC and 4 mol% FITC-DHPE are introduced to the GUVs, facilitating the targeted unloading and integration of the fluorescent cargo lipids into the vesicle membrane. (B) Representative differential interference contrast (DIC) and corresponding confocal fluorescence images of the GUVs following a 60 min incubation with the nanodiscs.

To enable stable imaging, the GUVs were tethered to the glass substrate via biotin-streptavidin interactions. Confocal optical sectioning reveals a distinct fluorescent ring corresponding to the vesicular cross-section, confirming the successful structural integration of the FITC-DHPE lipids into the GUV bilayer. (C) Representative bright-field and epifluorescence images of the nanodisc-treated GUVs. The wide-field imaging confirms robust, uniform FITC fluorescence across the entire observed population, further demonstrating the high delivery efficiency of the lipid cargoes into the target membranes.

(A)

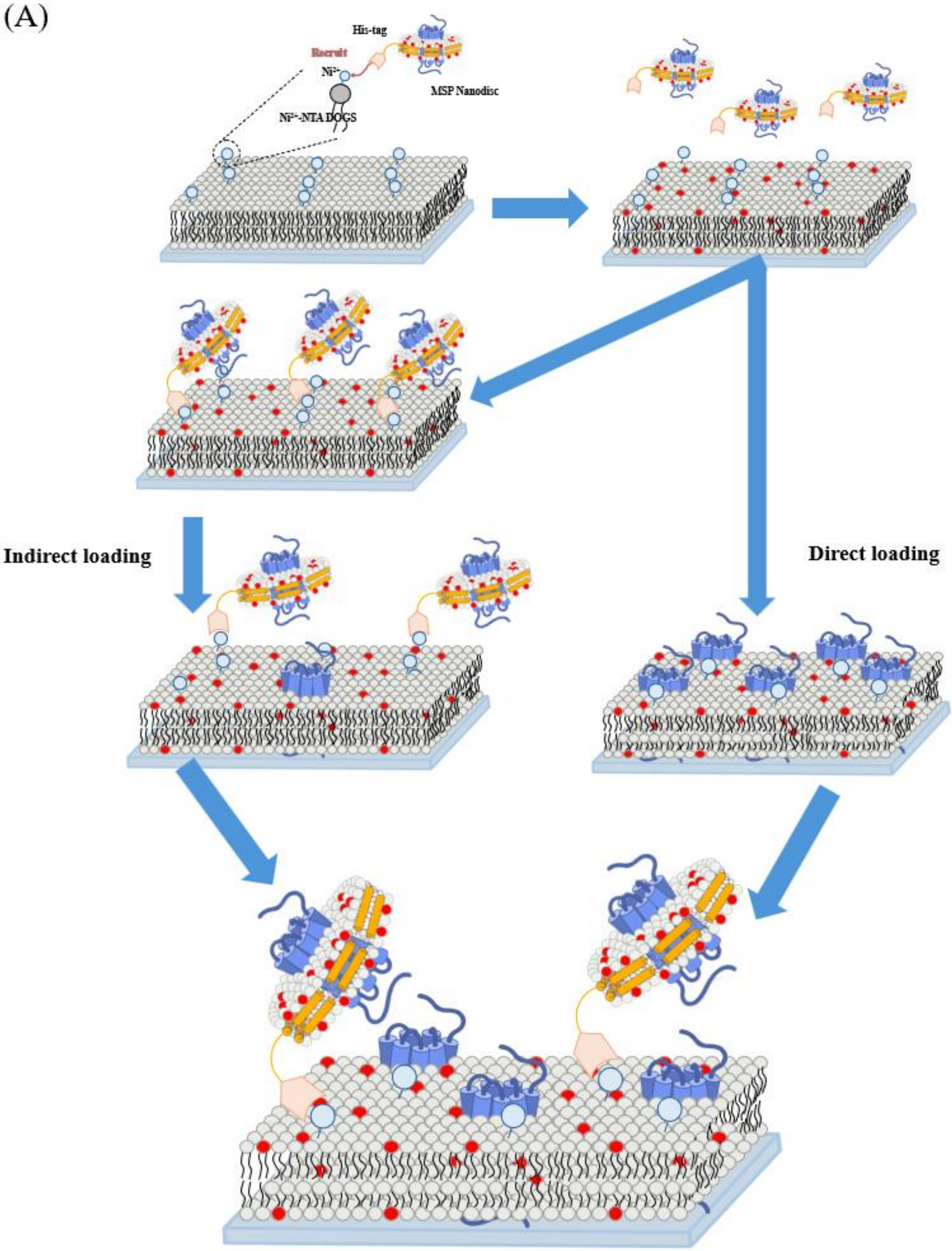
His-tag
Recruit
Ni²⁺
MSP Nanodisc
Ni²⁺-NTA DOGS
Indirect loading
Direct loading

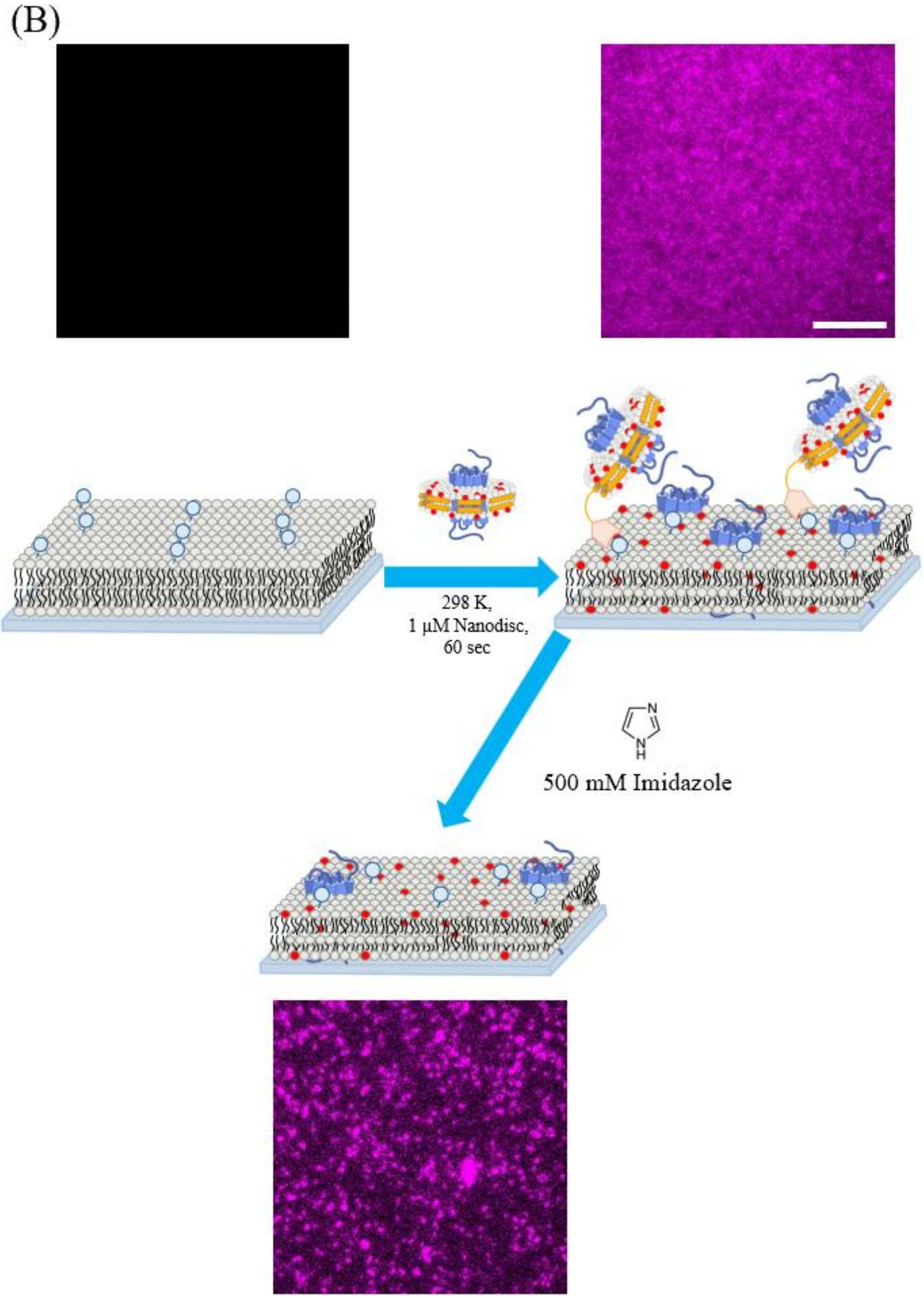


**Figure S4.** His-tag mediated nanodisc recruitment enhances the delivery efficiency of membrane proteins to supported lipid bilayers. (A) Schematic illustration comparing direct and indirect nanodisc loading mechanisms. The polyhistidine tag (His-tag) on the Membrane Scaffold Protein (MSP)—originally engineered for purification—can be repurposed to physically recruit the nanodisc to the SLB surface (indirect loading). In contrast to direct loading, where stochastic collisions between freely diffusing nanodiscs and the bilayer mediate cargo release, indirect loading tethers the nanodisc to the membrane. This forced physical proximity significantly increases the probability of successful lipid and membrane protein transfer. (B) Selective retention of fully integrated BsYetJ following imidazole-mediated washing. The upper TIRF images show strong fluorescence resulting from both intact, recruited nanodiscs and successfully delivered Alexa Fluor 647-labeled BsYetJ following a 60 s incubation. Subsequent washing with a 500 mM imidazole solution competitively dissociates the intact, His-tagged nanodiscs from the membrane surface, ensuring the remaining fluorescence signal originates exclusively from the Alexa Fluor 647-labeled BsYetJ successfully embedded within the bilayer. Scale bar = 10 μm.

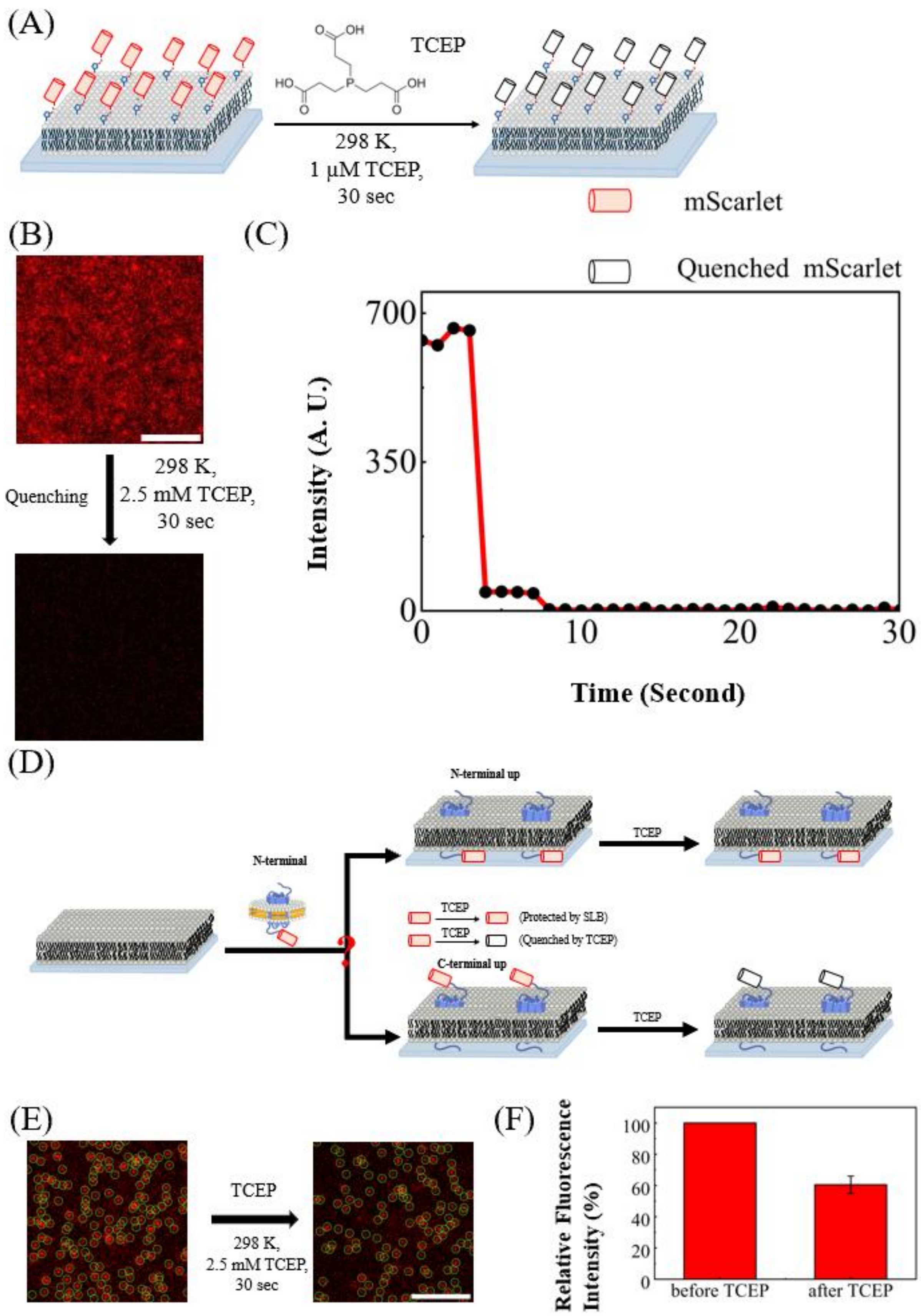


**Figure S5.** TCEP-mediated fluorescence quenching to determine the membrane insertion topology of nanodisc-delivered BsYetJ. (A) Schematic illustrating that TCEP fully quenches mScarlet anchored to the upper leaflet of SLB via maleimide chemistry. This validates TCEP as an effective, membrane-impermeable tool for determining mScarlet localization in parts D–F. (B) Representative TIRF images of SLB-anchored mScarlet before and after the addition of TCEP. (C) A real-time fluorescence intensity trace demonstrating that the fluorescence signal from upper-leaflet mScarlet is nearly completely quenched within 10 s. (D) Schematic of the topology assay. The C-terminally mScarlet-tagged BsYetJ construct is delivered into the SLB via MSP nanodiscs. TCEP is subsequently used as a quenching agent to identify the location of the mScarlet tag, thereby revealing the transmembrane orientation of BsYetJ. (E) Single-molecule TIRF images of nanodisc-delivered BsYetJ-mScarlet. The images show that approximately half of the mScarlet molecules are quenched upon TCEP addition. (F) Quantitative plot of the relative ratio of fluorescent mScarlet molecules before and after TCEP quenching. The ~50% reduction indicates that both insertion topologies of BsYetJ-mScarlet are roughly equally populated within the SLB.

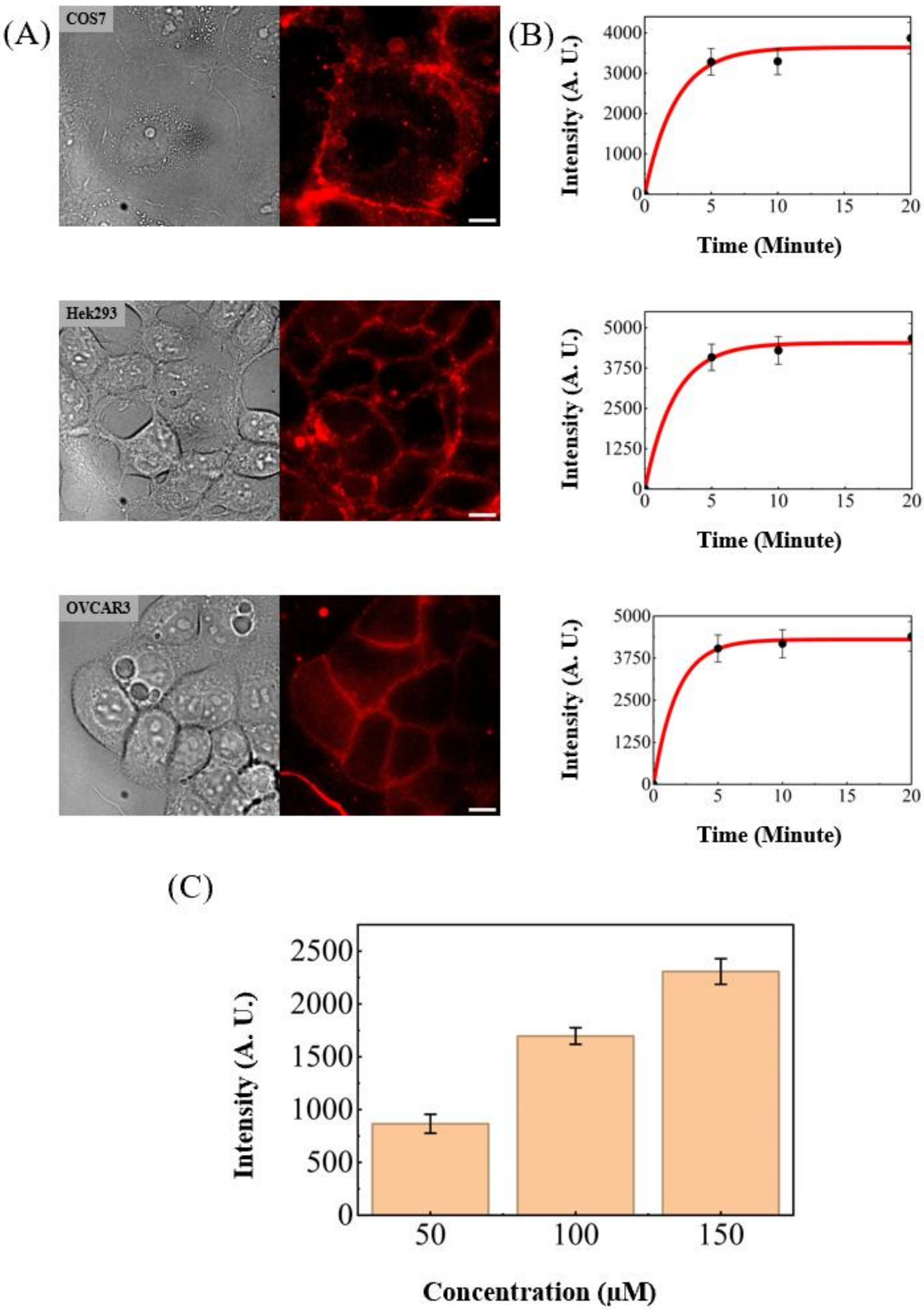


**Figure S6.** Rapid and versatile nanodisc-mediated delivery of cargo lipid into the plasma membrane of diverse mammalian cell lines. (A) Representative bright-field and epifluorescence images of the treated cells. The fluorescence signals clearly localize to the cellular plasma membrane, demonstrating successful integration. (B) The corresponding time-course quantification reveals the rapid membrane accumulation of lipids, with the Texas Red DHPE lipid fluorescence intensity reaching a plateau in less than 10 minutes. (C) Concentration dependence of lipid delivery assessed at MSP nanodisc concentrations of 50, 100, and 150 nM for HEK293 cells. Nanodiscs loaded with 4% Texas Red-DHPE and 96% DOPC were added to the cells in imaging buffer. Following an incubation period of 10 min, peripheral fluorescence intensities were quantified from epifluorescence images and plotted.

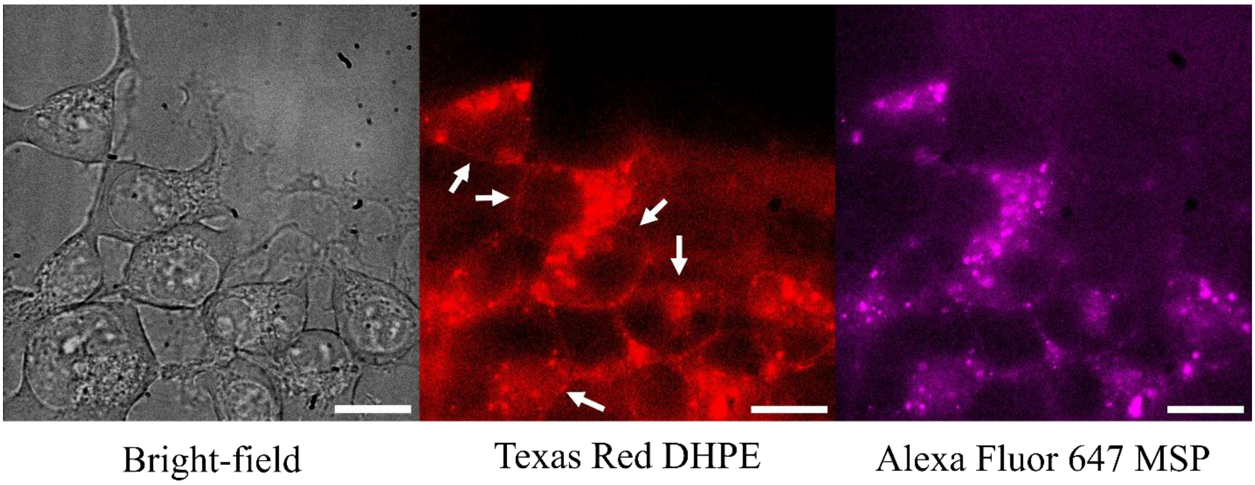


**Figure S7.** Epifluorescence tracking of dual-labeled nanodiscs reveals differences in spatial distribution between lipids and MSP in HEK293 cells. Representative bright-field and epifluorescence images of HEK293 cells following a 20-min incubation at 37 °C with dual-labeled nanodiscs, followed by an imaging buffer wash to remove unbound materials. The nanodiscs are composed of Alexa Fluor 647-labeled MSP (AF647-MSP), 96 mol% DOPC, and 4 mol% Texas Red-DHPE. The imaging indicates the delivery of Texas Red-DHPE lipids to the plasma membrane, as shown by fluorescence at the cell periphery (indicated by white arrows), alongside some intracellular lipid fluorescence attributed to cellular trafficking and membrane internalization during the incubation. By contrast, the AF647-MSP exhibits a different localization pattern, accumulating more predominantly within the cytosol with a less pronounced peripheral signal. This observation is consistent with the results from the experiments using the supported lipid bilayer (see Figure S4), where the AF647-MSP was observed to traverse the bilayer and become sterically trapped on the underlying glass substrate.

**(A)**

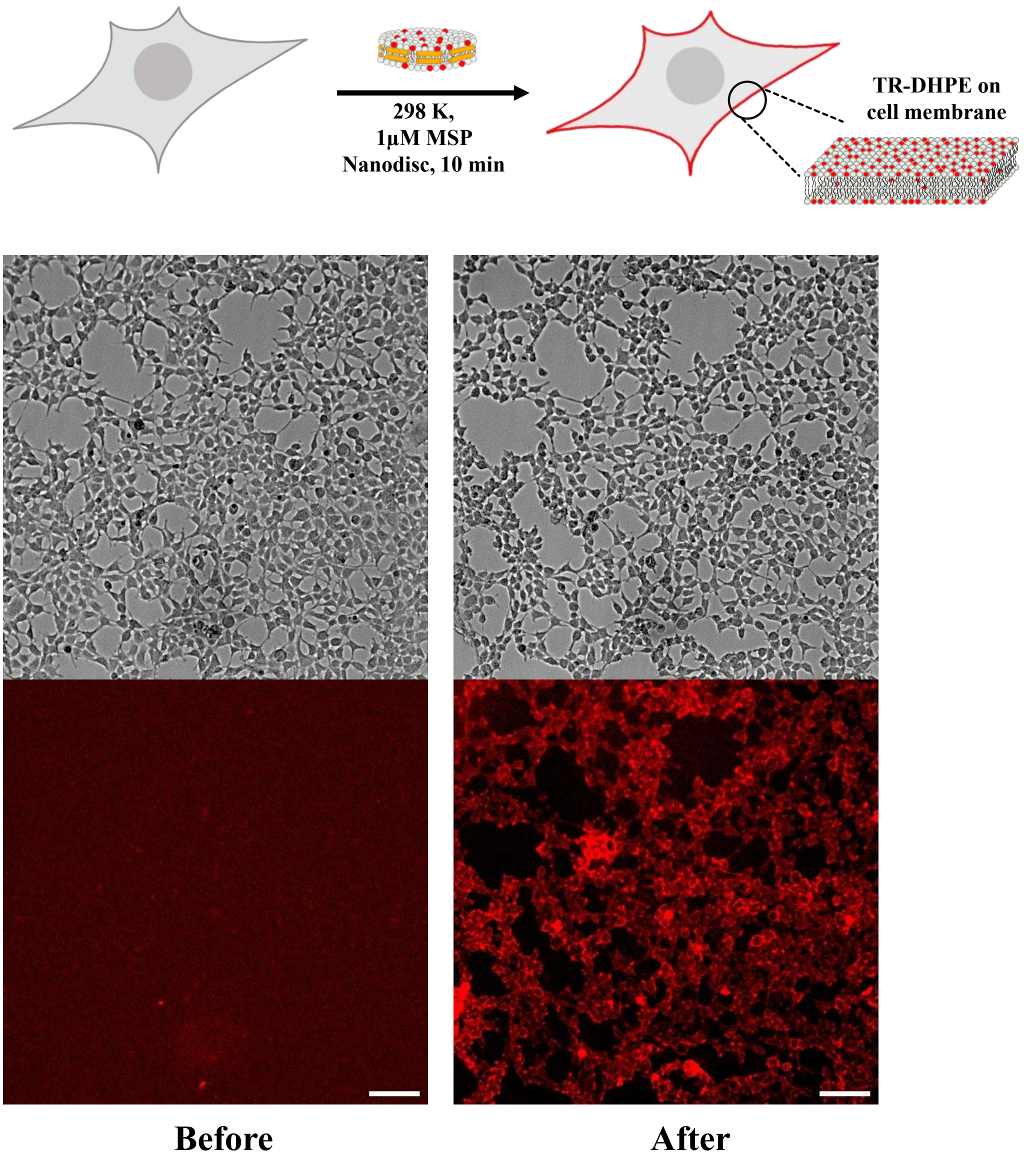

298 K,
1μM MSP
Nanodisc, 10 min
TR-DHPE on
cell membrane
Before
After

(B)

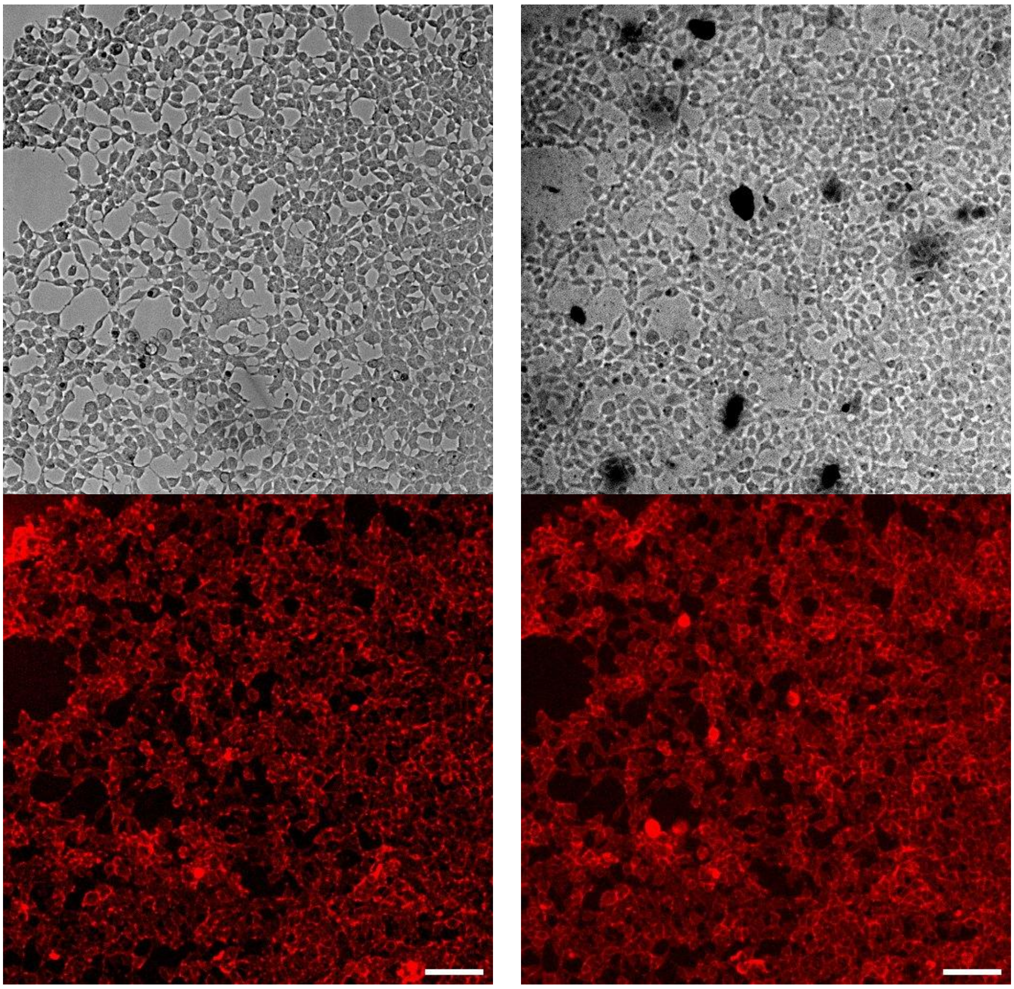

Before Trypan Blue　　After Trypan Blue

(C)

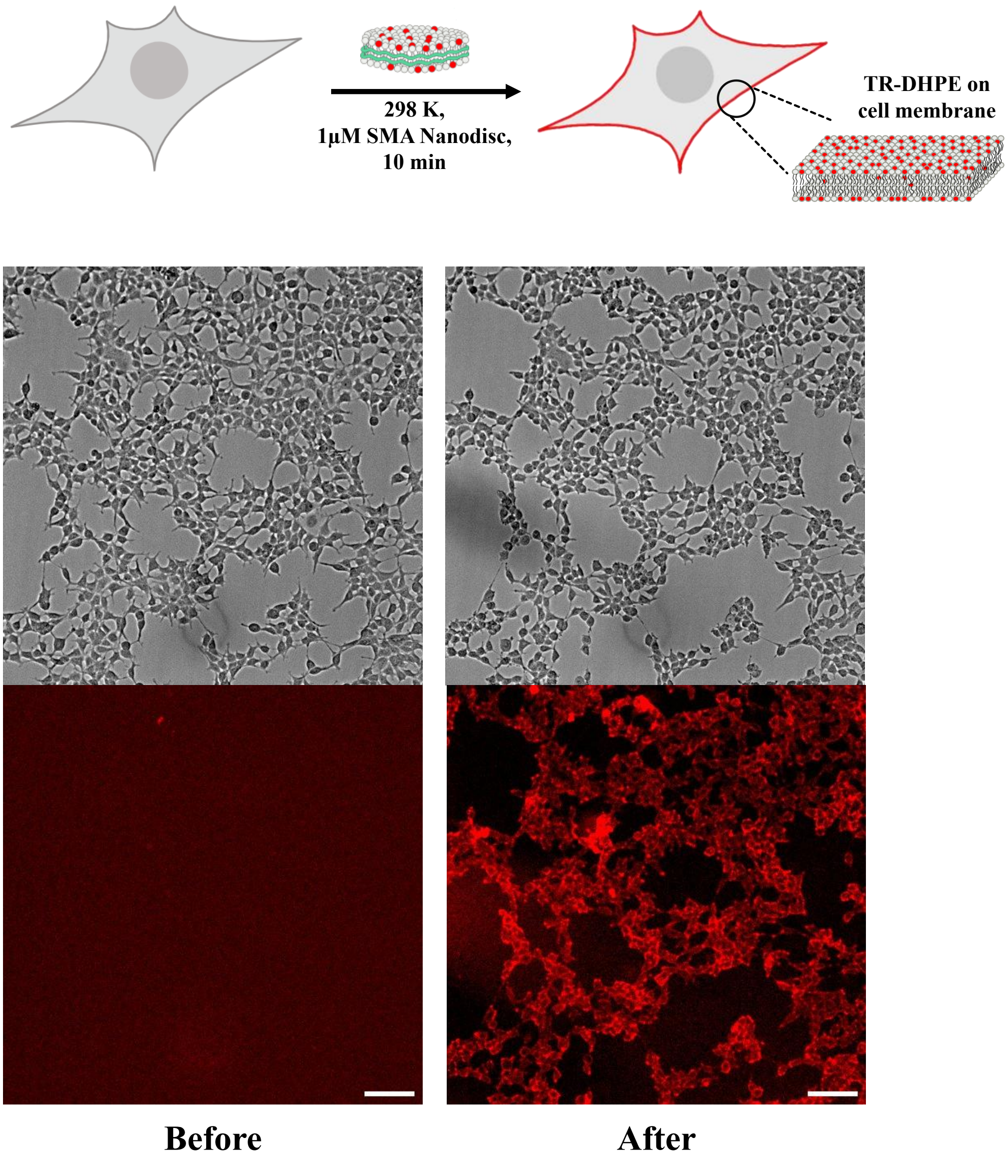
298 K,
1μM SMA Nanodisc,
10 min
TR-DHPE on
cell membrane
Before
After

(D)

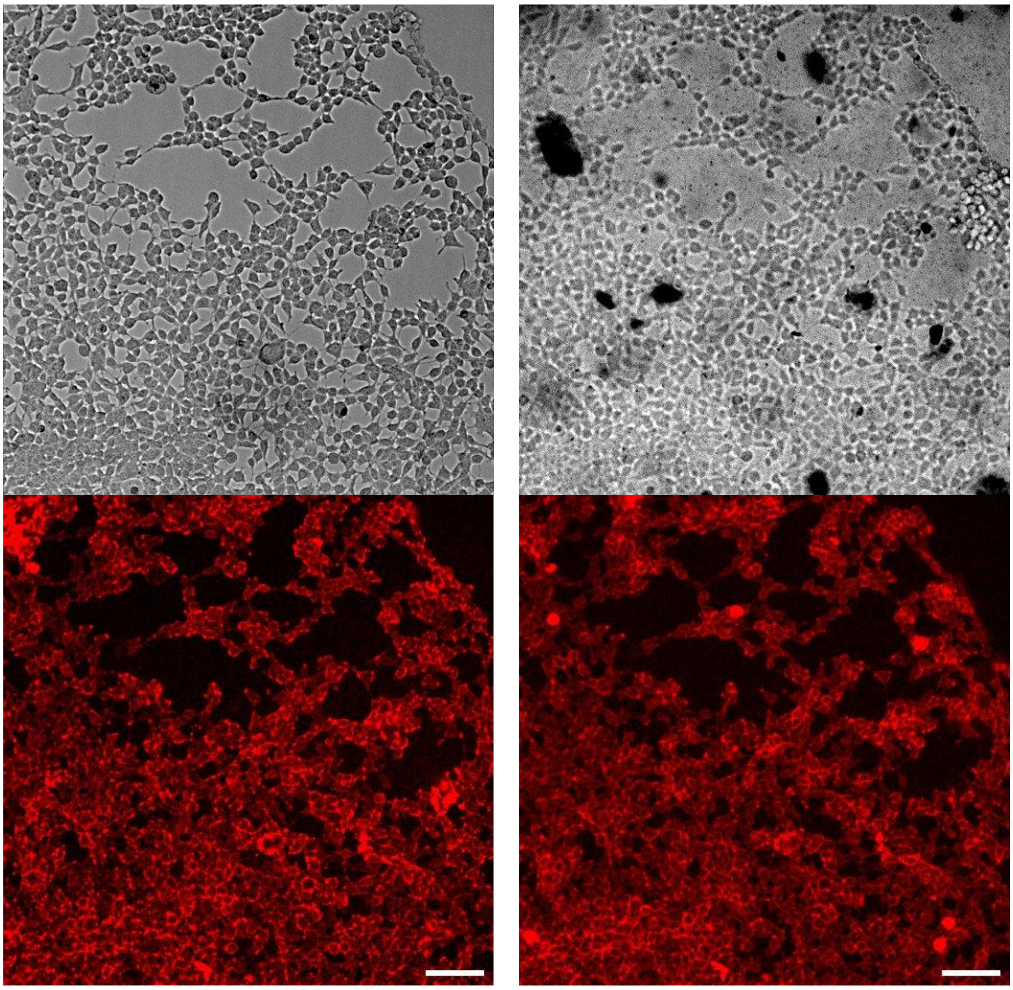

Before Trypan Blue After Trypan Blue

**Figure S8.** Assessment of nanodisc-mediated lipid delivery efficiency and cellular viability. (A) Schematic illustration depicting the rapid incorporation of Texas Red-DHPE into the plasma membranes of naive cells following the addition of loaded MSP nanodiscs. Corresponding low-magnification epifluorescence and bright-field images demonstrate that nearly all cells exhibit robust fluorescence, indicating ~100% lipid delivery efficiency. (B) Viability assessment of the cells from (A) using a Trypan Blue exclusion assay. The retention of cell adhesion to the coverslip and the absence of Trypan Blue intracellular staining following a buffer wash confirm near 100% cell viability during MSP nanodisc-mediated delivery. The dark particles visible in the bright-field image are undissolved Trypan Blue powder. (C) Parallel lipid delivery experiments utilizing SMA-type nanodiscs loaded with Texas Red-DHPE. Low-magnification imaging similarly reveals an approximate 100% lipid delivery efficiency. (D) Trypan Blue exclusion assay for the cells treated in (C), confirming that SMA nanodisc-mediated delivery also maintains near 100% cell viability and substrate adhesion. As in (B), the dark particles observed in the bright-field image represent residual Trypan Blue powder. Scale bars: 100 μm.

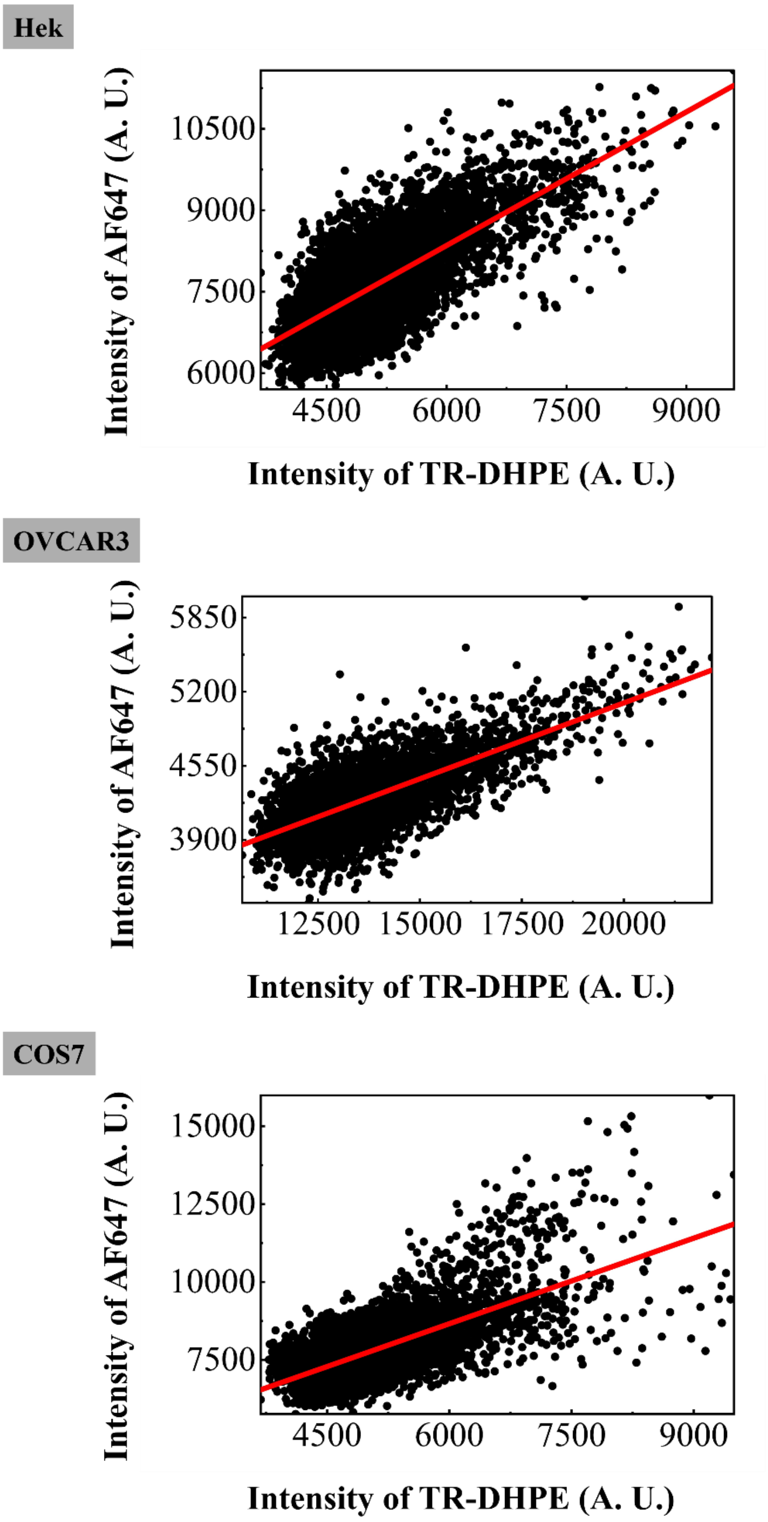


**Figure S9.** Quantitative Pearson colocalization analysis of nanodisc-delivered lipid and protein cargoes. Representative 2D cytofluorograms illustrating the spatial fluorescence intensity correlation between the Texas Red-DHPE and Alexa Fluor 647-labeled BsYetJ channels across the imaging field (using 4 × 4 pixel binning). Quantitative Pearson colocalization analysis yields strong positive correlation coefficients (r) across all tested cell lines (r = 0.711 for HEK, 0.721 for OVCAR3, and 0.676 for COS7). These values mathematically corroborate the synchronized co-delivery and integration of both the lipid and protein cargoes into the cellular plasma membrane.

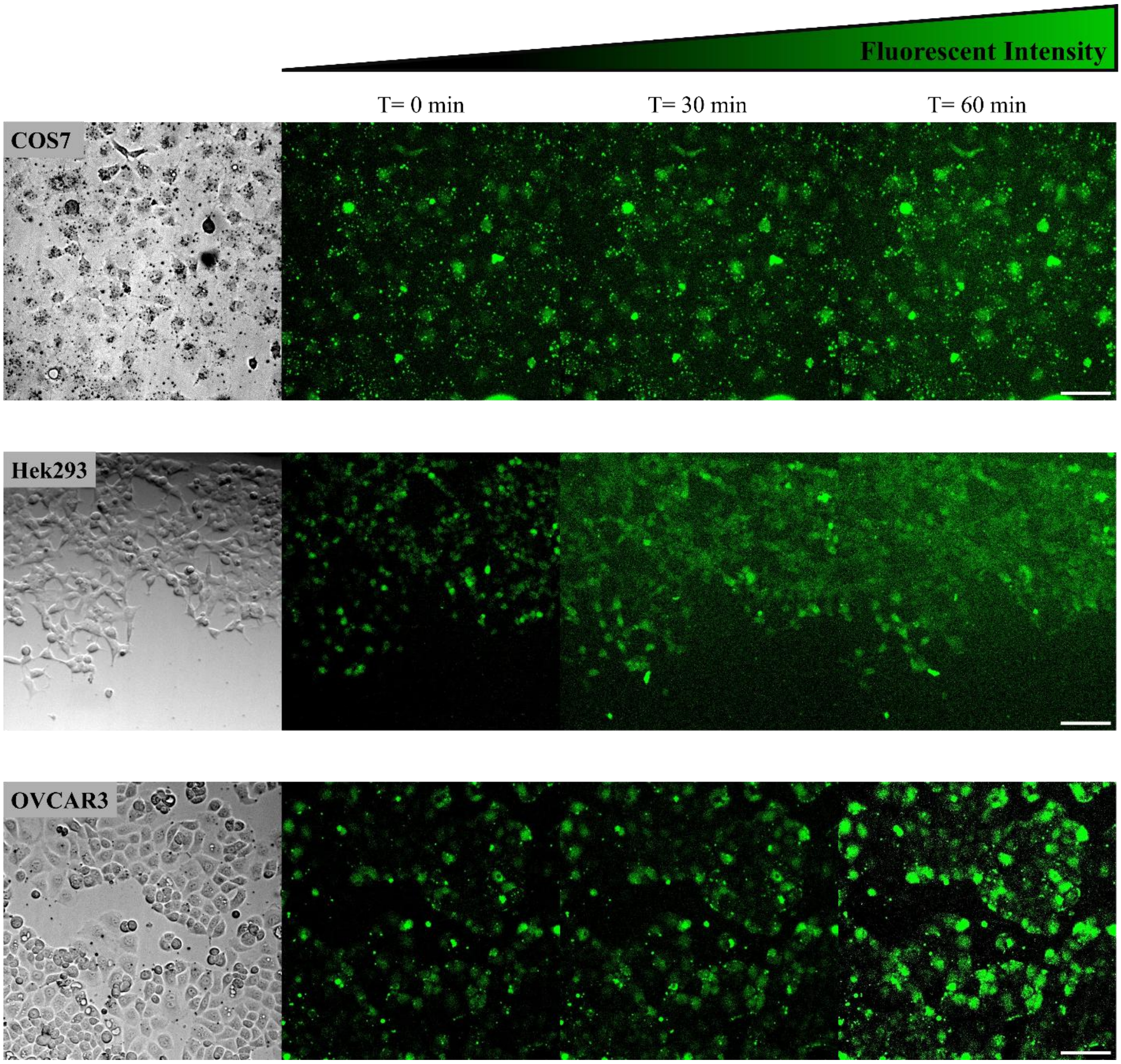


**Figure S10.** High-throughput epifluorescence imaging of nanodisc-mediated BsYetJ calcium conductance across diverse mammalian cell lines. Rows (from top to bottom) present COS7, HEK293, and OVCAR3 cell lines, respectively. The leftmost column displays representative bright-field images of the cells. The subsequent columns exhibit the corresponding Fluo-8 epifluorescence images, illustrating a progressive, time-dependent increase in fluorescence intensity at t = 0, 30, and 60 min following the introduction of $Ca^{2+}$ into the extracellular medium. The quantitative analysis of the averaged temporal fluorescence intensity changes for each respective cell line is detailed in Figure 4E–G.

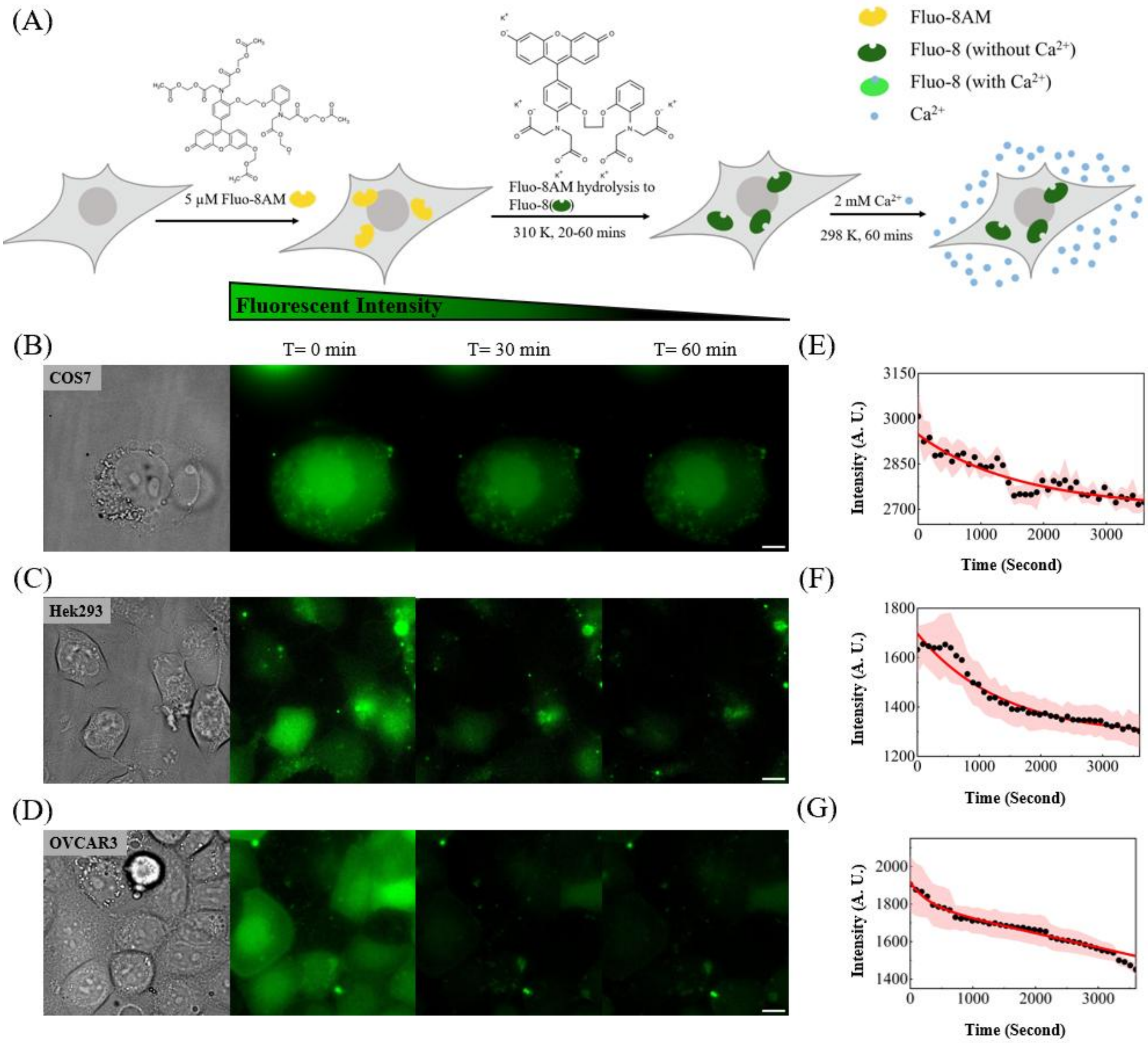


**Figure S11.** Negative control assays for intracellular calcium imaging in the absence of nanodisc-delivered BsYetJ. (A) Schematic illustration of the control intracellular calcium imaging assay. The calcium indicator Fluo-8 AM is introduced into the cells, but this step is not followed by the nanodisc-mediated delivery of BsYetJ. The cells are incubated in a $Ca^{2+}$-free medium for 45 min to allow for the complete intracellular hydrolysis of Fluo-8 AM to Fluo-8, ensuring the basal fluorescence intensity reaches a stable plateau. Upon the addition of $Ca^{2+}$ to the extracellular medium, no intracellular calcium influx occurs; instead, a progressive decrease in fluorescence intensity is observed due to the natural photobleaching of the Fluo-8 dye over time. (B–D)

Representative bright-field transmission images (left) and corresponding time-lapse Fluo-8 epifluorescence frames at t = 0, 30, and 60 min (right), tracking the baseline intracellular fluorescence dynamics across the different mammalian cell lines (COS7, HEK293, and OVCAR3). (E–G) Quantitative time-course fluorescence intensity traces demonstrating the gradual photobleaching of Fluo-8. To generate the robust statistical data presented in these panels, large field-of-view epifluorescence microscopy was utilized as a high-throughput assay to average data from multiple cells across various imaging fields (as detailed in Figure S11).

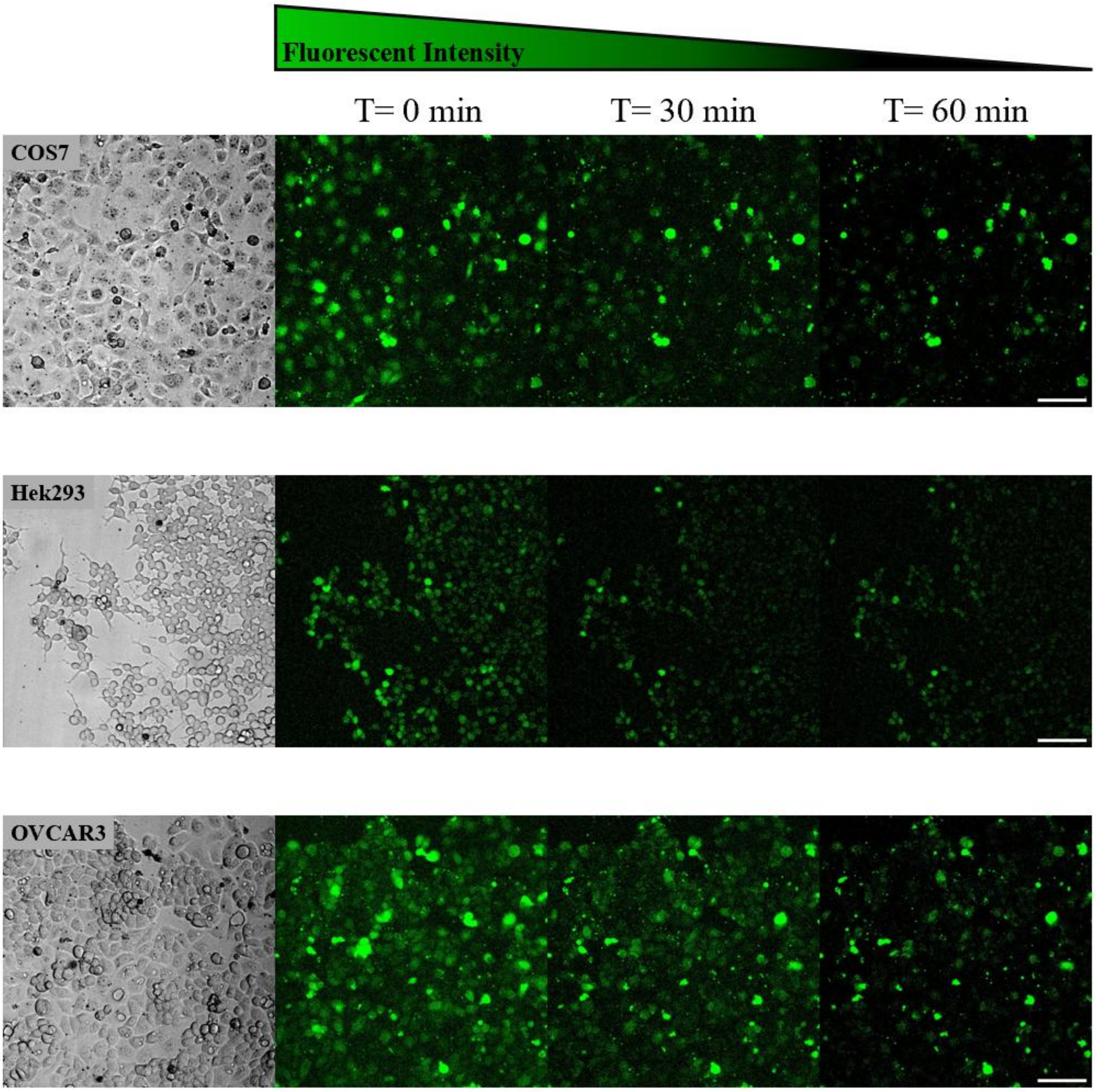


**Figure S12.** High-throughput epifluorescence imaging of negative control assays for intracellular calcium dynamics. The calcium indicator Fluo-8 AM is introduced into the cells, but this step is not followed by the nanodisc-mediated delivery of BsYetJ, serving as a negative control for the functional assays (see Figure S6). Rows (from top to bottom) present the COS7, HEK293, and OVCAR3 cell lines, respectively. The leftmost column displays representative bright-field images of the cells. The subsequent columns exhibit the corresponding Fluo-8 epifluorescence images. Because the BsYetJ channel is absent, no intracellular calcium influx occurs; instead, a progressive decrease in fluorescence intensity is observed due to the natural photobleaching of the Fluo-8 dye at t = 0, 30, and 60 min following the introduction of $Ca^{2+}$ into the extracellular medium. The quantitative analysis of the averaged temporal fluorescence intensity changes for each respective cell line is detailed in Figure S6E–G.

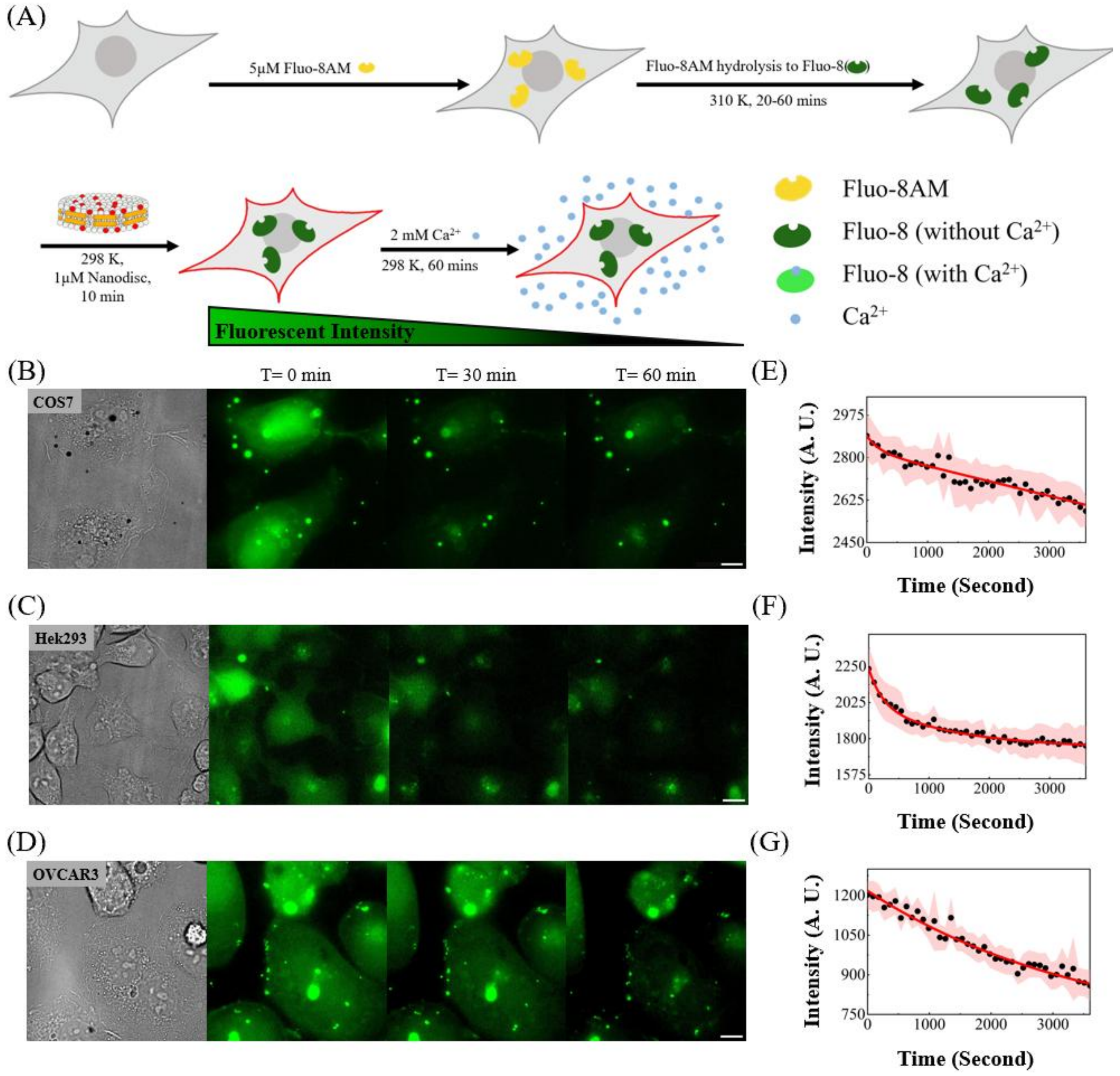


**Figure S13.** Vehicle control assays confirming no apparent intracellular calcium influx is induced by empty nanodiscs. (A) Schematic of the vehicle control assay. Following the introduction of the Fluo-8 AM calcium indicator, cells are treated with empty nanodiscs (lacking the BsYetJ channel). This rules out the possibility that the nanodisc delivery vehicle itself perturbs the cell membrane. After a 45-min incubation in $Ca^{2+}$-free medium to allow for dye hydrolysis and baseline stabilization, extracellular $Ca^{2+}$ is introduced. Crucially, the presence of the empty nanodiscs does not trigger calcium leakage; instead, only a progressive decrease in fluorescence—indicative of expected baseline photobleaching—is observed. (B–D) Representative bright-field (left) and time-lapse Fluo-8 epifluorescence images (right) of COS7, HEK293, and OVCAR3 cells treated with empty nanodiscs. The frames at t = 0, 30, and 60 min confirm the sustained absence of apparent calcium influx across all tested cell lines. (E–G) Normalized time-course fluorescence intensity traces illustrating the expected gradual photobleaching of Fluo-8. Consistent with the functional assays, robust statistical averaging was achieved by utilizing large field-of-view, high-throughput epifluorescence microscopy across multiple imaging fields (detailed in Figure S13).

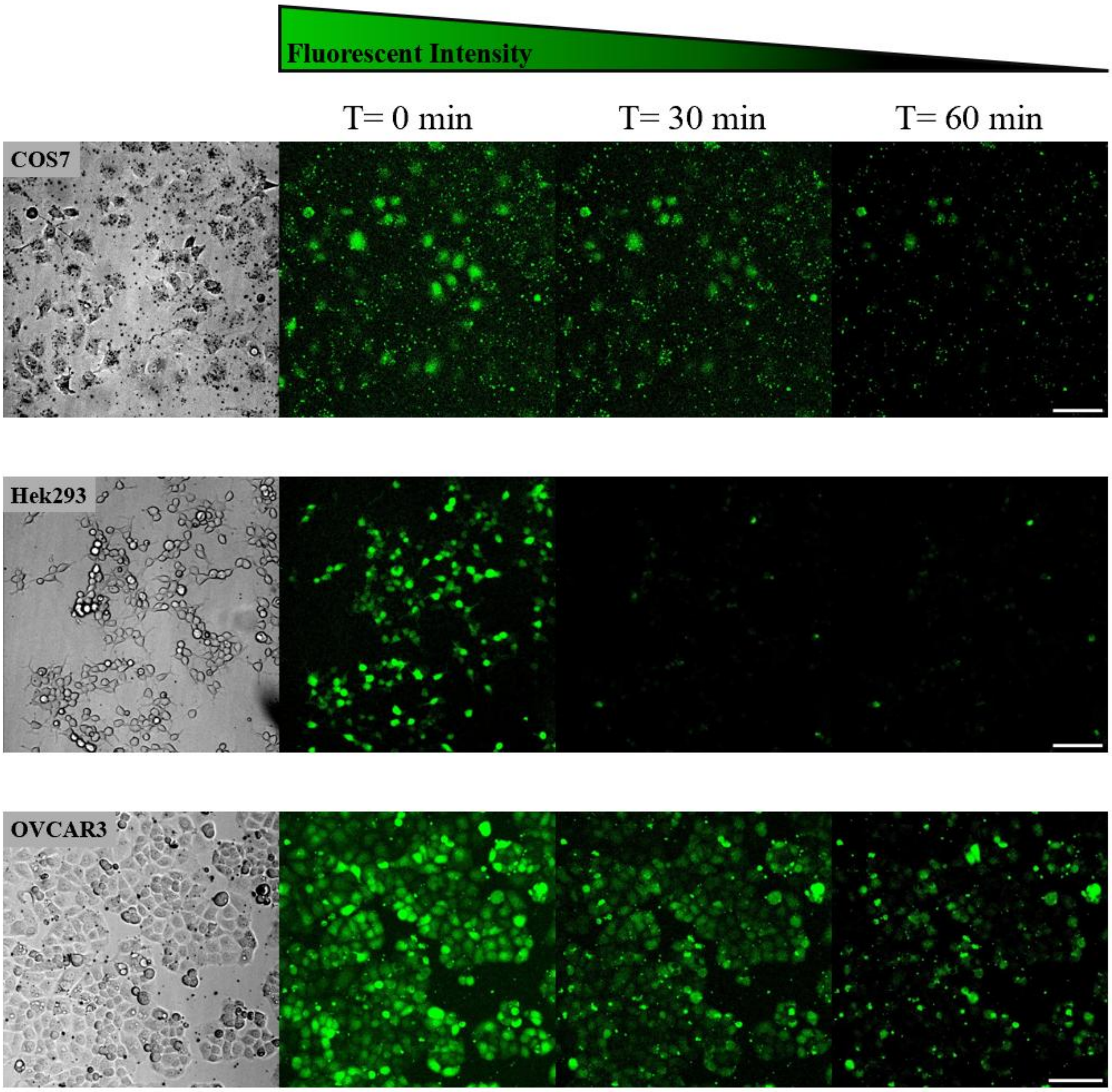


**Figure S14.** High-throughput epifluorescence imaging of vehicle control assays confirming no apparent calcium influx is induced by empty nanodiscs. Following the introduction of the Fluo-8 AM calcium indicator, the cells are treated with empty nanodiscs (lacking the BsYetJ channel), serving as a critical vehicle control for the functional assays (see Figure S8). Rows (from top to bottom) present the COS7, HEK293, and OVCAR3 cell lines, respectively. The leftmost column displays representative bright-field images of the cells, while the subsequent columns exhibit the corresponding Fluo-8 epifluorescence images. Because the BsYetJ channel is absent, no apparent intracellular calcium influx occurs; instead, a progressive decrease in fluorescence intensity is observed due to the natural photobleaching of the Fluo-8 dye at $t = 0$, 30, and 60 min following the introduction of $Ca^{2+}$ into the extracellular medium. The quantitative analysis of the averaged temporal fluorescence intensity changes for each respective cell line is detailed in Figure S8E–G.

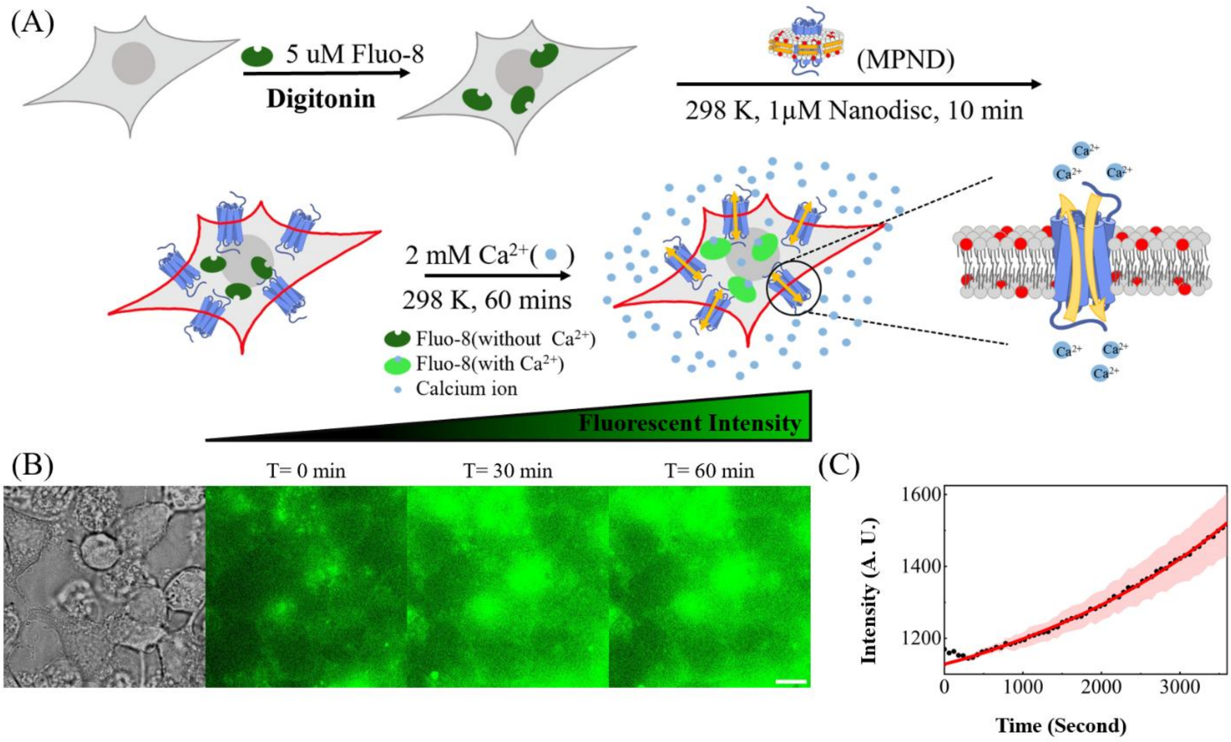


**Figure S15.** Comparative validation of intracellular calcium imaging utilizing detergent-mediated cytosolic loading of active Fluo-8. (A) Schematic illustration of the comparative calcium sensing assay. While standard intracellular calcium imaging relies on the esterase-mediated hydrolysis of membrane-permeable Fluo-8 AM to generate the active, carboxyl-containing Fluo-8 dye, this alternative assay utilizes detergent-mediated membrane permeabilization for the direct cytosolic loading of active Fluo-8. This initial loading step is subsequently followed by the nanodisc-mediated delivery and integration of both full-length BsYetJ and co-incorporated lipids into the cellular plasma membrane. This parallel series of validation experiments was performed to facilitate direct comparison with alternative protocols established by other research groups. Upon the addition of $Ca^{2+}$ to the extracellular medium, influx through the reconstituted channels is captured via chelation by the Fluo-8 indicator, resulting in a robust increase in fluorescence intensity. (B) Representative bright-field transmission (left) and corresponding time-lapse epifluorescence images (right) of the HEK293 cells. The frames at $t = 0$, 30, and 60 min visually track the intracellular calcium accumulation facilitated by the nanodisc-delivered channels following direct Fluo-8 loading. (C) Quantitative time-course fluorescence intensity traces. The distinct, time-dependent increase in cytosolic $Ca^{2+}$ levels mediated by the delivered channels confirms that BsYetJ preserves its native conformation and fully active ion-conductive properties upon nanodisc-mediated reconstitution.

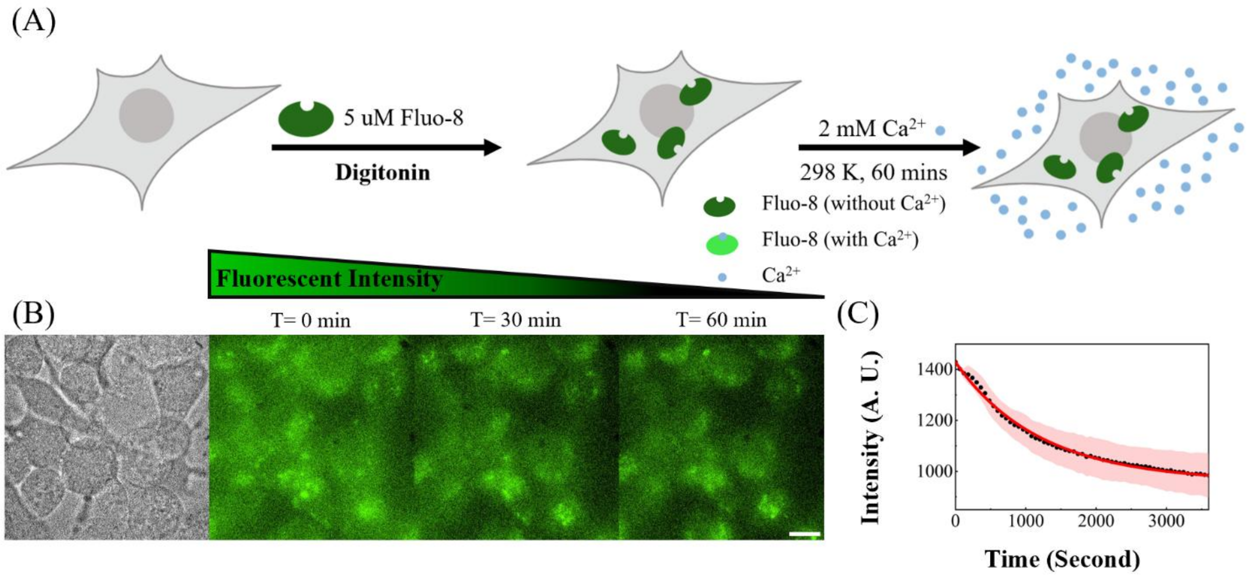


**Figure S16.** Negative control assays for detergent-mediated cytosolic loading of active Fluo-8 in the absence of BsYetJ. (A) Schematic illustration of the control intracellular calcium imaging assay. This alternative assay utilizes detergent-mediated membrane permeabilization for the direct cytosolic loading of active Fluo-8; however, this step is *not* followed by the nanodisc-mediated delivery of BsYetJ. Upon the addition of $Ca^{2+}$ to the extracellular medium, no apparent intracellular calcium influx occurs. Instead, a progressive decrease in fluorescence intensity is observed due to the natural photobleaching of the Fluo-8 dye over time. (B) Representative bright-field transmission (left) and corresponding time-lapse Fluo-8 epifluorescence images (right) of the HEK293 cells. The frames at $t$ = 0, 30, and 60 min visually track the baseline intracellular fluorescence dynamics in the absence of the channel. (C) Quantitative time-course fluorescence intensity traces. The data show a steady decay in signal, confirming the expected gradual photobleaching of Fluo-8 without nanodisc-mediated calcium conductance.

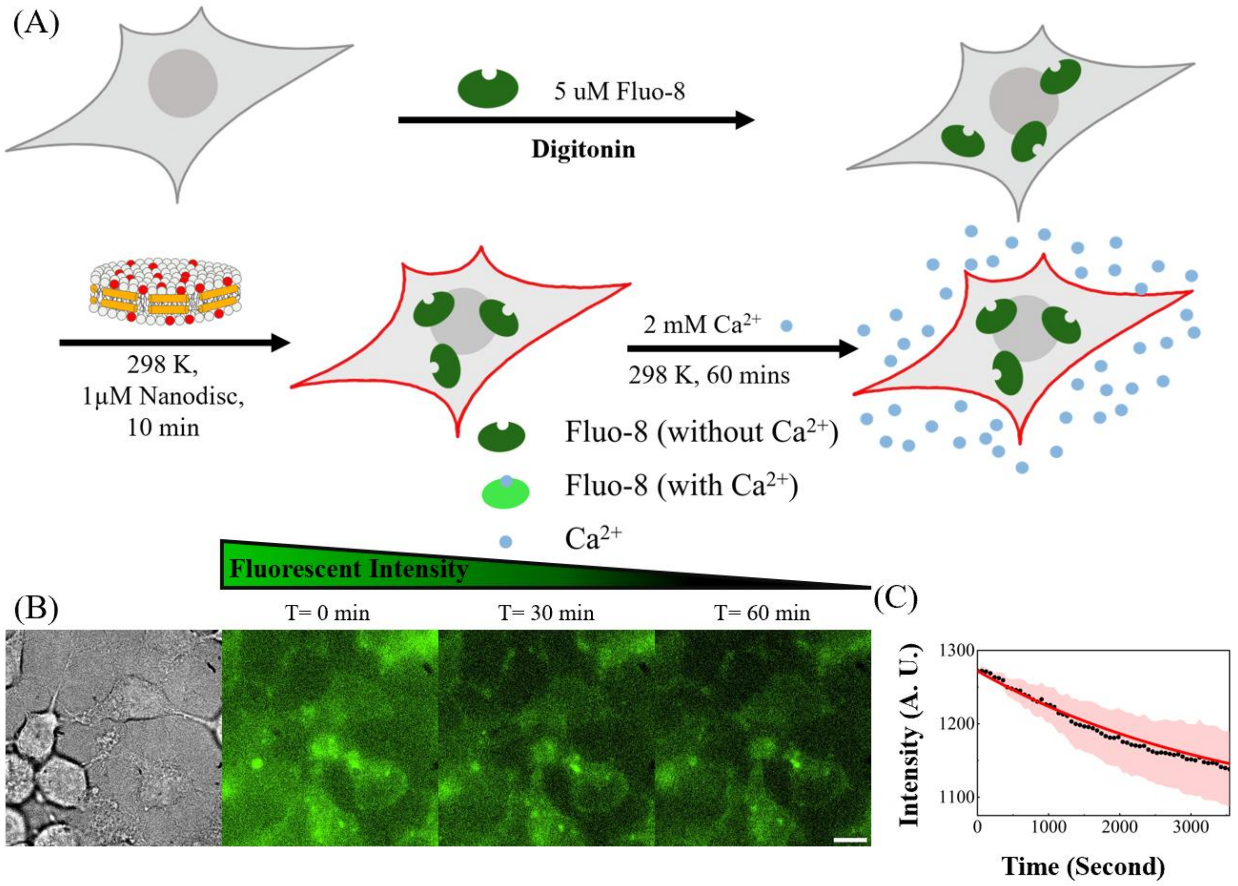


**Figure S17.** Vehicle control assays for detergent-mediated cytosolic loading of active Fluo-8 confirming no apparent intracellular calcium influx is induced by empty nanodiscs. (A) Schematic illustration of the vehicle control assay. Following detergent-mediated permeabilization and the direct cytosolic loading of active Fluo-8, the cells are treated with empty nanodiscs (lacking the BsYetJ channel). Upon the addition of $Ca^{2+}$ to the extracellular medium, no apparent intracellular calcium influx occurs. Instead, a progressive decrease in fluorescence intensity is observed due to the natural photobleaching of the Fluo-8 dye over time. (B) Representative bright-field transmission (left) and corresponding time-lapse Fluo-8 epifluorescence images (right) of the HEK293 cells treated with empty nanodiscs. The frames at $t$ = 0, 30, and 60 min show no observable increase in Fluo-8 fluorescence intensity, confirming the sustained absence of apparent calcium influx. (C) Quantitative time-course fluorescence intensity traces. The data show a steady decay in signal, confirming the expected gradual photobleaching of Fluo-8 in the absence of nanodisc-mediated calcium conductance.

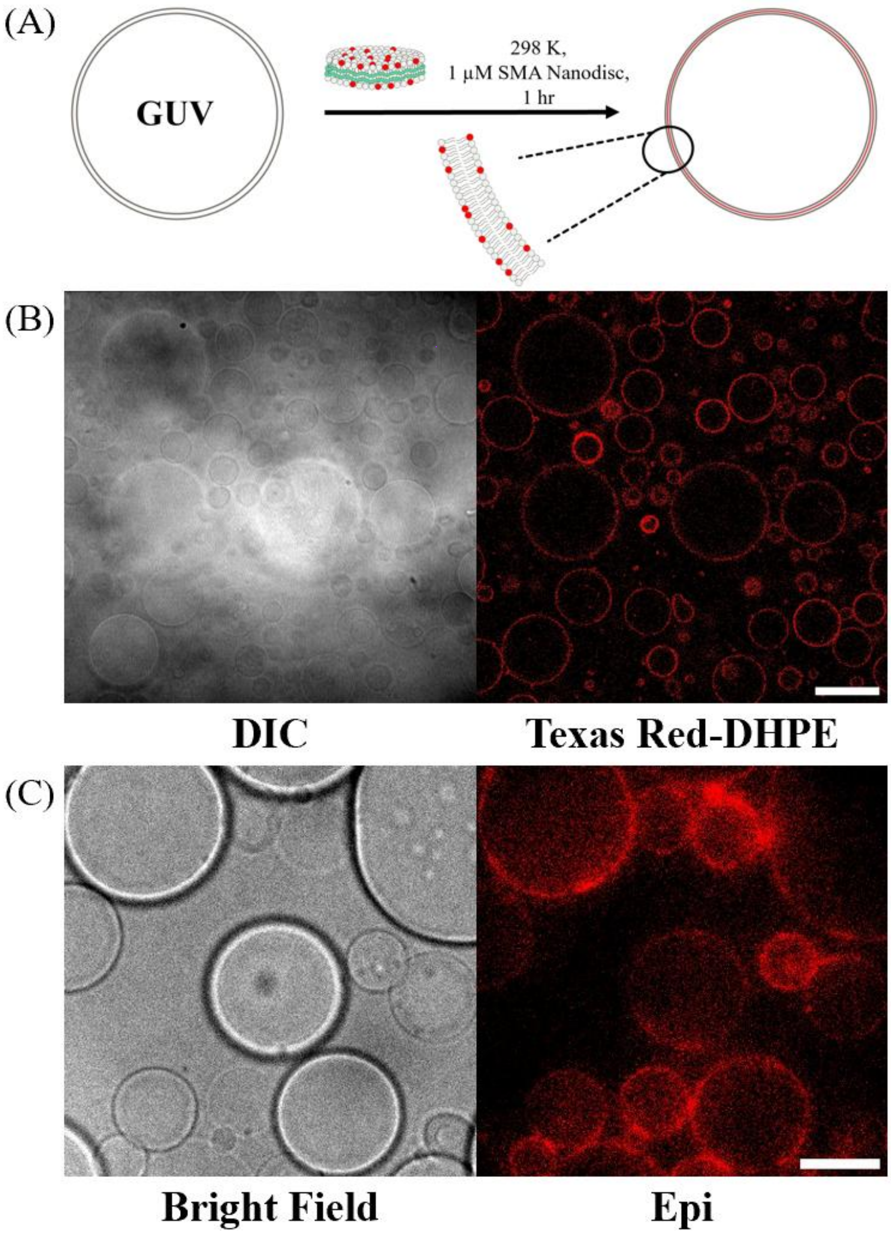


**Figure S18.** SMA nanodisc-mediated delivery of fluorescent lipid cargoes into giant unilamellar vesicle (GUV) model membranes. (A) Schematic illustration of the nanodisc-mediated lipid delivery process. SMA nanodiscs, prepared from small unilamellar vesicles (SUVs) composed of 96 mol% DOPC and 4 mol% Texas Red-DHPE (see Materials and Experimental Methods), are introduced to the GUVs, facilitating the targeted unloading and integration of the fluorescent cargo lipids into the vesicle membrane. (B) Representative differential interference contrast (DIC) and corresponding confocal fluorescence images of the GUVs following a 60 min incubation with the nanodiscs. To enable stable imaging, the GUVs were tethered to the glass substrate via biotin-streptavidin interactions. Confocal optical sectioning reveals a distinct fluorescent ring corresponding to the vesicular cross-section, confirming the successful structural integration of the Texas Red-DHPE lipids into the GUV bilayer. (C) Representative bright-field and epifluorescence images of the nanodisc-treated GUVs. The wide-field imaging confirms robust, uniform Texas Red fluorescence across the entire observed population, further demonstrating the high delivery efficiency of the lipid cargoes into the target membranes.

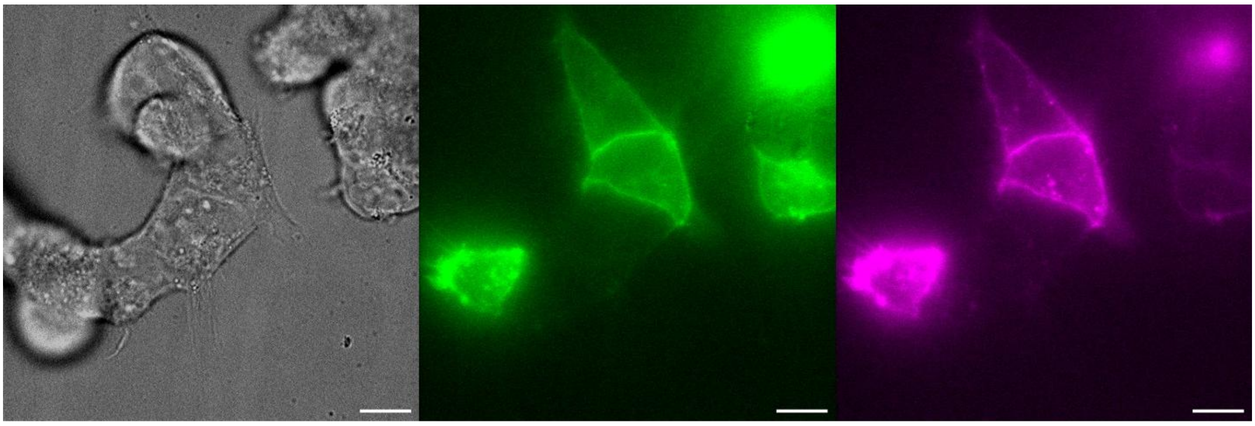

**Bright field** **Epi D2R-mNeonGreen** **Epi AF647-PPHT**

**Figure S19.** Validation of the bioactivity and specificity of the custom-synthesized Alexa Fluor 647-labeled PPHT agonist. Representative bright-field transmission image (left) of a HEK293 cell. The corresponding epifluorescence image (middle) visualizes the expression of D2R, where an mNeonGreen fluorescent tag is genetically fused to the C-terminus of the receptor via a flexible linker. To assess ligand specificity, the cells were incubated with the custom fluorescent agonist (Alexa Fluor 647-labeled PPHT) for 5 min, followed by an imaging buffer wash to remove unbound molecules prior to imaging. The resulting epifluorescence image (right) confirms the bioactivity of the synthesized agonist, demonstrating that robust binding occurs exclusively in cells expressing D2R. Scale bar: 10 μm.

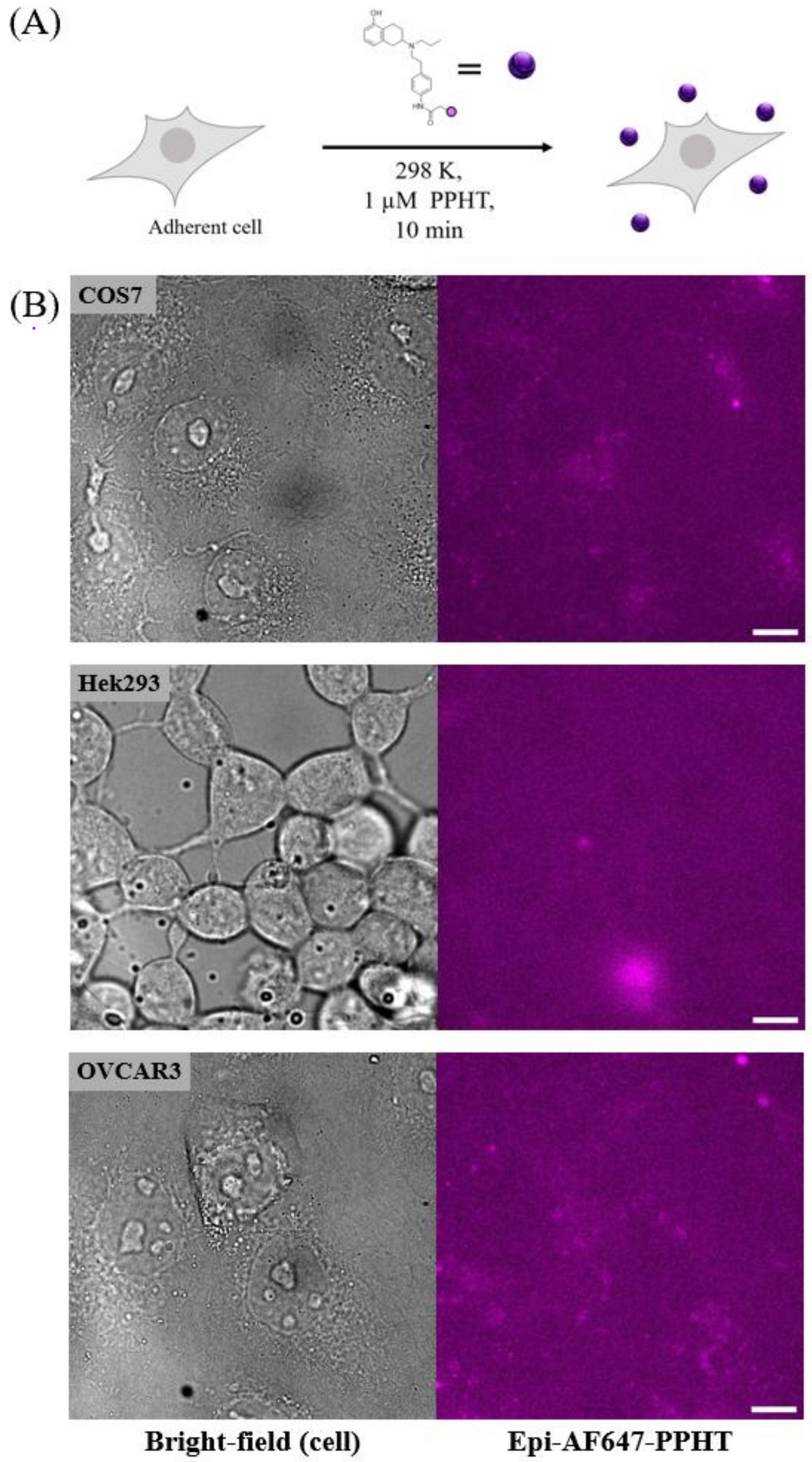


**Figure S20.** Negative control assays confirming the specific binding of the fluorescent agonist to nanodisc-delivered D2R. (A) Schematic illustration of the negative control binding assay. In the absence of SMA nanodisc-mediated delivery of mNeonGreen-fused D2R to the cellular plasma membrane, the custom fluorescent agonist (Alexa Fluor 647-labeled PPHT) lacks its specific receptor target. Consequently, the agonist cannot bind to the plasma membrane, and any unbound molecules are removed via an imaging buffer wash prior to fluorescence visualization. (B) Representative bright-field and corresponding epifluorescence images of various naive mammalian cell lines. Because the D2R channel is absent, the epifluorescence images reveal no apparent agonist binding, displaying only baseline background fluorescence in the Alexa Fluor 647 channel. These results confirm that the robust agonist fluorescence observed in Figure 13 is strictly dependent on the successful nanodisc-mediated integration of D2R.

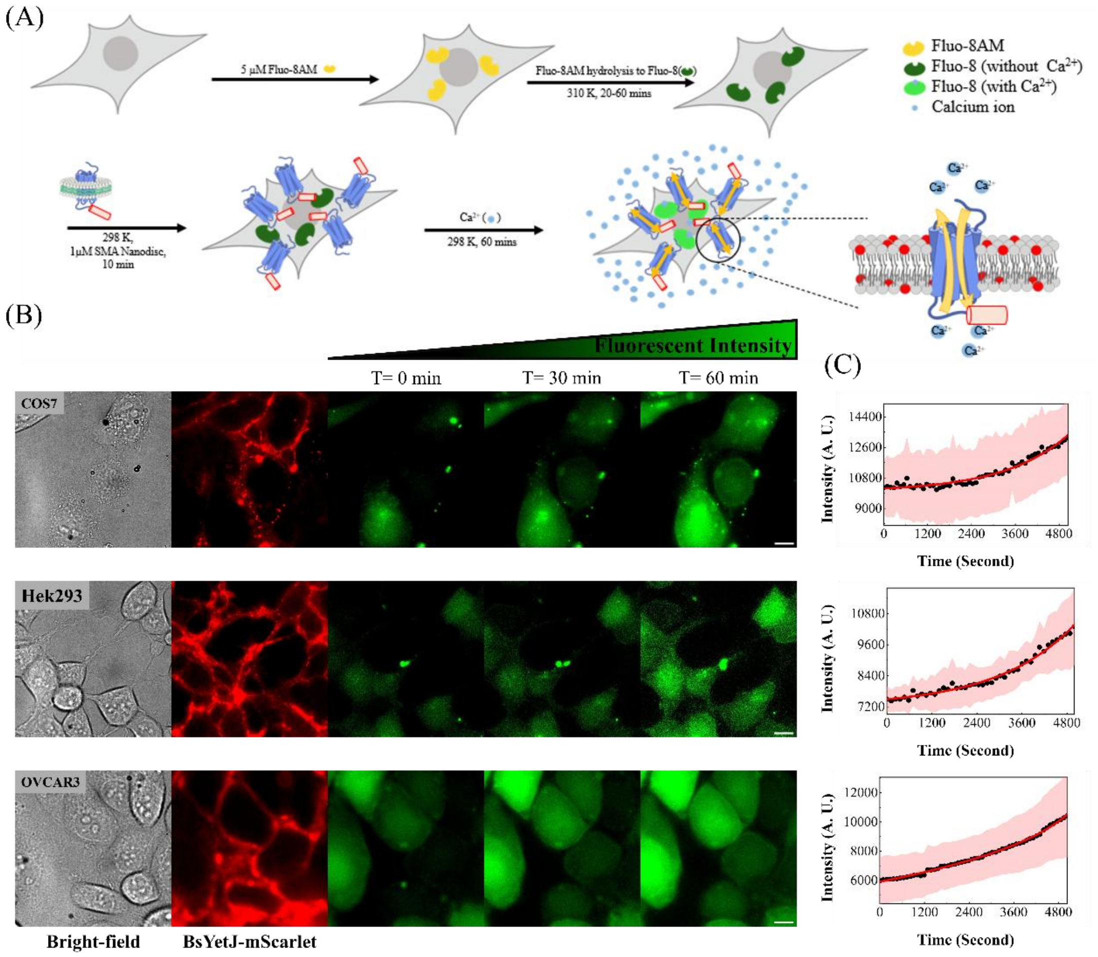


**Figure S21.** Functional validation of intracellular calcium conductance mediated by SMA nanodisc-delivered BsYetJ-mScarlet in different cell lines. (A) Schematic illustration of the intracellular calcium imaging assay. The SMA nanodiscs are loaded with BsYetJ-mScarlet, a construct in which the C-terminus of BsYetJ is genetically fused to the mScarlet fluorescent protein via a flexible linker (GSGSGSENLYFQGGSGSGS). Notably, BsYetJ-mScarlet is directly extracted from the bacterium along with its native membrane lipids (see Materials and Experimental Methods). Following the intracellular loading of the membrane-permeable calcium indicator Fluo-8 AM, full-length BsYetJ-mScarlet is delivered and integrated into the cellular plasma membrane via the SMA nanodiscs. Upon the addition of $Ca^{2+}$ to the extracellular medium, calcium influx through the reconstituted channels is captured via chelation by the active, hydrolyzed Fluo-8 indicator, resulting in a robust increase in fluorescence intensity. (B) Representative bright-field transmission (left) and BsYetJ-mScarlet epifluorescence (second from left) images of the treated cells. The corresponding time-lapse Fluo-8 epifluorescence frames (right) at $t = 0$, 30, and 60 min visually track the calcium influx across the membrane, evidenced by the distinct increase in Fluo-8 fluorescence intensity. (C) Quantitative averaged time-course fluorescence intensity trace. The data demonstrate a distinct, time-dependent increase in cytosolic $Ca^{2+}$ levels mediated by the delivered channels. This confirms that BsYetJ preserves its native conformation and fully active ion-conductive properties following SMA nanodisc-mediated reconstitution.

**The Chemical Synthesis of PPHT Ligand:**

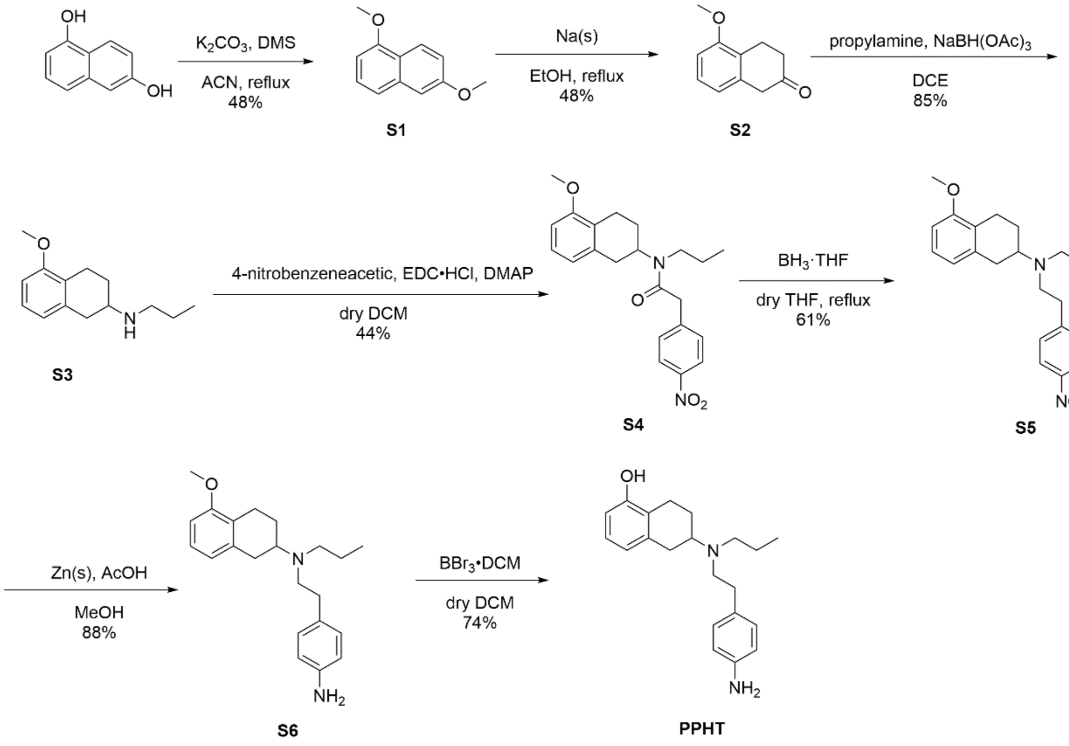


**Scheme S1.** Synthetic route for PPHT ligand.

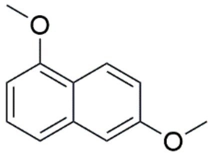

**1,6-dimethoxynaphthalene (S1)**

To a solution of 1,6-dihydroxynaphthalene (1.72 g, 10.7 mmol) and $K_2CO_3$ (1.27 g, 37.7 mmol) in acetonitrile(13.4 mL) was added dimethyl sulfate (4.74 g, 37.6 mmol) dropwise. The reaction mixture was then heated to reflux and stirred overnight. After completion of the reaction, the mixture was cooled to room temperature and concentrated under reduced pressure. The residue was extracted with ethyl acetate and water, and the combined organic layers were dried over anhydrous $Na_2SO_4$ and concentrated under reduced pressure. The crude product was purified by column chromatography (EA / Hexanes = 1 : 5). The product was obtained as a white solid **S1**

(0.94 g, 5.0 mmol, 46%). $^1$H NMR (400 MHz, $CDCl_3$) δ 8.16 (d, J = 8.8 Hz, 1H, ArH), 7.37-7.30 (m, 2H, ArH), 7.13-7.10 (m, 2H, ArH), 6.69 (dd, J = 6.9, 1.7 Hz, 1H, ArH), 3.98 (s, 3H, $OCH_3$), 3.92 (s, 3H, $OCH_3$). The proton NMR spectrum was consistent with literature data[1].

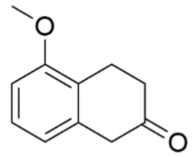

**5-Methoxy-2-tetralone (S2)**

To a solution of compound **S1** (449.0 mg, 2.39 mmol) in EtOH (16 mL) was heated to 50 °C, and Na (614.4 mg, 26.7 mmol) was added in small portions. After the sodium had completely dissolved, the reaction mixture was heated to reflux and stirred for 6 h. The mixture was then cooled to room temperature and acidified to pH 1 at 0 °C with 12 N HCl (aq). The resulting mixture was subsequently heated to reflux and stirred for an additional 1 h before being cooled to room temperature. Water was added, and the mixture was concentrated under reduced pressure. The residue was extracted with DCM, and the organic layers were combined, dried over $Na_2SO_4$ (s), and concentrated under reduced pressure. The crude product was purified by column chromatography (EA / Hexanes = 1 : 5). The product was obtained as a yellow oil **S2** (190.2 mg, 1.08 mmol, 45%) $^1$H NMR (400 MHz, $CDCl_3$) δ 7.18 (t, J = 8.0 Hz, 1H, ArH), 6.78 (d, J = 8.0 Hz, 1H, ArH), 6.74 (d, J = 8.0 Hz, 1H, ArH), 3.85 (s, 3H, $OCH_3$), 3.57 (s, 2H, Ar$CH_2$CO), 3.09 (t, J = 6.8 Hz, 2H, Ar$CH_2$$CH_2$CO) , 2.52 (t, J = 6.8 Hz, 2H, Ar$CH_2$$CH_2$CO). The proton NMR spectrum was consistent with literature data[1].

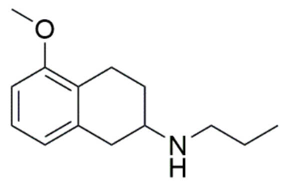

**2-(N-propylamino)-5-methoxytetraline (S3)**

To a solution of compound **S2** (132.3 mg, 0.75 mmol) in 1,2-dichloroethane (8 mL) under an argon atmosphere, n-propylamine (92 μL, 1.11 mmol) was added dropwise at room temperature. The reaction mixture was stirred for 1 h, after which $NaBH(OAc)_3$ was added, and the mixture was stirred overnight at room temperature. Saturated $NaHCO_3$ (aq) and ethyl acetate were added to the reaction mixture, and the layers were separated. The organic layer was collected, and the aqueous layer was extracted with ethyl acetate. The combined organic layers were dried over $Na_2SO_4$ (s) and concentrated under reduced pressure. The crude product was purified by column chromatography (EA / MeOH / TEA = 92 : 7 : 1). The product was obtained as a yellow oil **S3** (136.6 mg, 0.63 mmol, 85%) $^1$H NMR (400 MHz, $CDCl_3$) δ 7.09 (t, J = 8.0 Hz, 1H, ArH), 6.70 (d, J = 8.0 Hz, 1H, ArH), 6.60 (d, J = 8.0 Hz, 1H, ArH), 3.81(s, 3H, $OCH_3$), 3.0 (dd, J = 16.0, 3.4 Hz, 1H, Ar$\mathit{CH_2}$CHNH), 2.93-2.86 (m, 2H, Ar$\mathit{CH_2}$$CH_2$CHNH, Ar$\mathit{CH_2}$CHNH), 2.68 (t, J = 7.34, 2H, NH$\mathit{CH_2}$$CH_2CH_3$), 2.61-2.52 (m, 2H, Ar$\mathit{CH_2}$$CH_2$CHNH, Ar$CH_2$*CH*NH), 2.09-2.05 (m, 1H, Ar$CH_2$$\mathit{CH_2}$CHNH), 1.60-1.51 (m, 3H, Ar$CH_2$$\mathit{CH_2}$CHNH, NH$CH_2$$\mathit{CH_2}$$CH_3$), 0.94 (t, J = 7.4 Hz, NH$CH_2CH_2$$\mathit{CH_3}$). The proton NMR spectrum was consistent with literature data[1].

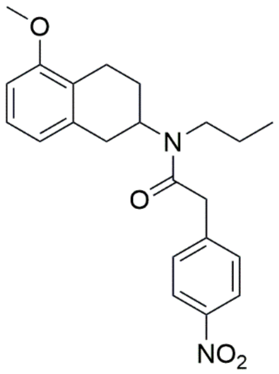

***N*-(5-methoxy-2-tetralinyl)-2-(4-nitrophenyl)-*N*-propylacetamide (S4)**

To a solution of 4-nitrobenzeneacetic acid (180.5 mg, 1.0 mmol) in dry dichloromethane (10 mL) under an argon atmosphere, EDC・HCl (250.7 mg, 1.31 mmol), DMAP (19.0 mg, 0.16 mmol), and compound **S3** (136.6 mg, 0.62 mmol) were added sequentially. The reaction mixture was stirred at room temperature overnight then acidified to pH < 1 with 1 N HCl, and the layers were separated. The aqueous layer was basified with saturated $NaHCO_3$ (aq) and extracted with dichloromethane. The combined organic layers were dried over $Na_2SO_4$ (s) and concentrated under reduced pressure. The crude product was purified by column chromatography (EA / Hexanes = 1 : 1). The product was obtained as a dark brown oil **S4** (55.4 mg, 0.15 mmol, 23%). 1H NMR (400 MHz, $CDCl_3$) : δ 8.20 (d, *J* = 8.7 Hz, 2H, ArH), 8.17 (d, *J* = 8.7 Hz, 2H, ArH), 7.46 (d, *J* = 8.7 Hz, 2H, ArH), 7.41 (d, *J* = 8.7 Hz, 2H, ArH), 7.12 (t, *J* = 8.0 Hz, 1H, ArH), 7.08 (t, *J* = 8.0 Hz, 1H, ArH), 6.71-6.63 (m, 4H, ArH), 4.63-4.51 (m, 1H, $ArCH_2C\mathit{H}N$), 4.03-3.91(m, 1H, $ArCH_2C\mathit{H}N$), 3.85 (d, J = 2.4 Hz, $ArC\mathit{H_2}CO$), 3.82 (d, J = 3.4 Hz, $ArC\mathit{H_2}CO$), 3.81 (s, $OC\mathit{H_3}$), 3.80 (s, $OC\mathit{H_3}$), 3.30-3.14 (m, 4H, $Ar\mathit{CH_2}CHN$), 3.04-2.96 (m, 4H, $Ar\mathit{CH_2}CH_2CHN$), 2.87-2.81 (m, 1H, $ArCH_2\mathit{CH_2}CHN$), 2.72-2.66 (m, 1H, $ArCH_2\mathit{CH_2}CHN$), 2.64-2.57 (m, 1H, $ArCH_2\mathit{CH_2}CHN$), 2.49-2.40 (m, 1H, $ArCH_2\mathit{CH_2}CHN$), 2.00-1.89 (m, 2H, $N\mathit{CH_2}CH_2CH_3$), 1.86-1.79 (m, 2H, $N\mathit{CH_2}CH_2CH_3$), 1.72-1.62 (m, 4H, N, $NCH_2\mathit{CH_2}CH_3$), 0.95(t, J = 7.4 Hz, 3H,

$NCH_2CH_2CH_3$), 0.90 (t, J = 7.4 Hz, 3H, $NCH_2CH_2CH_3$). The proton NMR spectrum was consistent with literature data[1].

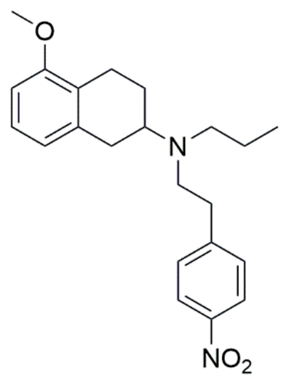


**2-[*N*-(4-nitrophenethyl)-*N*-propylamino]-5-methoxytetraline (S5)**

To a solution of compound **S4** (100.0 mg, 0.261 mmol) in dry THF (0.35 mL) under an argon atmosphere at 0 °C, 1M $BH_3$ · THF (1 mL) was added dropwise. The reaction mixture was stirred at 0 °C for 5 min and then heated to reflux overnight. After cooling to 0 °C, the reaction mixture was carefully acidified to pH 2 with 6 N HCl to decompose the excess $BH_3$. The mixture was then basified to pH 13 with 10% aqueous NaOH and extracted with dichloromethane. The combined organic layers were dried over $Na_2SO_4$ (s) and concentrated under reduced pressure. The crude product was purified by column chromatography (EA / Hexanes = 1 : 3). The product was obtained as a yellow oil **S5** (50.7 mg, 0.138 mmol, 52%). $^1H$ NMR (400 MHz, $CDCl_3$) : δ 8.14 (d, $J$ = 8.8 Hz, 2H, ArH), 7.35 (d, $J$ = 8.8 Hz, 2H, ArH), 7.08 (t, $J$ = 7.8 Hz, 1H, ArH), 6.68 (d, $J$ = 7.8 Hz, 1H, ArH), 6.65 (d, $J$ = 8.0 Hz, 1H, ArH), 3.80 (s, 3H, $OCH_3$), 3.00-2.94 (m, 2H, $ArCH_2CH_2CHN$, $ArCH_2CHN$), 2.84-2.66 (m, 5H, $ArCH_2CHN$, $NCH_2CH_2Ar$), 2.54-2.46 (m, 3H, $ArCH_2CH_2CHN$, $NCH_2CH_2CH_3$), 2.00-1.97 (m, 1H, $ArCH_2CHN$), 1.58-1.51 (m, 2H, $ArCH_2CH_2CHN$), 1.45-1.40 (m, 2H, $NCH_2CH_2CH_3$), 0.86 (t, $J$ = 7.3 Hz, 3H, $NCH_2CH_2CH_3$). The proton NMR spectrum was consistent with literature data[1].

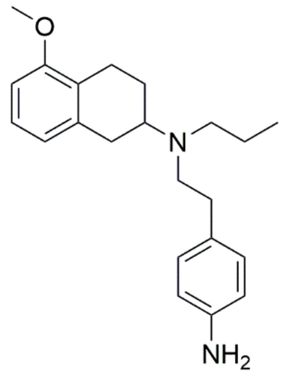

**2-[*N*-(4-aminophenethyl)-N-propylamino]-5-methoxytetraline (S6)**

To a solution of compound **S5** (50.7 mg, 0.138 mmol) in methanol (1.6 mL) was added zinc powder (31.6 mg, 0.483 mmol) and acetic acid (0.54 mL), then stirred for 5 hours at room temperature under an argon atmosphere. The reaction mixture was concentrated under reduced pressure, then adjusted to pH = 2 by 1 N HCl (aq), washed with ethyl acetate several times. The resulting solution was basified with 10 % NaOH (aq) until pH = 13, then extracted with dichloromethane several times. The combined organic phase was dried over $Na_2SO_4$ (s) and concentrated under reduced pressure to give dark brown oil **S6** (41.0 mg, 0.121 mmol, 88 %). The crude product was used for the next step without further purification. $^1$H NMR (400 MHz, $CDCl_3$) : δ 7.08 (t, J = 7.8 Hz, 1H, ArH), 7.00-6.97 (m, 2H, ArH), 6.70 (d, J = 7.5 Hz, 1H, ArH), 6.66 (s, 1H, ArH), 6.64-6.61 (m, 2H, ArH), 3.80 (s, 3H, OCH3), 3.54 (br s, 2H, $NH_2$), 3.00-2.93 (m, 2H, ArC*H*$_2$CH$_2$CHN, ArC*H*$_2$CHN), 2.93-2.80 (m, 1H, ArC*H*$_2$CHN), 2.80-2.69 (m, 3H, ArC*H*$_2$CH$_2$CHN, NC*H*$_2$CH$_2$Ar), 2.69-2.63 (m, 2H, NCH$_2$C*H*$_2$Ar), 2.56-2.46 (m, 3H, ArCH$_2$C*H*N, NC*H*$_2$CH$_2$CH$_3$), 2.10-2.00 (m, 1H, ArCH$_2$C*H*$_2$CHN), 1.54-1.46 (m, 3H, ArCH$_2$C*H*$_2$CHN, NCH$_2$C*H*$_2$CH$_3$), 0.90 (t, J = 7.3 Hz, 3H, NCH$_2$CH$_2$C*H*$_3$). The proton NMR spectrum was consistent with literature data[1] .

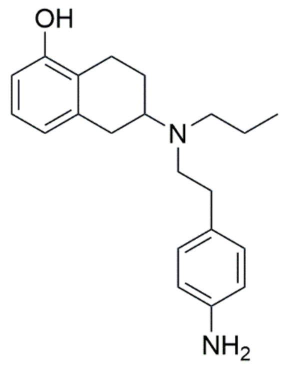

**2-[*N*-(4-aminophenethyl)-*N*-propylamino]-5-hydroxytetraline (PPHT)**

To a cold (-78 °C) solution of compound **S6** (41 mg, 0.121 mmol) in dry dichloromethane (2.0 mL) was added a 1M $BBr_3$ solution in dichloromethane (0.24 mL, 0.24 mmol) under an argon atmosphere. The reaction mixture was stirred for 4 h at -78°C and was then allowed to warm to rt, where it was stirred for overnight. The final mixture was extracted with brine. The combined aqueous layers were basified to pH = 8 with 10 % NaOH (aq) and subsequently extracted with dichloromethane several times. The combined organic phase was dried over $Na_2SO_4$ (s) and concentrated under reduced pressure. The residue was purified by column chromatography (MeOH /DCM = 1 : 10) to give yellow oil **PPHT** (29.0 mg, 0.090 mmol, 74 %). $^1$H NMR (400 MHz, $CDCl_3$) : δ 7.00-6.96 (m, 3H, ArH), 6.69 (d, *J* = 7.5 Hz, 1H, ArH), 6.63 (d, *J* = 8.3 Hz, 2H, ArH), 6.59 (d, *J* = 7.8 Hz, 1H, ArH), 3.56 (br s, 2H, $NH_2$), 3.00-2.77 (m, 3H, ArC$H_2$CH$_2$CHN, ArC$H_2$CHN), 2.77-2.67 (m, 3H, ArC$H_2$CH$_2$CHN, NC$H_2$CH$_2$Ar), 2.67-2.60 (m, 2H, NCH$_2$C$H_2$Ar), 2.60-2.50 (m, 3H, ArCH$_2$C*H*N, NC$H_2$CH$_2$CH$_3$), 2.12-2.04 (m, 1H, ArCH$_2$C$H_2$CHN), 1.50-1.43 (m, 3H, ArCH$_2$C$H_2$CHN, NCH$_2$C$H_2$CH$_3$), 0.90 (t, J = 7.2 Hz, 3H, NCH$_2$CH$_2$C$H_3$). The NMR spectrum was consistent with literature data.[1] .

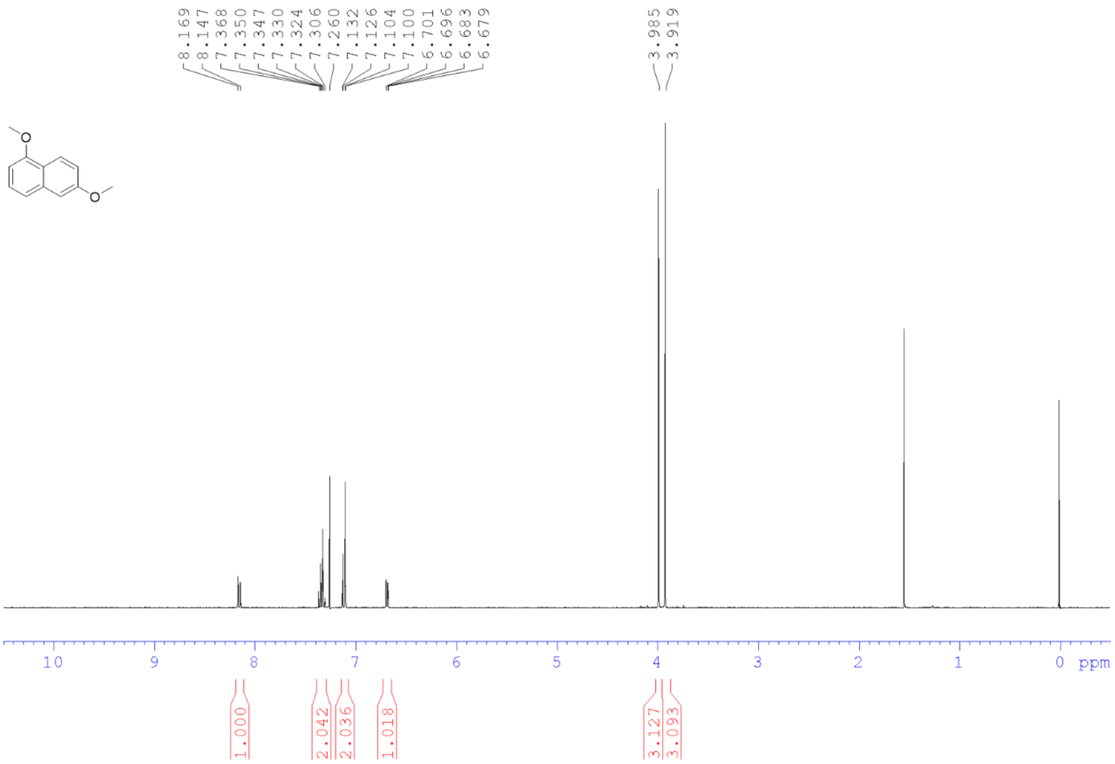


**Figure SOOO.** $^{1}$H NMR spectrum of **S1** in $CDCl_3$ (400 MHz).

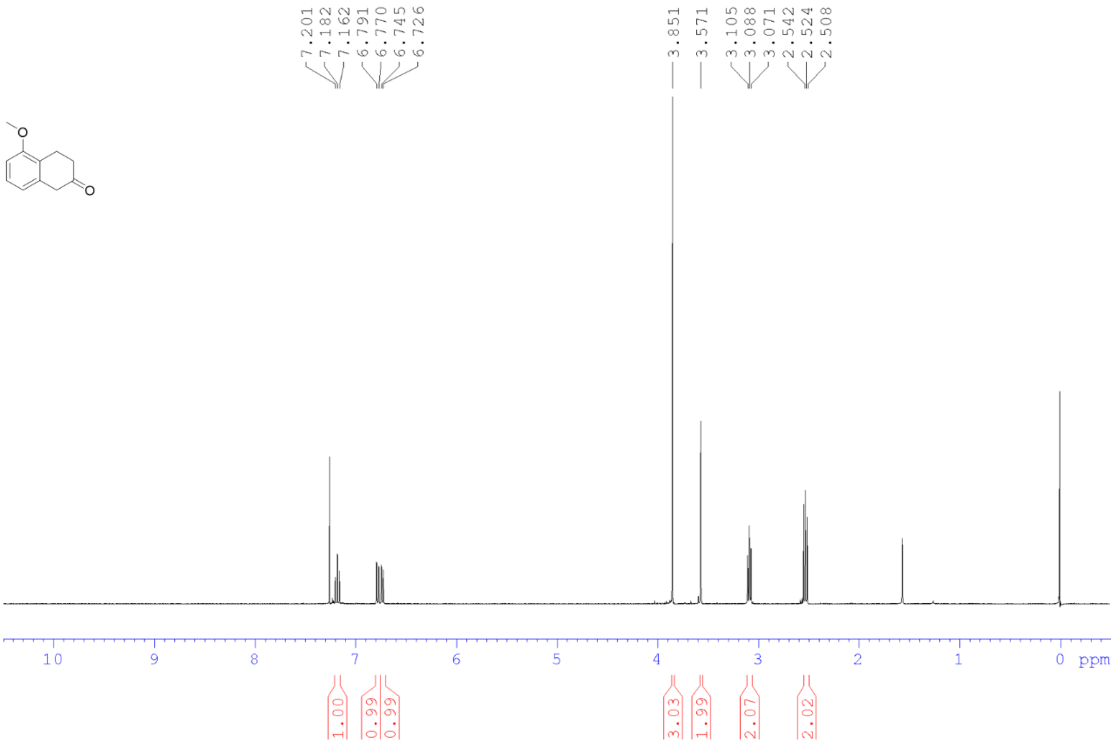


**Figure SOOO.** $^{1}$H NMR spectrum of **S2** in $CDCl_3$ (400 MHz).

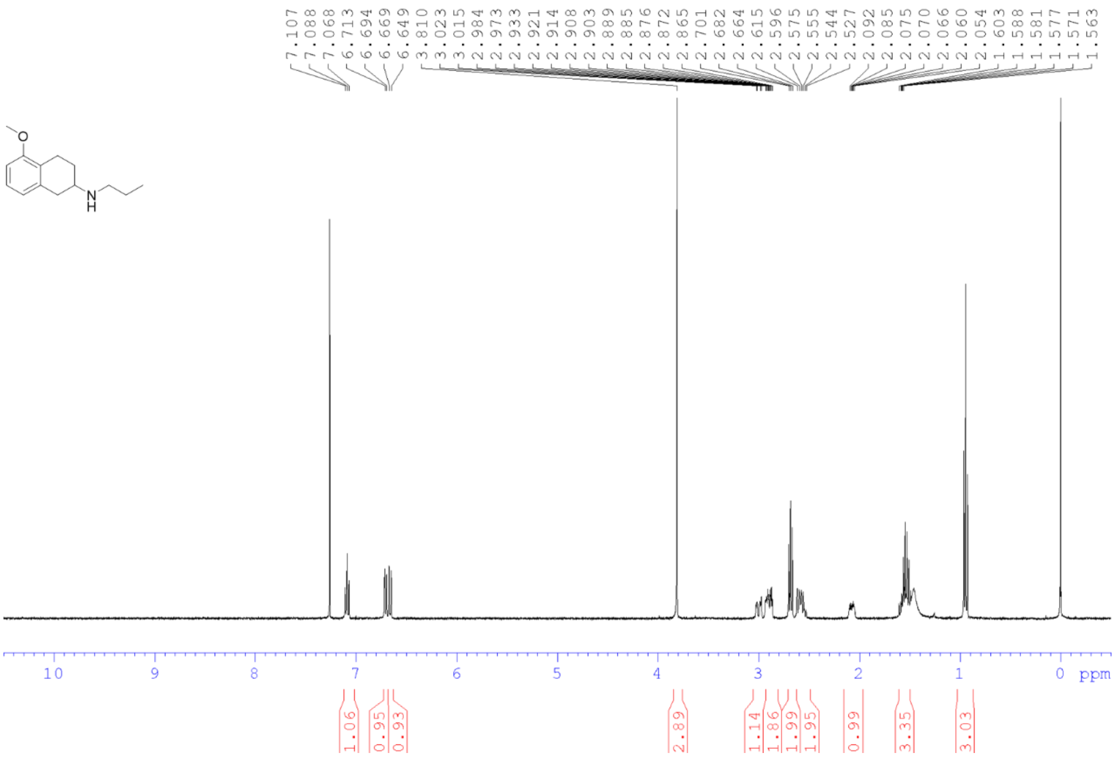

**Figure SOOO.** $^{1}$H NMR spectrum of **S3** in $CDCl_3$ (400 MHz).

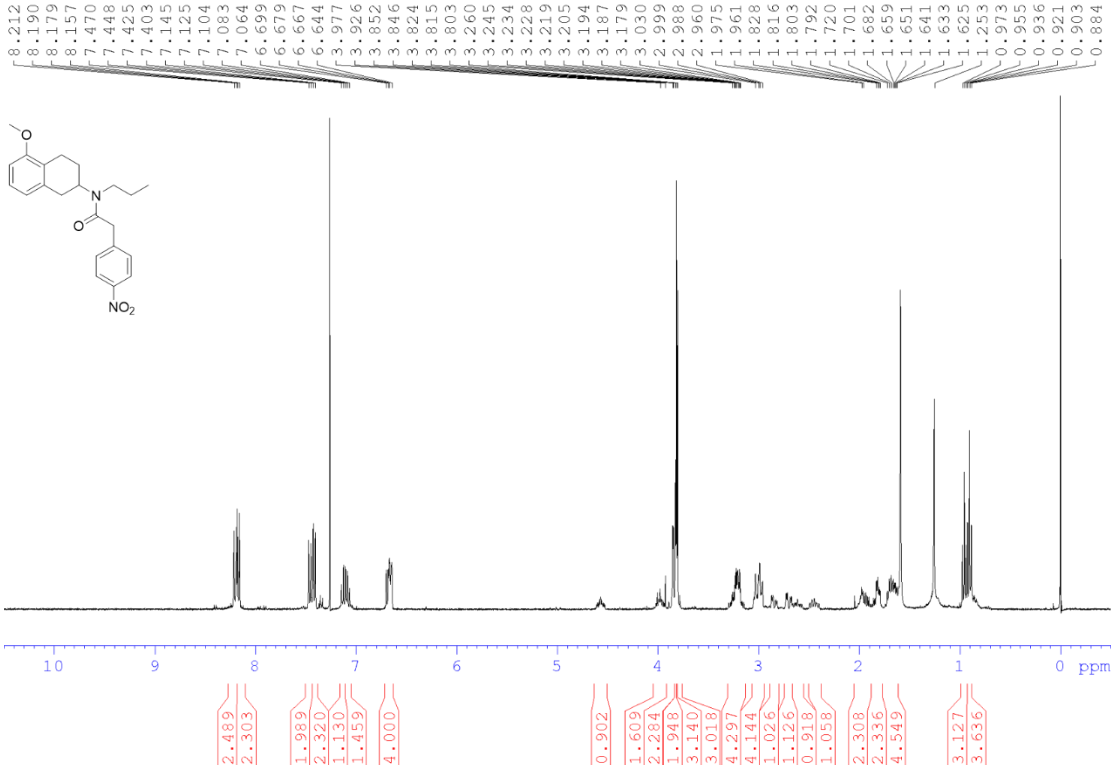

**Figure SOOO.** $^{1}$H NMR spectrum of **S4** in $CDCl_3$ (400 MHz).

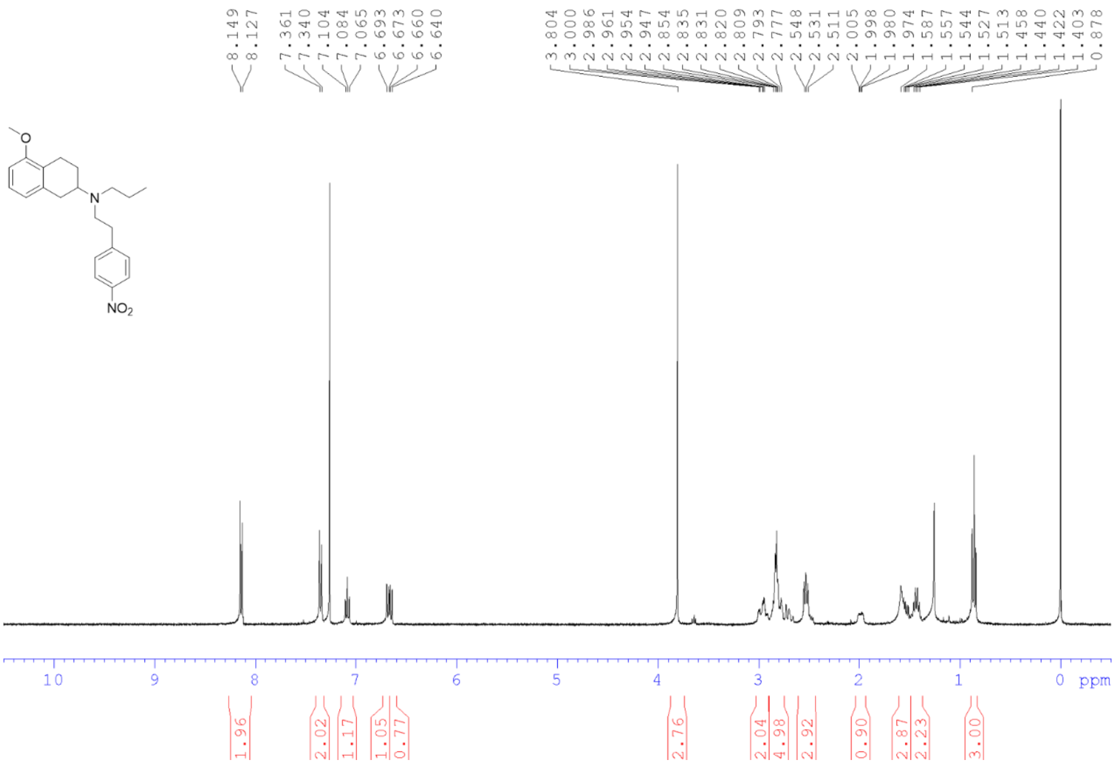

**Figure SOOO.** [1]H NMR spectrum of **S5** in $CDCl_3$ (400 MHz).

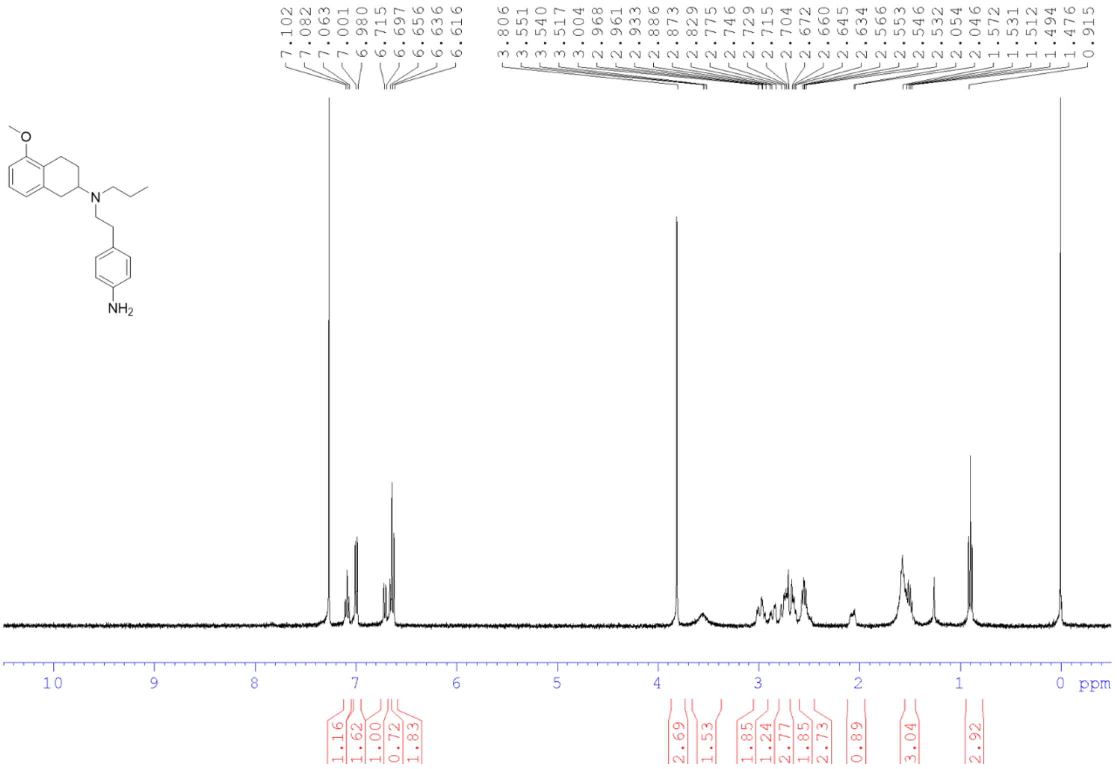

**Figure SOOO.** [1]H NMR spectrum of **S6** in $CDCl_3$ (400 MHz).

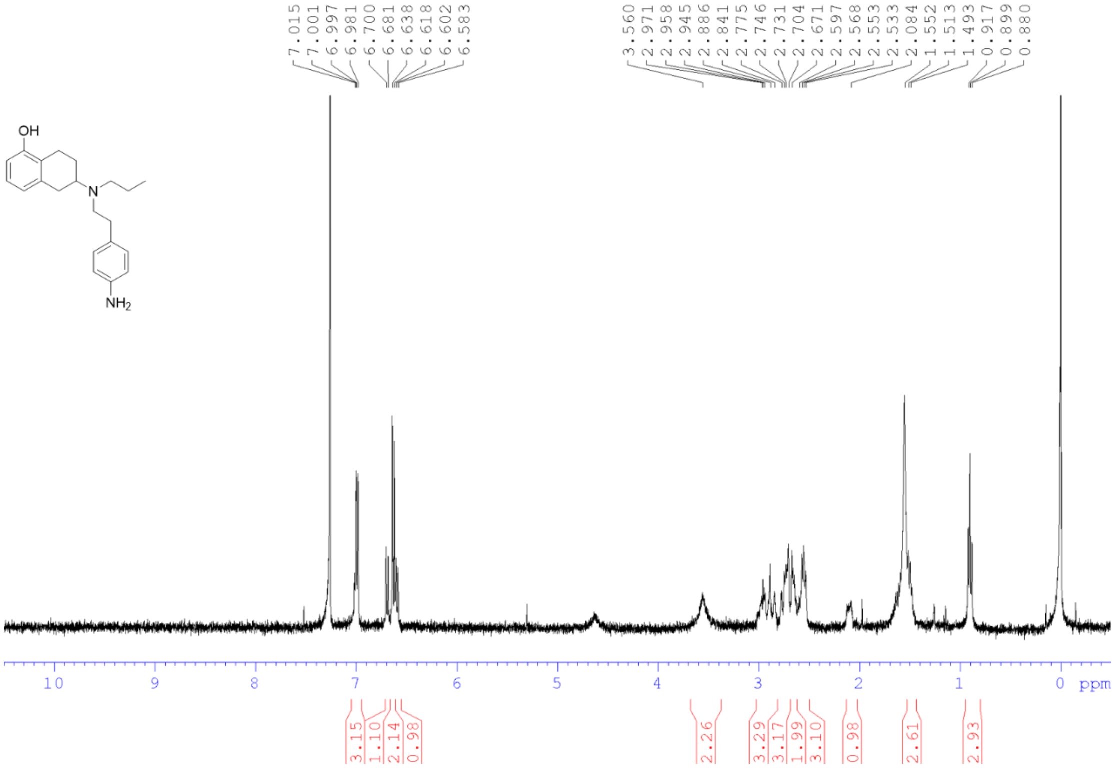


**Figure SOOO.** $^1$H NMR spectrum of **PPHT** in $CDCl_3$ (400 MHz).

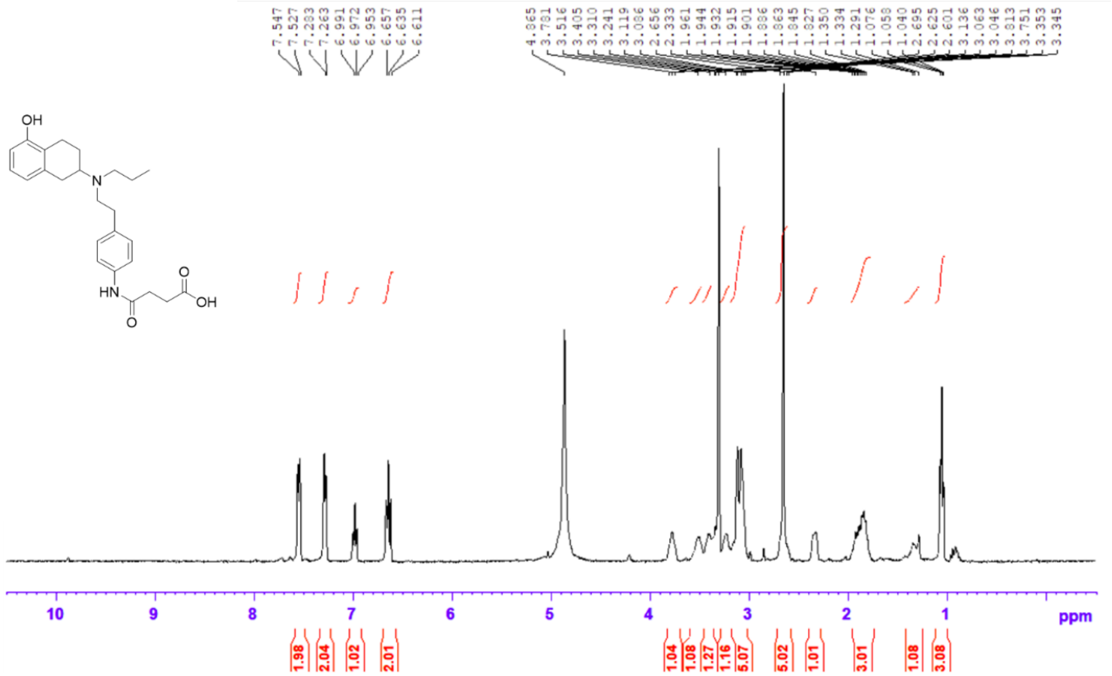


**Figure SOOO.** $^1$H NMR spectrum of **1** in $CDCl_3$ (400 MHz).

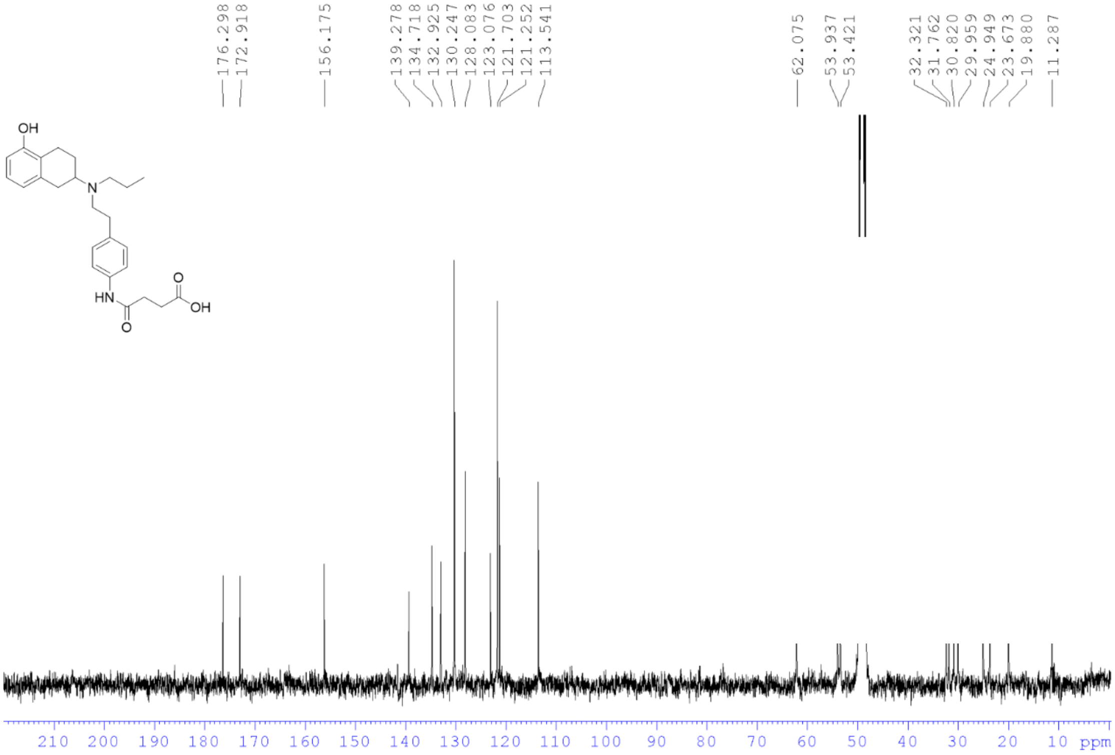

**Figure SOOO.** $^{13}C$ NMR spectrum of **1** in $CDCl_3$ (100.6 MHz).

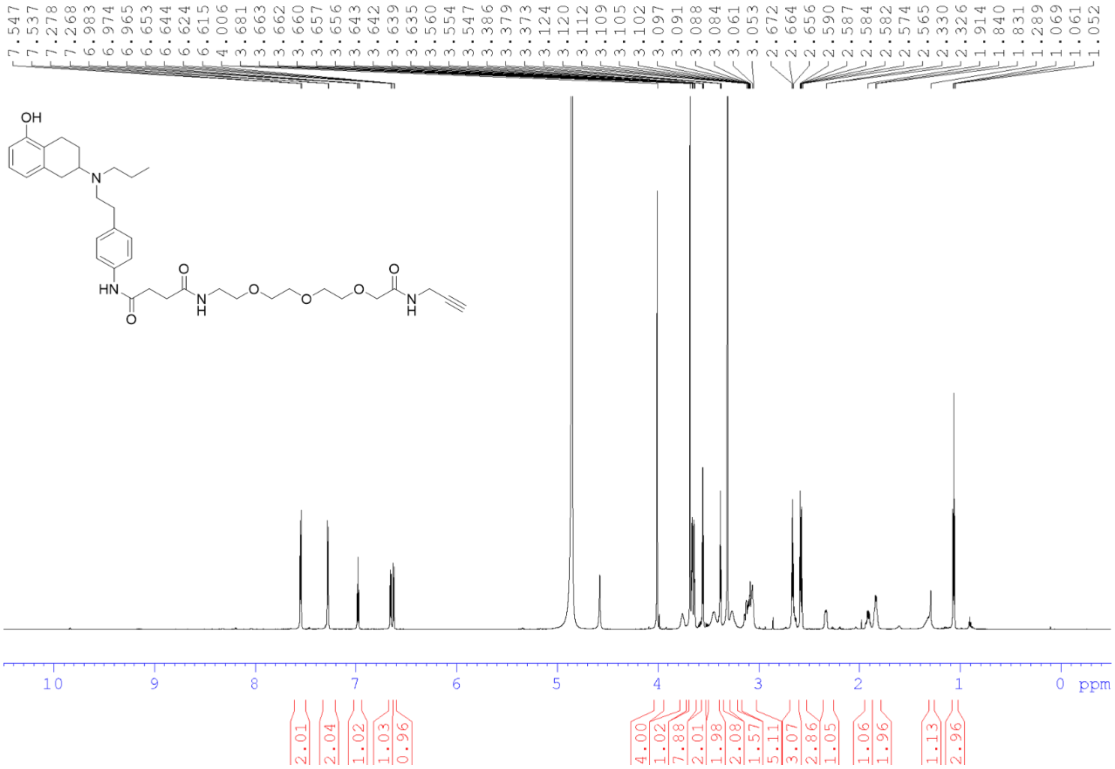

**Figure SOOO.** $^{1}H$ NMR spectrum of **2** in $CDCl_3$ (850 MHz).

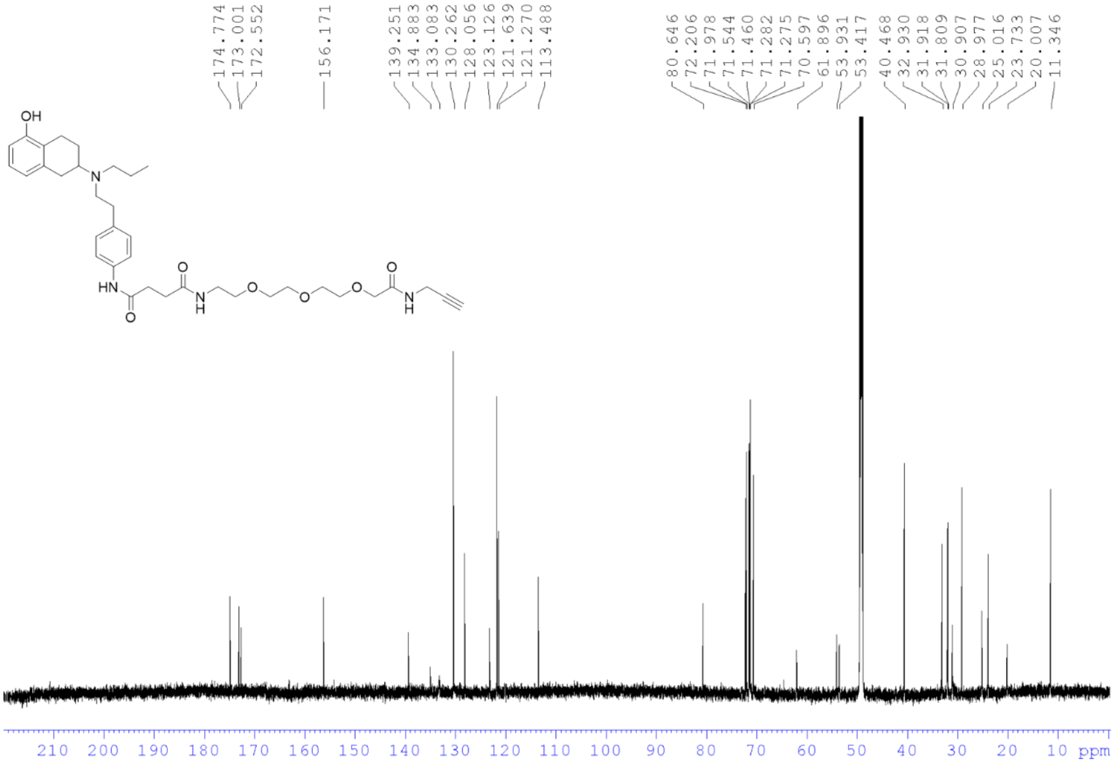

**Figure SOOO.** $^{13}$C NMR spectrum of **2** in $CDCl_3$ (212.5 MHz).

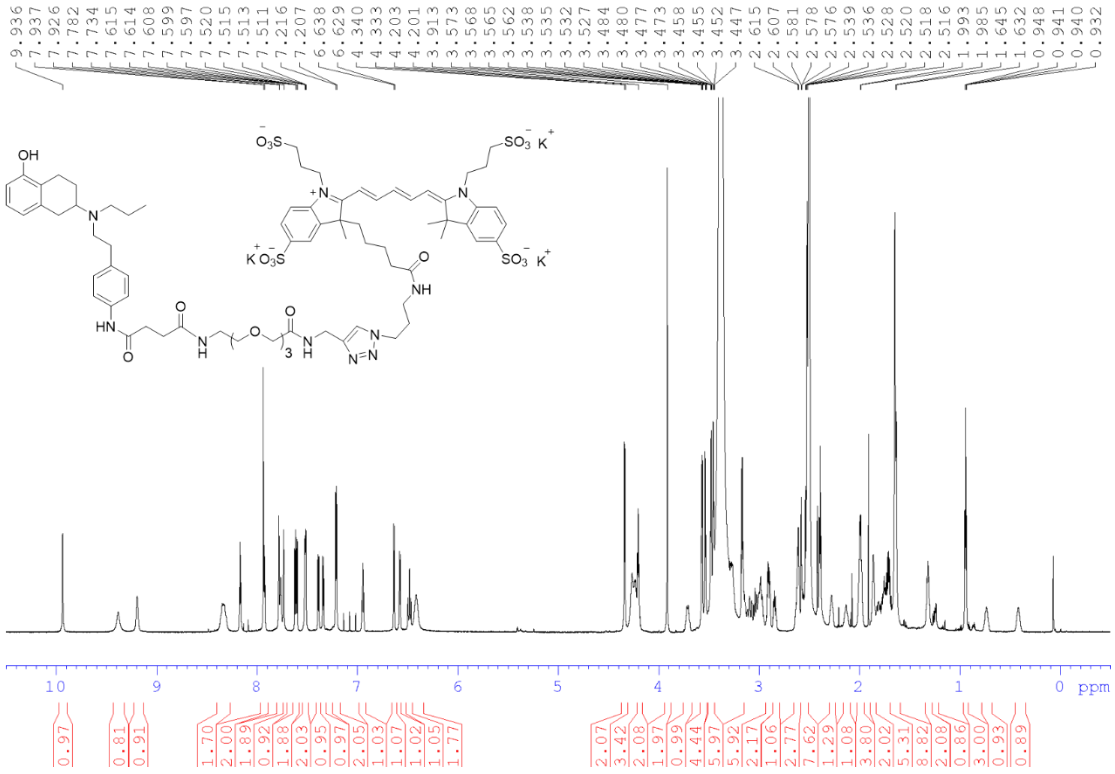

**Figure SOOO.** $^{1}$H NMR spectrum of **AF647-labeled PPHT** in $d^6$-DMSO (850 MHz).

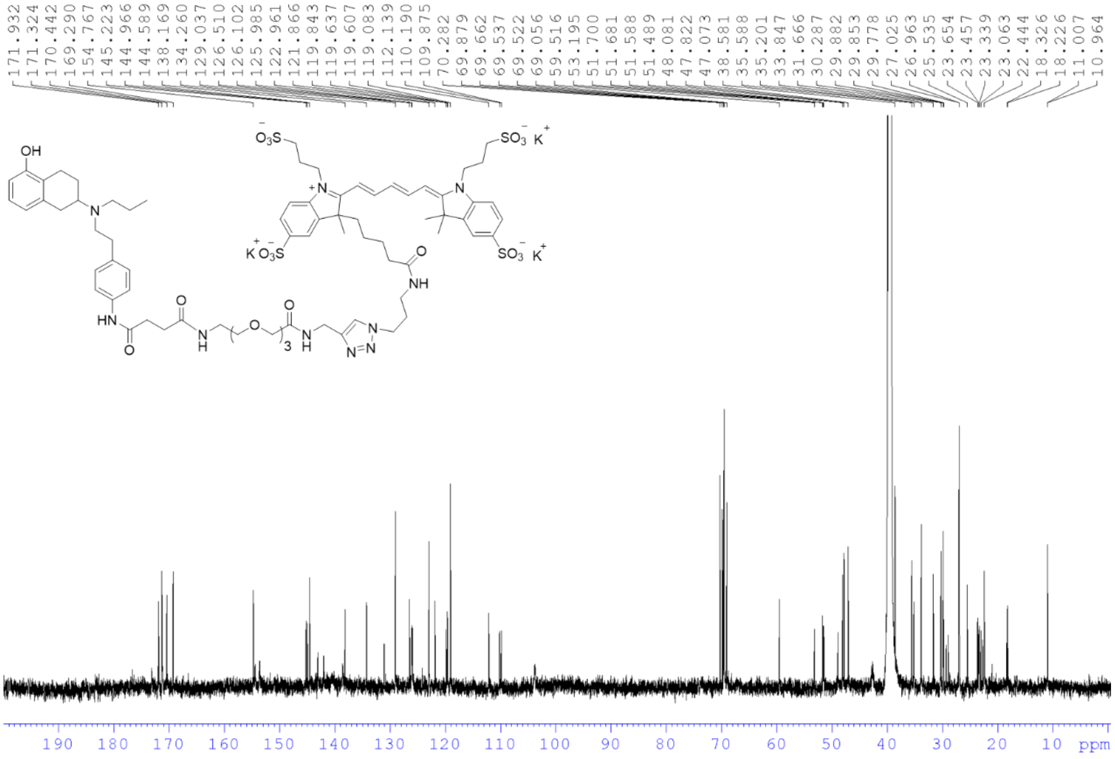

**Figure SOOO.** $^{13}$C NMR spectrum of **AF647-labeled PPHT** in d$^6$-DMSO (212.5 MHz).

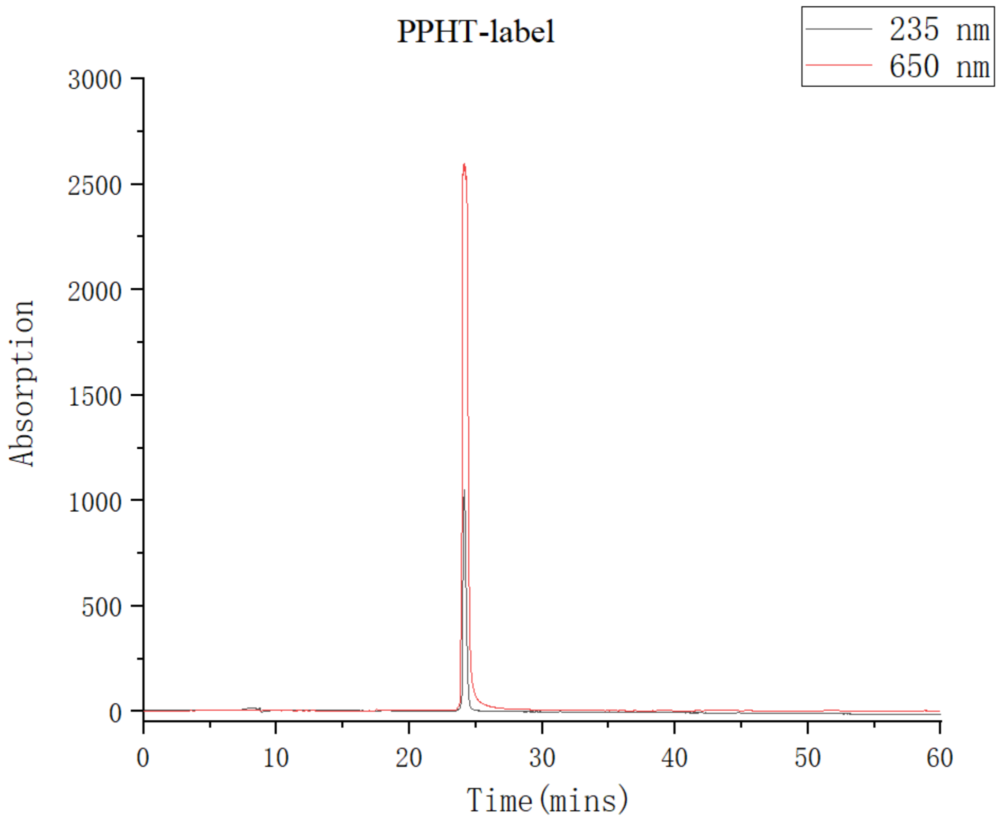


**Figure SOOO.** Analytical HPLC elution profile of **AF647-labeled PPHT**. The analysis was achieved using a linear gradient of 5–90% acetonitrile containing 0.1% trifluoroacetic acid in $H_2O$ over 60 minutes at a flow rate of 0.5 mL/min. The retention time (*t*R) of the major peak is observed at 23.8 min.
